# Linkage analysis and haplotype phasing in experimental autopolyploid populations with high ploidy level using hidden Markov models

**DOI:** 10.1101/415232

**Authors:** Marcelo Mollinari, Antonio Augusto Franco Garcia

## Abstract

Modern SNP genotyping technologies allow to measure the relative abundance of different alleles for a given locus and consequently to estimate their allele dosage, opening a new road for genetic studies in autopolyploids. Despite advances in genetic linkage analysis in autotetraploids, there is a lack of statistical models to perform linkage analysis in organisms with higher ploidy levels. In this paper, we present a statistical method to estimate recombination fractions and infer linkage phases in full-sib populations of autopolyploid species with even ploidy levels in a sequence of SNP markers using hidden Markov models. Our method uses efficient two-point procedures to reduce the search space for the best linkage phase configuration and reestimate the final parameters using the maximum-likelihood of the Markov chain. To evaluate the method, and demonstrate its properties, we rely on simulations of autotetraploid, autohexaploid and autooctaploid populations and on a real tetraploid potato data set. The results demonstrate the reliability of our approach, including situations with complex linkage phase scenarios in hexaploid and octaploid populations.

**Author summary:** In this paper, we present a complete multilocus solution based on hidden Markov models to estimate recombination fractions and infer the linkage phase configuration in full-sib mapping populations with even ploidy levels under random chromosome segregation. We also present an efficient pairwise loci analysis to be used in cases were the multilocus analysis becomes compute-intensive.

## Introduction

Polyploids are organisms with more than two sets of chromosomes. They are very important in agriculture and play a fundamental role in evolutionary processes, such as differentiation of species [48]. The number of sets of chromosomes in an organism is called *ploidy level.* These multiple chromosome sets can originate from the combination of genomes from different, but related species, or from duplicated genomes from the same species [4, 10]. In the first scenario, they are called *allopolyploids*; in the second, *autopolyploids.* Polyploid organisms are also characterized according to their pattern of inheritance. In general, allopolyploids exhibit diploid-like (or *disomic*) segregation, since homologous chromosomes tend to form bivalents within each sub-genome. Autopolyploids, however, have more than two homologous chromosomes per homology group, forming either random bivalents or multivalents during the meiosis resulting in *polysomic segregation* [39, 49, 52]. Since the molecular mechanics of polyploid organisms are quite complex, this dichotomy is often broken, and polyploids can display intermediate modes of inheritance [39, 40]. Throughout this paper, the term autopolyploid (or autotetraploid, autohexaploid, etc.) will refer to polyploid organisms that exhibit polysomic segregation.

Despite all advances in genetic studies in autotetraploids [13,14,20,21,27,30,32,34,61,64,65], there is still a shortage of statistical methods to address organisms with higher ploidy levels, such as sweet potato [1, 24, 47], sugarcane [16, 59], some ornamental flowers and forage crops (reviewed in [50]). In this work, we denote as *high-level autopolyploids* those autopolyploid organisms with ploidy level greater than four. A fundamental class of statistical methods that are lagged behind in high-level autopolyploid studies is the construction of genetic maps. A reliable genetic map is a crucial step in quantitative trait loci (QTL) analysis, as well as the assembly of reference genomes and the study of evolutionary processes [28, 29, 31]. Although understanding the concept of genetic mapping is rather easy, the construction of such maps in high-level autopolyploids is challenging. Even under bivalent pairing, there is a large number of possible configurations during the meiosis, and this number gets exponentially larger as the ploidy level increases. Denoting *m* as the ploidy level, it is possible to find up to *m* different alleles for a locus in one individual. Furthermore, if some of those alleles are not distinguishable, it is necessary to consider the number of copies of each different allelic form, also known as *dosage.*

The construction of a genetic map in a full-sib population can be summarized in five basic steps: *i*) estimation of pairwise recombination fractions and associated statistical tests; *ii*) separation of markers into linkage groups; *iii*) order of markers within each linkage group using an optimization technique; *iv*) parental phasing, recombination fraction update and likelihood computation and *v*) if the order is optimal, the map is complete, otherwise, return to step *iii*. Historically, genetic maps in high-level autopolyploids have been constructed using only alleles present in one homologous chromosome, called *single-dose* or *simplex* markers [51, 60]. In a full-sib population, these markers segregate in a 1:1 ratio (if they are present only in one parent), or in a 3:1 ratio (if present in both parents, also called *double simplex*). Given this level of simplification, it is possible to use the five-step procedure coupled with a standard software suitable for diploid populations. Nevertheless, it is well accepted that the use of single-dose markers imposes limitations on the construction of adequate genetic maps. These approaches sub-sample the genome [16, 21], which precludes further consideration of multiallelic effects in models for QTL mapping and subsequent studies. Moreover, there is low statistical power to detect linkage when markers are in repulsion phase configurations [44, 60]. Although some authors have addressed this problem by including multi-dose (or multiplex) markers when constructing genetic maps and performing QTL mapping [12, 44], the limitations on the genotyping technologies at the time required that the allelic dosage had to be inferred based on expected segregation rates. Because of the high amount of hidden information imposed by marker systems on those studies [44, 60], the estimation of recombination fraction between multi-dose markers was highly impaired.

Quantitative genotyping technologies for single nucleotide polymorphism (SNPs) evaluation have opened the door for further genetic mapping studies in high-level autopolyploids. It is now possible to measure the abundance of specific alleles within a locus in a polyploid genome [2,16,21,37,45,56]. This technology, combined with the genotypic distribution in the population, makes it possible to infer the allelic dosage by using the ratio between the abundances of the two alternative alleles [45]. Once the dosage of the markers is estimated, the construction of linkage maps can be significantly improved by taking this information into account, as previously done in autotetraploids [21] and [19].

Genetic linkage maps can be constructed based on two-point or multipoint estimates of the recombination fraction. Two-point methods use information of pairs of markers, and even though they are less computationally demanding than multipoint methods, they require a higher amount of information in the markers to provide reliable results. Multipoint approaches, instead, use information of multiple markers present in a linkage group, increasing the statistical efficiency of the analysis [23,25,27,36]. This feature is particularly important in polyploid linkage analysis, where markers are mostly partially informative. One widely used procedure to obtain multipoint estimates is the hidden Markov model (HMM) [25]. The construction of the genetic map using this method provides the estimates of the recombination fractions between all adjacent markers in a linkage group, as well the multipoint likelihood, which has been shown to be an excellent criterion to evaluate and compare linkage phase configurations and orders of markers [36]. [27] presented a statistical framework in which HMMs were applied to reconstruct genetic linkage maps, but it was limited to autotetraploids. Recently, software packages such as polymapR [5], pergola [17] and netgaws [3], have been developed to build genetic maps in high-level autopolyploids. However, only polymapR is capable of estimating recombination fractions and inferring parental linkage phases in outcrossing populations, though it does not use multipoint procedures to perform those tasks.

The main challenges we address in this paper are the inference of the haplotypes of the multiple homologous chromosomes and the multipoint estimation of recombination fractions in high-level autopolyploids. Although [65] proposed a probabilistic multilocus haplotype reconstruction model for autotetraploids considering double reduction, this remains as an open question for organisms with higher ploidy levels. Our method relies on an HMM and is developed for species with even ploidy levels under random chromosome segregation (complete polysomic inheritance). We also present a two-point method which is capable of dealing with hundreds of markers even in high ploidy level scenarios. Hence, we are proposing solutions for steps *i* and *iv* in high-level autopolyploids. Step *ii* is straightforward from step *i* using clusterization algorithms, as proposed by [54]. Even though step iii is a challenging task in genetic mapping, it can be addressed using pairwise recombination fractions or the resulting likelihood of the Markov model as it has been proposed by several studies [8,11,26,41,55,58,63]. To evaluate our method, and to show its properties, we rely on simulations of autotetraploid, autohexaploid, and autooctaploid data and on a real tetraploid potato data set. We also perform a set of hexaploid simulations to compare our method to the one implemented in polymapR [5]. The R computer codes to reproduce all simulations and analysis are publicly available.

## Methods

In this section, we define the notation used throughout this article and present the probabilistic model for the gamete formation in autopolyploids. Then, we move to the calculation of the *transition probabilities* for adjacent marker loci (Eq 6) and follow to the *initial state* (Eq 7) and *emission probability* distributions (Eqs 8 and 9) which are fundamental in an HMM. We conclude by explaining the complexity of estimating linkage phases between markers, presenting an efficient two-point algorithm that simplifies the problem in a way that allows the phasing to be inferred using real data.

### Notation

Consider a mapping population derived from a cross between two autopolyploid individuals *P* and *Q* with the same ploidy level (full-sib family). The ploidy level is denoted by *m*, and can be any even number greater than zero. Let the vectors 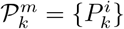 and 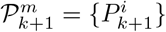, and 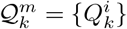 and 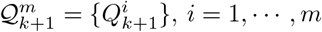, denote the genotype of two adjacent multiallelic loci *k* and *k* + 1 in *P* and *Q*, respectively. The superscript *i* indicates one of the possible alleles for the loci, and each locus has *m* different alleles in each parent. For example, for a cross between two autohexaploid individuals, 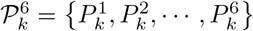; similarly, this can be done for 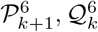 and 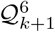. All alleles denoted by the same superscript number are in the same homologous chromosome (e.g., 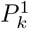 and 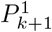 are in homologous chromosome 1, etc). The following assumptions are made to ensure random chromosome segregation [22, 38] and no double reduction [9]: *i*) there is only formation of bivalents during the meiosis; ii) there is no preferential pairing during the formation of bivalents; *iii*) all bivalents have the same recombination fraction between loci *k* and *k* + 1; *iv*) bivalents are independent and *υ*) there is separation of sister chromatids during the meiosis II. Consequences of violations of these assumptions will be addressed later using simulations.

### Bivalent formation

It occurs during meiosis I (more specifically, at the pachytene stage of prophase). In diploid cells, there is only one possible pairing configuration: two duplicated homologous from a homology group pair to form one bivalent. However, in autopolyploid cells, given the previous assumptions, the number of possible pairing configurations, i.e., the number of possible bivalent chromosomal pairing for a given homology group during meiosis is

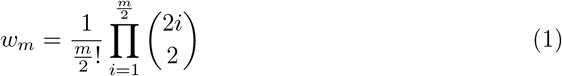

The orientation of the bivalents does not affect the expected frequencies of each gamete type, and therefore will not be considered. For example, for an autotetraploid individual, there are two bivalents and three possible bivalent configurations. Homologous chromosome pair as 1 with 2, and 3 with 4; or, 1 with 3 and 2 with 4; or 1 with 4 and 2 with 3 [18]. We denote Ψ = {*ψ_j_*}, *j* = 1, …, *w_m_* a set of all bivalent configurations for a given ploidy level.

### Expected gametic frequency for a given bivalent configuration

We will present the expected gametic frequencies considering parent *P*. Since parent *Q* undergoes a similar process, it is possible to combine the expected gametic frequencies to obtain the expected genotypic frequency in the full-sib population. Each of the bivalents obtained for a given configuration *ψ_j_* can result in two types of chromosomes for loci *k* and *k* + 1: *parental*, which results from bivalents with zero or any other even number of recombinations between *k* and *k* + 1; and *recombinants,* which results from bivalents with any odd number of recombinations. As presented by [12], the probabilities of all chromosome types for any single bivalent can be represented always as

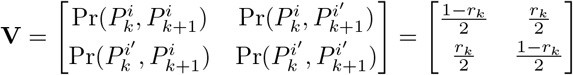

where *r_k_* is the recombination fraction between *k* and *k* +1, *i* = *i*′. For a given configuration *ψ_j_*, the expected frequencies for all possible gametes derived from that configuration is

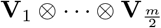

where ⊗ denotes the Kronecker product of matrices and subscripts in V indicate the corresponding bivalent. All elements of this product are of the form

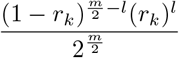

where *I* denotes the number of total recombinant bivalents between loci *k* and *k* + 1, *I* ∈ {0, …, *m*/2}. From this, we can define the probability of observing any gamete (for two loci) given a bivalent configuration *ψ_j_* as

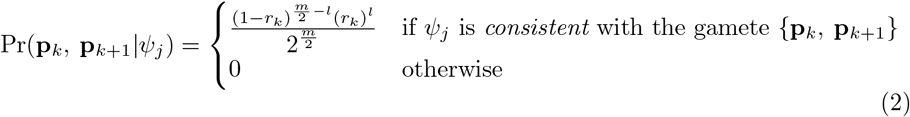

where vectors **p**_*k*_ and **p**_*k*+1_ denote a subset of 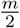 alleles present in 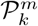 and 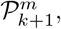, respectively; {**p**_*k*_, **p**_*k*+1_} indicates a gamete for loci *k* and *k* + 1 from parent *P*. *Consistent* means that the gamete can be produced from bivalent configuration *ψ_j_*. Notice that some gametes cannot be obtained from *ψ_j_* once the bivalents are formed.

Since we assume that alleles with the same superscript are in the same homologous chromosome, *l* can be obtained by a simple examination of superscripts of elements contained in **p**_*k*_ and **p**_*k*+1_. Consider, for example, *ψ*_1_ = {(1, 2), (3,4), (5,6)} (*m* = 6, Fig 1). If one observes 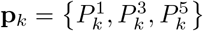 and 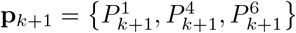, the number of recombinant chromosomes is *l* = 2. Therefore, 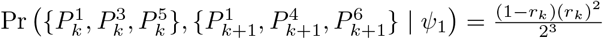. On the other hand, 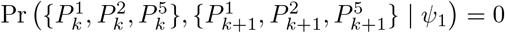, since it is impossible to obtain this gamete from configuration *ψ*_1_, i.e., it is not *consistent* with *ψ*_1_.

**Figure 1.**
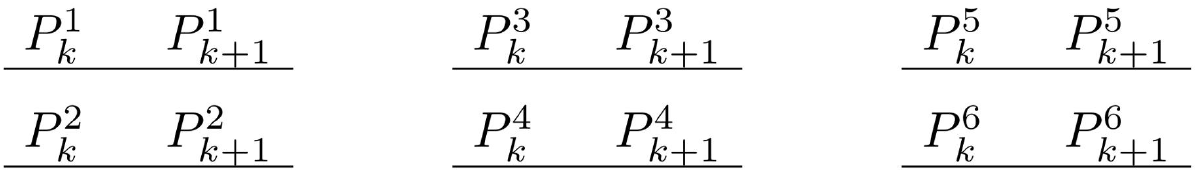
One possible pairing configuration in an autohexaploid, namely *ψ*_1_. 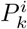 denotes one allele present in homologous chromosome *i* for locus *k* in parent *P*. Notice that some allelic configurations, such as 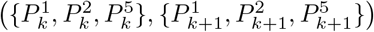, are impossible to be obtained in this bivalent pairing. In this case, the homologous chromosomes containing alleles 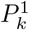 and 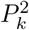 will migrate to opposite poles of the cell during meiosis I. Therefore, 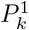 and 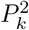 will not be present in the same gamete.

### Gametic frequency unconditional to bivalent configurations

In reality *ψ_j_* is unknown, thus the conditional probability given by Eq (2) must be considered for all possible *ψ_j_*. The probability of observing a gamete {**p**_*k*_, **p**_*k*+1_}, unconditional to *ψ_j_*, can be expressed as

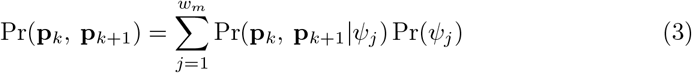

It is important to notice that only a subset of Ψ is *consistent* with the observed gamete, and consequently Pr(**p**_*k*_, **p**_*k*+1_ | *ψ_j_*) > 0 only for some **ψ_j_**’s. Fig 2 shows a graphical representation of Eqs 2 and 3 for autohexaploid gametes.

**Figure 2.**
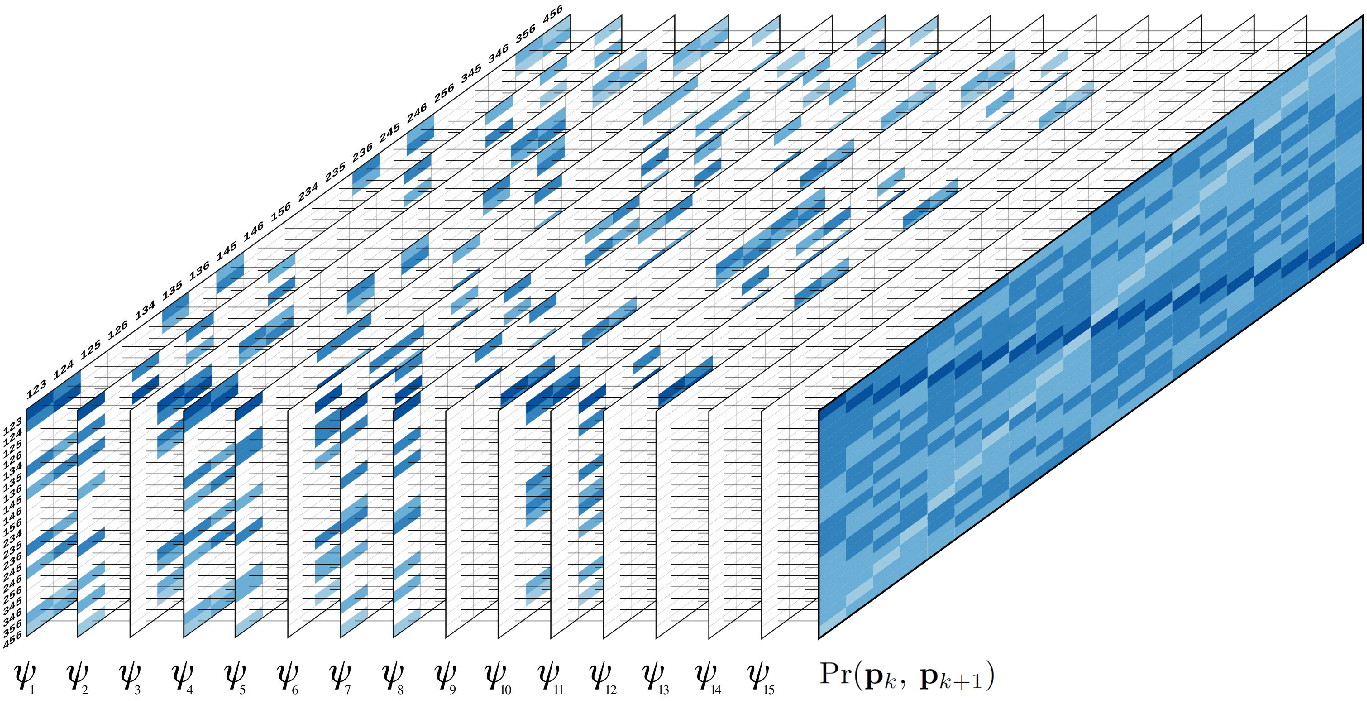
Graphical representation of Eqs 2 and 3 for autohexaploid gametes. The first 15 tables represent the gametic probabilities given different bivalent configurations *ψ*. (Eq 2). The rows and the columns indicate gametic configurations for loci *k* and *k* + 1, respectively. For simplification, only the superscripts of the gametic configurations were presented. For example, row 123, column 123, represent the gamete 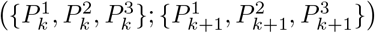. Colored cells indicate the probability of gametic configurations *consistent* with the bivalent configuration *ψ*. The color scale indicates the number of recombinant bivalents associated to the gametic probability varying from 0 (dark blue) to 3 (light blue). Blank cells indicate *non-consistent* configurations. The far right full table represents the sum over all *ψ* configurations, weighted by their probability (Eq 3).

The probability of observing a specific gamete is always the same for each *ψ_j_* in this consistent subset (Eq 2). Therefore, under random pairing (assumption ii), our task reduces to finding the number of elements in this subset that are consistent with the observed gamete and multiply Pr(**p**_*k*_, **p**_*k*+1_ |*ψ_j_*) Pr(*ψ_j_*) by this number. The result is the probability of observing a gamete unconditional to the bivalent configuration.

For every gamete, *l* can change from zero to *m*/2 recombinant homologous chromosomes. The observed gamete is the result of homologous chromosomes that migrate to one pole of the cell at anaphase I. Since we are assuming that there is separation of sister chromatids during anaphase II, if *l* = 0 (all chromosomes are of parental type), there is no information about the pairing configuration of the homologous chromosomes that migrate to the opposite pole of the cell. In this situation, there are 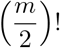 possible pairing configurations, and the number of possible *ψ_j_* that can. produce gametes with *l* = 0 is 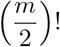. Therefore, for *l* > 0, there are 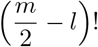 possible pairing configurations of parental chromosomes. For the remaining *l* recombinant chromosomes, the number of possible pairing configurations is *l*!. Thus, the total number of possible pairing configurations that can produce a specific gamete is 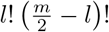. This is precisely the number of elements in the subset of Ψ consistent with the observed gamete. Given the assumption of no preferential pairing during the formation of bivalents, 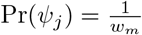, the probability of a gamete {**p**_*k*_, **p**_*k*+1_}, unconditional to *ψ_j_*, can be simplified to

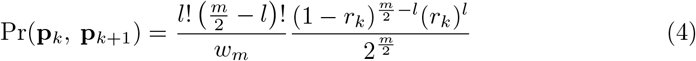

### Map reconstruction via hidden Markov model

The construction of a genetic map involves the estimation of the genetic distance and order between markers within linkage groups. If the origin of the haplotypes (i.e., linkage phase) for the parents of the mapping population is unknown, it also needs to be estimated. For several years, hidden Markov models have been proven to be an excellent avenue for obtaining these estimates [23,25,27,36]. The multipoint likelihood obtained using HMMs is employable as a criterion to compare marker orders and judge which one is best, and also to provide a reliable estimation of recombination fraction and linkage phases. [42] defines an HMM as a generative process composed of three well-defined probability distributions: *transition*, *initial state* and *emission*. In genetic mapping context, the transition probability distribution is defined as the probability of having a particular genotype at position *k* + 1, given the genotype at position *k*. Using Eq (4) the gametic transition probabilities Pr(**p**_*k*+1_|**p**_*k*_), or the conditional probability of a gamete genotype at locus *k* + 1 given the gamete genotype at locus k, is simply

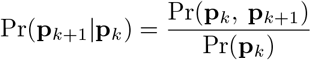

Under random chromosome segregation, both **p**_*k*_ and **p**_*k*+1_ can have 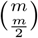 different genotypes. Let 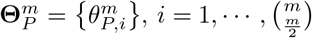 denote all possible genotypes that **p**_*k*_ can assume for locus *k*. Also, assume that genotypes in 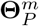 are arranged according to the lexicographical order of their superscripts. For example, in an autotetraploid, 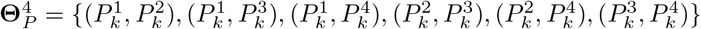 for locus *k*. After some simplifications (see Supplementary Information, File S1) the transition probability, i.e., the conditional probability of a gametic genotype 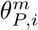 in locus *k* +1 given the gametic genotype 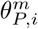, in locus *k*, is

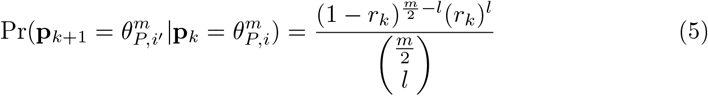

where 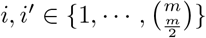. The initial state and the emission probability distributions will be addressed in the next section (Eqs 7 to 9).

### Including information of both parents

Any given individual in a full-sib population is formed by the union of gametes from both parents, *P* and *Q*. Each parent can form 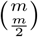 different gametes for locus *k*. Since the formation of gametes in both parents is independent, the genotypic transition probability distribution can be written as

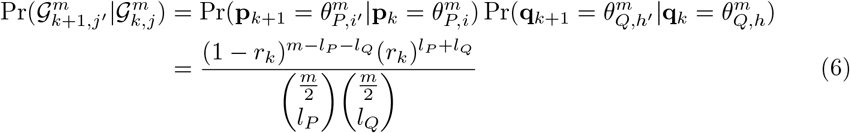

where 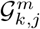 denotes the genotype of an individual derived from the union of gametes 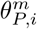 and 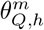 at locus *k*. The same reasoning applies to 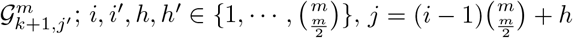, and 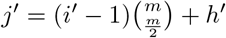. *I_p_* and *I_q_* denote the number of recombinant bivalents between loci *k* and *k* + 1 in parents *P* and *Q*, respectively. Let 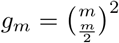 denote the number of possible genotypes derived from the cross between 2 individuals *P* and *Q*. For simplification and without loss of generality, let 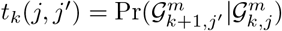. For a comprehensive example of the transition probabilities and the indexation used in Eq. 6, see Table S3.8, File S3 in Supplementary Information.

Given a ploidy level *m* and a recombination fraction *r_k_*, the only information required to obtain *t_k_*(*j, j*’) in Eq (6) is *l_P_* and *l_q_*. Since the genotypes in 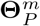 and 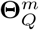 are arranged according to the lexicographical order of their superscripts, it is possible to obtain (*l_P_, l_q_*) for any given pair (*j, j*’) using the algorithm presented in Supplementary Information, File S2. Although the number of possible transitions between positions *k* and *k* + 1 is (*g_m_*)^2^, which can be a very large number even for modest ploidy levels, it is possible to obtain the transition between any specific genotypes in *j* and *j*′ without computing the entirety of the transition space.

The initial state distribution is the probability of observing a specific genotype. Given the assumption that there is no preferential pairing during the formation of bivalents, a uniform probability density function can be employed as the initial state probability function

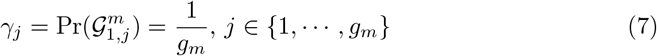

To this point, both transition and initial state distributions consider different allelic variants for all *m* homologous chromosomes in both parents. This scenario can only be achieved when using fully informative markers. In reality, autopolyploid species may have the same allelic variant in some homologous chromosomes. Besides, even if all homologous have different allelic forms, modern genotyping platforms are usually capable of detecting polymorphisms at the nucleotide level (SNPs), which are essentially biallelic. Due to this lack of identity between the observed data and the full transition space, we make use of the emission function, which is defined as the probability of observing a molecular phenotype given a genotype 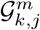.

The detection of the allelic variants in modern genotyping platforms is based on the abundance of different alternative nucleotides. In the autopolyploid setting, this can be translated as the *dosage* of a SNP at a specific locus. The dosage of a SNP can be estimated using the ratio between the abundance of its two allelic forms. Several methods were proposed to perform this task including [56], [45] and [2]. Here we introduce a biallelic derivation of the emission probability distribution. Although the function presented here use biallelic information, other distributions can be derived for partial informative multiallelic marker systems following the same reasoning.

Let 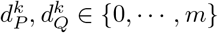 denote the *observed dosage* of one allelic form in locus *k* for parents *P* and *Q*, respectively. The choice of the allelic form denoted by 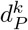 is arbitrary, as long as the same allelic form is used in 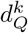. The dosage observed in parent *P* can be originated from alleles present in 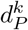 of the *m* homologous chromosomes. Let 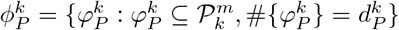 denote a set of size 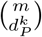 containing all possible subsets in 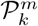 that originate the observed dosage 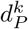. The operator #{.} is the cardinality of a set. The same reasoning applies for 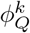. For instance, in an autotetraploid, if 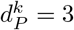, the three doses present in locus *k* can be derived from four distinct subsets 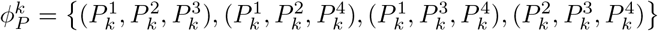. Given two particular subsets 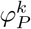 and 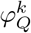 in 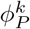 and 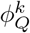, each one of the *g_m_* genotypic states in the full transition space can be associated to a dosage. The dosage associated to the *j*-th state is obtained by counting the number of alleles present in the intersection between the parental allelic set 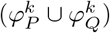 and 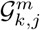. Thus, the emission function can be defined as

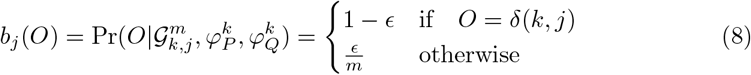

where 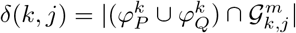 and *ϵ* denotes the global genotype error rate. In addition to the punctual estimate of the dosage, the genotyping calling methods cited above also provide the probability distribution of the dosages for a particular marker for all individuals of the biparental population. If this information is available, a more general emission function can be derived. Instead of modeling a global error rate *ϵ*, we use the prior information provided by the genotyping calling procedure. Let 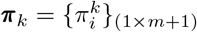 denote the probability distribution vector associated to the dosages 0, · · ·, *m* at position *k* for a particular individual in the biparental population. For example, 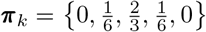 denotes a tetraploid individual with probabilities 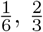 and 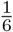 of having one, two and three doses, respectively, and zero for the remaining ones. Then, the emission probability function can be written as

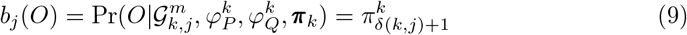

In this case, the observation *O* can be any dosage from 0 to *m* and the information about the genotypes will be contained in the probability distribution of the dosages **π**_*k*_. Thus, the probability of observing any dosage given a genotype 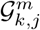 associated to a particular dosage *δ*(*k, j*) can be obtained by simply assessing the corresponding value in the probability distribution provided by the genotype calling procedure. Notice that Eq 8 can be reduced to Eq 9 using the appropriate **π**_*k*_. For example, in autotetraploids, when the observed dosage for locus *k* is one, *O* = 1, 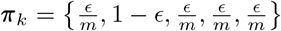. Moreover, for missing values, it is possible to use the probability distribution of the genotypic classes under polysomic segregation, as presented by [45].

### Multipoint likelihood and the estimation of recombination fraction

Suppose there are *z* markers in a homology group in a known order represented by *M*_1_, …, *M_k_*, …, *M_z_*. Let *r* = (*r*_1_, …, *r_k_*, …, *r*_*z*-1_) denote the recombination fraction vector between all marker intervals in this sequence. Also, assume linkage phase configurations in parents *P* and *Q* denoted respectively by 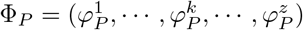 and 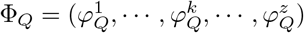. The sequence of observations for the *z* markers is denoted by (*O*_1_, …, *O_k_*, …, *O_z_*) and its underlying probability distributions is denoted by Π = (**π**_1_, …, **π**_k_, …, **π**_*z*_). The likelihood of *M*_1_, …, *M_k_*, …, *M_z_* can be obtained using Eqs (6), (7) and (9) following the classical *forward procedure* [42]. Let 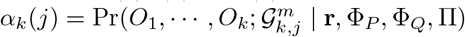 denote the probability of the partial observation sequence (*O*_1_, …, *O_k_*) and genotype 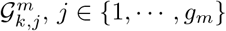 given the sequence of recombination fractions r, the linkage phase configurations Φ_*P*_ and Φ_*Q*_ and the probability distributions for the sequence of observations Π. The forward procedure follows the steps below:

1. Initialization:

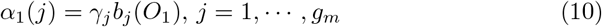
2. Induction:

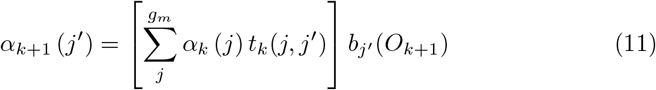

where *k* = 1,…, *z* – 1 and j′ = 1, …, *g_m_*
3. Termination:

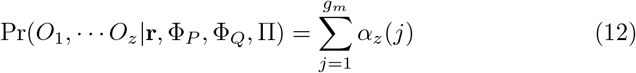

Then, the likelihood of the model is defined as

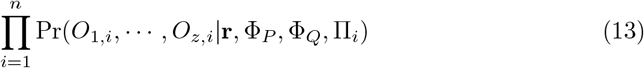

where *n* is the number of individuals in the full-sib population, *O*_1,*i*_, …, *O*_*z,i*_ is the sequence of marker observations for individual *i* and Π_*i*_ is a (*m* +1) × *z* matrix where the *k*-th column denotes the probability distributions associated to the marker *M_k_*, individual i. The multipoint maximum likelihood estimate of *r* can be obtained using the *forward-backward* procedure coupled with the EM algorithm [42]. For the backward procedure, consider the variable 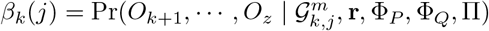 as the probability of the partial observation sequence from *k* + 1 to *z*, given the genotype 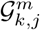, the recombination fraction vector *r*, the linkage phase configurations Φ_*P*_ and Φ_*Q*_ and the probability distributions for the sequence of observations Π. The solution to *β_k_*(*j*) was also described by [42] as follows:

1. Initialization:

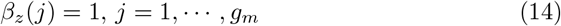
2. Induction:

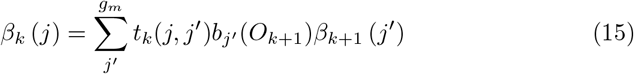

where *k* = *z* –1, *z* – 2, …, 1 and *j* = 1, …, *g_m_*

To estimate the recombination fraction for all intervals in the marker sequence we need to define *ξ_k_*(*j, j*’) as the probability of state 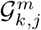 at position *k* and state 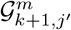 at position *k* +1 given the sequence of observations *O*_1_, … *O_z_* and their underlying probability distributions Π, the recombination fraction vector r and the linkage phase configurations Φ_*P*_ and Φ_*Q*_

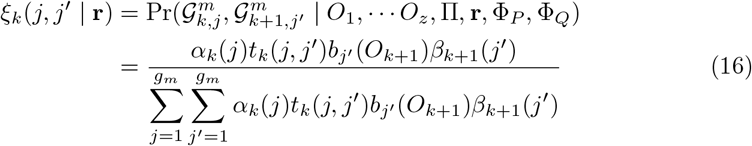

The recombination frequency *r_k_* can be estimated through an iterative process using

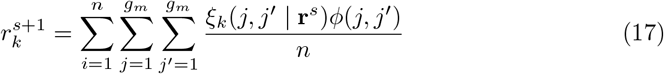

where *ξ_k_* (*j, j*’ | **r**^*s*^) is calculated for individual 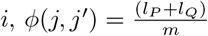 is the proportion of recombinations between markers *k* and *k* + 1 for individuals with genotypes 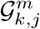 and 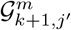 and **r**^*s*^ is the vector of recombination fractions in the iteration (*s*) and **r**^*s*+1^ is the updated recombination fraction vector [7].

### Estimation of linkage phase

Let the Cartesian product 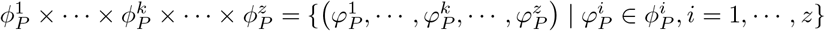 denotes a set containing all possible linkage phase configurations in parent *P*. Also, let 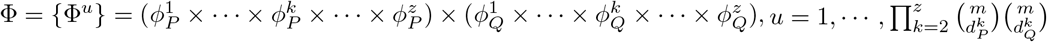, denote a set containing all possible linkage phase configurations in both parents. The probability of the linkage phase configurations can be obtained using Bayes’ rule

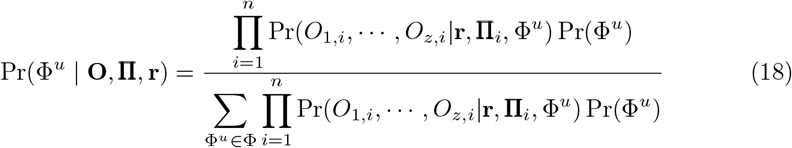

where **O** is an array containing the observation for *z* markers in *n* individuals, and Π is the underlying probability distribution for all marker observations. Since the prior probability Pr(Φ^*u*^) can be assumed to be uniform, the posterior probability is proportional to the likelihood of the model, which can be used to select the best linkage phase configuration. Depending on the dosage and number of markers, some of these configurations are equivalent and will result in the same likelihood. The search space for the best linkage phase configuration can be unwieldy depending on the ploidy level, dosage and number of markers. Also, the transition space on the HMM gets larger as the ploidy level increases. To circumvent these problems, we propose a very efficient two-point procedure to reduce the search space for linkage phases.

### Two-point algorithm for high-level autopolyploids

When the linkage analysis is conducted only between two markers (two-point analysis), the information contained in these markers does not propagate into the rest of the chain. Thus, based on the dosage and linkage phase configuration of the markers involved in the analysis, the *g_m_* genotypic states present in the full transition space can be collapsed into a small number of states, and a straightforward likelihood function can be derived. It is worthwhile to mention that the estimates obtained using the two-point procedure are the same as those obtained using the multipoint algorithm for two markers. However, the two-point computation is extremely faster.

Consider a biallelic marker in an autopolyploid biparental cross with ploidy *m*. The number of possible genotypic states in the progeny for a given locus at position *k* is 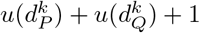, where the operator 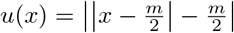 and |·| denotes module. For example, in an autohexaploid biparental cross, if the dosage of the marker at position *k* in parent *P* is two 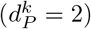 and in parent *Q* is three 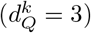, the number of possible genotypic classes expected in the progeny is six. Depending on the linkage phase configuration, each of the *g_m_* genotypic states in the full transition space corresponds to one of these expected genotypic classes, as presented in the emission function (Eqs 8 and 9). Thus, in the previous example, all the *g_m_* states could be collapsed into six different classes. To perform this reduction of dimensionality, let 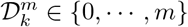 denote one of the possible genotypes based on the dosage of one individual in the progeny of an autopolyploid biparental cross for position *k* with ploidy *m*. The joint probability of 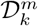 and 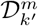, for a given genotypic configuration at positions *k* and *k*′ can be written as

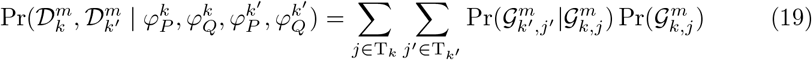

where 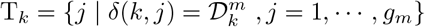 and *δ*(*k, j*) was defined in Eq 8; the same applies to T_*k′*_. Since in a two-point analysis the probability distribution of the genotypic states in locus *k* can be assumed to be uniform, i.e., 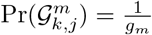, Eq (19) can be rewritten as a sum of weighted terms from Eq (6)

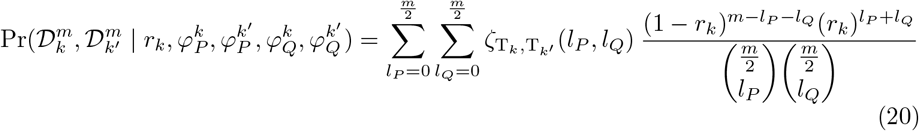

where

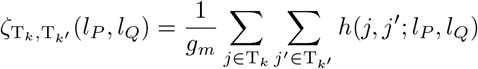

*h*(*j, j*’; *l_P_,I_Q_*) is 1 if (*j,j*’) corresponds to (*l_P_, l_Q_*) according to the procedure described in Supplementary Information, File S2, and zero otherwise. Eq 20 can be expressed in matrix form as

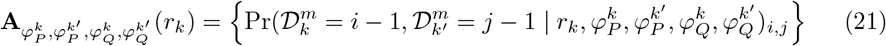

where 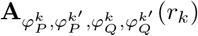 is a (*m* +1) × (*m* +1) matrix. Yet, in a two-point analysis with biallelic markers, the linkage phase configuration can be summarized in an ordered pair 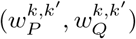 indicating the number of homologous chromosomes that share allelic variants for loci *k* and *k*′ in parents *P* and *Q*, respectively. For a given pair 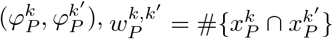, where 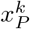 and 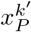 denote the set of homologous chromosomes inherited by parent *P* in positions *k* and *k*′, which can be assessed using the superscripts in 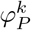 and 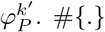 indicates the cardinality of the set. Notice that 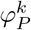 and 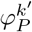 can assume several linkage phase configurations resulting in the same 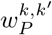. Let 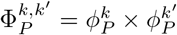 denote a set containing all possible pairs 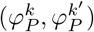 for a given pair 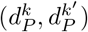. In this set, there are 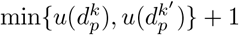 partitions, each one corresponding to a different 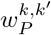. Fig 3 shows an example of 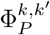 for 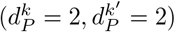 in an autotetraploid homology group. The size of the set is 36, and it can be subdivided into three partitions where 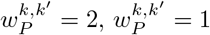 and 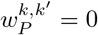.

In a two-point context, the likelihood function derived from any of the configurations belonging to the same partition (same 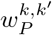) will be the same. Thus, any of them can be used to obtain the likelihood function for a given 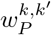. Let 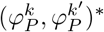 denote one of the possible pairs 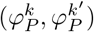 that correspond to 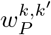. The same reasoning applies to parent *Q*. Without loss of generality, the two-point likelihood function of biallelic observed molecular phenotypes for markers *k* and *k*’ given 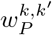 and 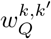 is

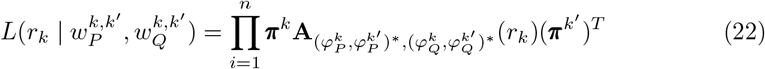

where *n* is the number of individuals and *T* denotes transposition of a vector. In Eq (22), *r_k_* can be estimated using iterative procedures such as EM or Newton-Raphson. As in Eq (18), it is possible to list all linkage phase configurations and evaluate them based on their likelihood. Here we use the LOD Score (base-10 logarithm of likelihood ratios) in relation to the highest likelihood. Thus, models with high likelihoods will yield LOD Scores close to zero. We also use the LOD Score to assess the evidence for linkage between the two markers using the ratio between the model under 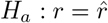 and under the null hypothesis of no linkage *H_o_*: *r* = 0.5, given a linkage phase configuration.

As previously shown, it is possible to enumerate all linkage phase configurations for parent *P* using the Cartesian product 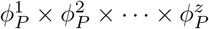. To reduce this Cartesian space based on two-point analysis, we add a restriction where all pairs 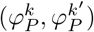 in a sequence of configurations 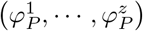 must be contained in 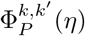, where 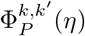 is a subset of all partitions in 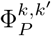 in which the associated LOD Score is smaller than *n* Thus, a reduced subset of linkage phases in parent *P* based on two-point analysis can be obtained using

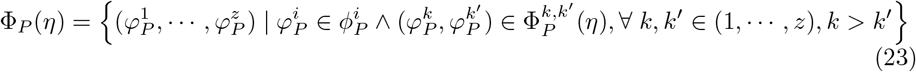

It is important to note that it is not necessary to represent the whole Cartesian space {Φ_*P*_} to restrict the linkage phase configurations to the condition 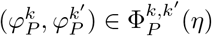. This procedure can be done through the sequential addition of markers from *M*_1_ to *M_z_*. For each marker added to the end of the chain, the ordered pair (*k, k*’), *k*’ = 2, …, *z* and *k* = *k*’ – 1, …, 1, is evaluated and only linkage phase configurations that meet the condition 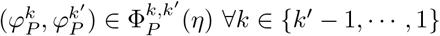 are considered.

Some of the configurations selected using the previous procedure can be equivalent once they are products of a permutation of the same set of homologous chromosomes. In order to remove this redundancy, let each one of the selected configurations be represented as a binary matrix of dimensions (*m* × *k*’) such as

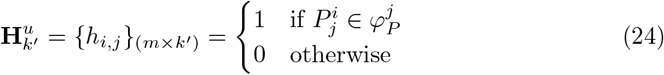

where *u* ∈{1, …, *U*}, *U* is the number of selected linkage phase configurations, and *k*’ indicates that *M_k′_*, was the last marker inserted in the chain. The rows of matrix 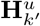, represent the homologous chromosomes for the u-th linkage phase configuration with the insertion of the *k*′-th marker at the end of chain; 1 denotes the presence of an allelic variation, and 0 denotes its absence. If a matrix **H**_k′_, could be obtained from a matrix 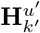, just by permuting the rows (permuting the order of the homologous chromosomes), these two linkage configurations yield the same likelihood. Thus, one of the configurations should be excluded from consideration. The same reasoning applies to parent *Q*. This procedure can be done recursively until all redundancy is eliminated. The reduced linkage phase configurations search space considering both parents is obtained using Φ(*η*) = Φ_*P*_(η) × Φ_*Q*_(*η*), such as #{Φ(*η*)} ≪ #{Φ}, combined with the redundancy elimination for homology groups. This sequential procedure results in a set of linkage phase configurations containing markers up to *M_k′_*, which are evaluated using the HMM likelihood. A LOD Score threshold in relation to the most likely configuration is assumed to determine which configurations should be taken into consideration in the next round of marker inclusion (Fig. 4). Additionally, it is possible to limit the two-point search space reduction to a window of SNPs in the terminal part of the chain to speed up the phasing process. Finally, with all markers inserted, the multipoint likelihood of the whole map is used to find the best configuration among the remaining ones, and the recombination fractions are reestimated. To demonstrate the mechanics of the two-point analysis coupled with the multipoint procedure, a simple example is presented in Supplementary Information, File S3.

**Figure 3.**
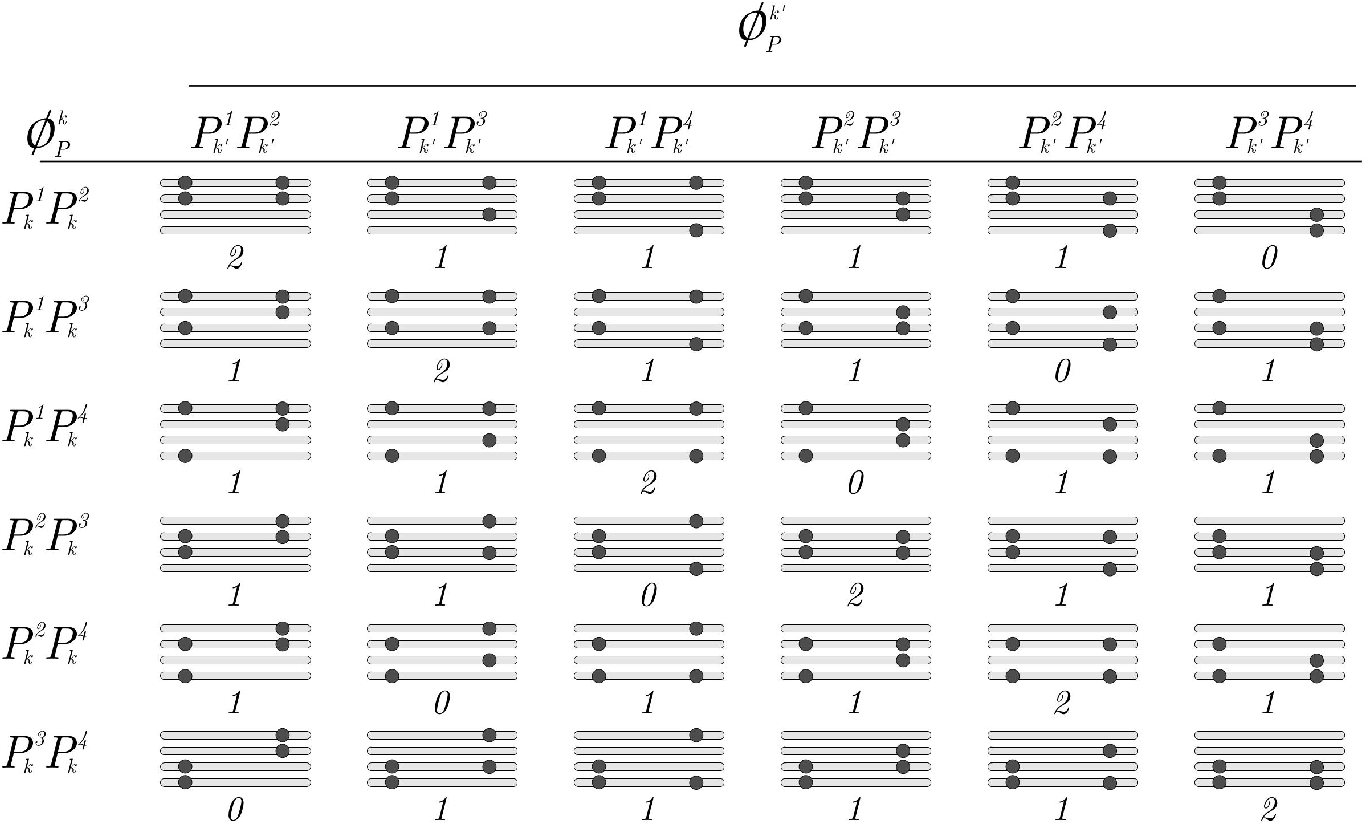
Example of 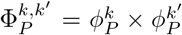 for an autotetraploid homology group with observed dosages 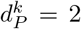 and 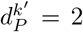 homologous chromosomes sharing alleles. In this case, 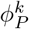 denotes a set of size six, containing all possible subsets of size two in 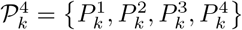. The same reasoning applies to 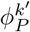. The horizontal bars represent homologous chromosomes forming a homology group and the dots represent allelic variations of a biallelic marker. The number below each homology group represents the number of homologous chromosomes that share allelic variants 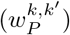. This defines three partitions: 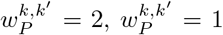 and 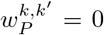. Notice that, from a homology group within a specific partition, it is possible to obtain the same linkage phase configuration observed in another homology group within that partition by permuting the its homologous chromosomes

**Figure 4.**
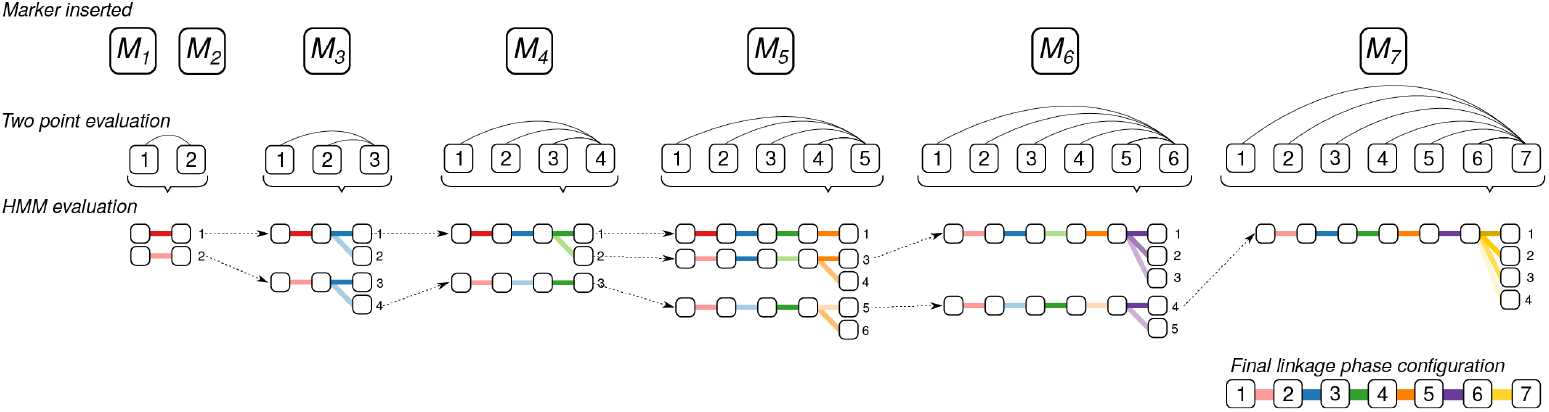
Example of linkage phase configuration estimation using sequential search space reduction and HMM evaluation. Only one parent is presented. The two-point search reduction is composed of two parts: the first one evaluates the LOD Scores obtained through pairwise recombination fraction likelihoods. The second detects equivalent configurations by performing all possible permutations of the homologous chromosomes. The remaining configurations are evaluated using the HMM-based likelihood. In the first step, linkage phase configurations of *M*_1_ and *M*_2_ are evaluated using the two-point analysis. Color shades indicate different linkage phase configurations provided by the two-point analysis. In this example, there are two possible linkage phases represented by two shades of red. In the second step, we evaluate the linkage phases between markers *M*_3_ and *M*_2_, and *M*_3_ and *M*_1_. Configurations with LOD Scores smaller than *η* are maintained to be evaluated by HMM. There are two possible linkage phases given a certain *η*, represented by two shades of blue. These two configurations are combined with the configurations from the previous step, resulting in four configurations evaluated using HMM likelihood. Given a likelihood threshold, only configurations 1 and 4 are eligible for the next step. The same reasoning applies for the remaining markers. A final linkage phase configuration is obtained after inserting the last marker and choosing the one that yields the highest HMM-based likelihood.

## Simulations

### Simulation 1 - local performance under random bivalent pairing

the aim of this simulation study was to evaluate the local performance of the algorithm considering three ploidy levels (*m* = 4, *m* = 6 and *m* = 8) under the mapping model assumptions (i.e., random pairing and bivalent formation). To be in accordance with molecular data that have been made available through sequence technologies, we simulated bi-allelic markers that can be observed in terms of dosage in parents and progeny. Three different linkage phase scenarios were simulated. In scenario A, for each marker, one of the allelic variants was assigned to the first homologous chromosome in the homology group and the remaining variants of the same type were assigned to the subsequent homologous chromosomes. In scenario B, the allelic variant was randomly assigned to one of the first 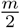 homologous chromosome and the remaining were assigned to the subsequent homologous chromosomes. In scenario C, the allelic variants were randomly assigned to the *m* homologous chromosomes. Thus, it is expected an increasing difficulty to detect recombination events from scenario A, where the allelic variants were concentrated in the same homologous chromosomes, to scenario C, where they were randomly distributed.

For each combination of ploidy level and linkage phase scenario, we simulated five different parental haplotypes. In total, 45 parental configurations were considered (3 × 3 × 5, Supplementary Information, Figure S4). For autotetraploid and autohexaploid configurations, we simulated 1000 full-sib populations. For autooctaploids, this number was reduced to 200 due to the high demand of computer processing required to reconstruct such maps. Each population was comprised of 200 individuals with one linkage group containing 10 markers positioned at a fixed distance of 1 centimorgan (cM) between them. For each combination, the percentage of correctly estimated linkage phase configuration in each parent was recorded. Also, for the cases where the linkage phases were correctly estimated, we calculated the average Euclidean distance between the distances of the estimated and simulated maps using 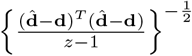 where 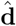 is the vector of distances for a estimated map, **d** is the vector of distances for the simulated map, *z* is the number of markers and *T* indicates vector transposition. For example, a value of 1 cM indicates that the maps differ 1 cM in average from each other [36]. We used the sequential two-point procedure to reduce the search space assuming that linkage phase configurations with associated *LOD* < 3.0 should be investigated using HMM multipoint strategies (η = 3). For the remaining configurations evaluated using HMM, we kept those with *LOD* < 10.0 to be evaluated in the next round of marker insertion.

### Simulation 2 - chromosome-wise performance under preferential pairing and multivalent formation

In this simulation study, we evaluated the performance of the algorithm in dense maps, allowing for multivalent formation and preferential pairing. We used Scenario C from the previous study as a template to simulate five tetraploid and five hexaploid parental haplotypic configurations, each one comprising 200 equally spaced markers with a final length of 100.0 cM (Supplementary Information, Figure S5). For each parental configuration, we simulated 200 full-sib populations of 200 offspring considering a combination of three levels of preferential pairing (0.00, 0.25 and 0. 50) and three levels of cross-like quadrivalent formation proportion (0.00, 0.25 and 0. 50). No hexavalents were simulated in this study. For autohexaploids, the multivalent configurations were always composed by a cross-like quadrivalent plus a bivalent. The centromere was positioned at 20.0 cM from the beginning of the chromosome (subtelocentric centromere with arms ratio 1:4) to study the effect of the double reduction at the distal end of both chromosome arms. All simulations were conducted using the software PedigreeSim [57]. In addition to the statistics recorded in Simulation 1, we computed the rate of double reduction observed in each marker for all constructed maps using the “founderalleles” file provided by PedigreeSim. We also evaluate two values for the LOD Score threshold associated to the two-point analysis (*η* = 3 and *η* = 5). We used a multipoint LOD Score threshold of 10.0 and also limited the two-point search to a 50 SNP window in the terminal part of the map.

### Simulation results

#### Simulation 1

Table 1 shows the percentage of data sets where the linkage phase configuration was correctly estimated in both parents *P* and *Q*. In scenario (A) the method was capable of recovering the correct linkage phase configuration in all situations for all ploidy levels. In scenarios (B) and (C) there was a slight decrease on the ability to correctly estimate the linkage phase configuration, especially for *m* = 6 and *m* = 8. Although in these cases the percentage of correctly estimated linkage phases was lower, the numbers are considerably high, varying from 100% to 88.8%. This indicates a very good performance to estimate the linkage phase configurations, even using the two-point procedure to narrow the search space.

**Table 1.** Percentage of data sets where linkage phase configuration was correctly estimated for five different parental P and Q haplotypes in simulation 1.

| Ploidy level | A |  | B |  | C |  |
| --- | --- | --- | --- | --- | --- | --- |
| | $P$ | $Q$ | $P$ | $Q$ | $P$ | $Q$ |
| Autotetraploid<br>($m = 4$) | 100 | 100 | 99.7 | 99.8 | 100 | 100 |
|  | 100 | 100 | 99.7 | 99.7 | 99.9 | 99.7 |
|  | 100 | 100 | 100 | 100 | 99.7 | 99.8 |
|  | 100 | 100 | 99.9 | 99.7 | 99.9 | 99.9 |
|  | 100 | 100 | 99.9 | 99.8 | 100 | 100 |
| Autohexaploid<br>($m = 6$) | 100 | 100 | 96.6 | 97.2 | 96.1 | 94.6 |
|  | 100 | 100 | 97.3 | 97.5 | 95.8 | 95.6 |
|  | 100 | 100 | 96.5 | 96.7 | 94.7 | 94.6 |
|  | 100 | 100 | 97.3 | 97.4 | 96.1 | 94.7 |
|  | 100 | 100 | 97.2 | 97.4 | 95.2 | 94.5 |
| Autooctaploid<br>($m = 8$) | 100 | 100 | 93.6 | 94.4 | 93.2 | 95.7 |
|  | 100 | 100 | 97.6 | 96.8 | 92.1 | 93.9 |
|  | 100 | 100 | 96.8 | 97.6 | 90.4 | 89.2 |
|  | 100 | 100 | 97.7 | 98.4 | 90.6 | 90.0 |
|  | 100 | 100 | 96.9 | 94.6 | 88.8 | 90.6 |

Fig 5 shows the distributions of the average Euclidean distances between the estimated and simulated distance vectors for the correctly estimated linkage phase configuration. In all cases, the majority of the recombination fractions were consistently estimated once the medians of all distributions are very close 0.5, with no practical problems in terms of mapping construction. These results show that, apart from a relatively small percentage of entangled linkage phase configurations, the method successfully performed the phasing and managed to estimate the recombination fraction of 10 markers in all situations evaluated.

**Figure 5.**
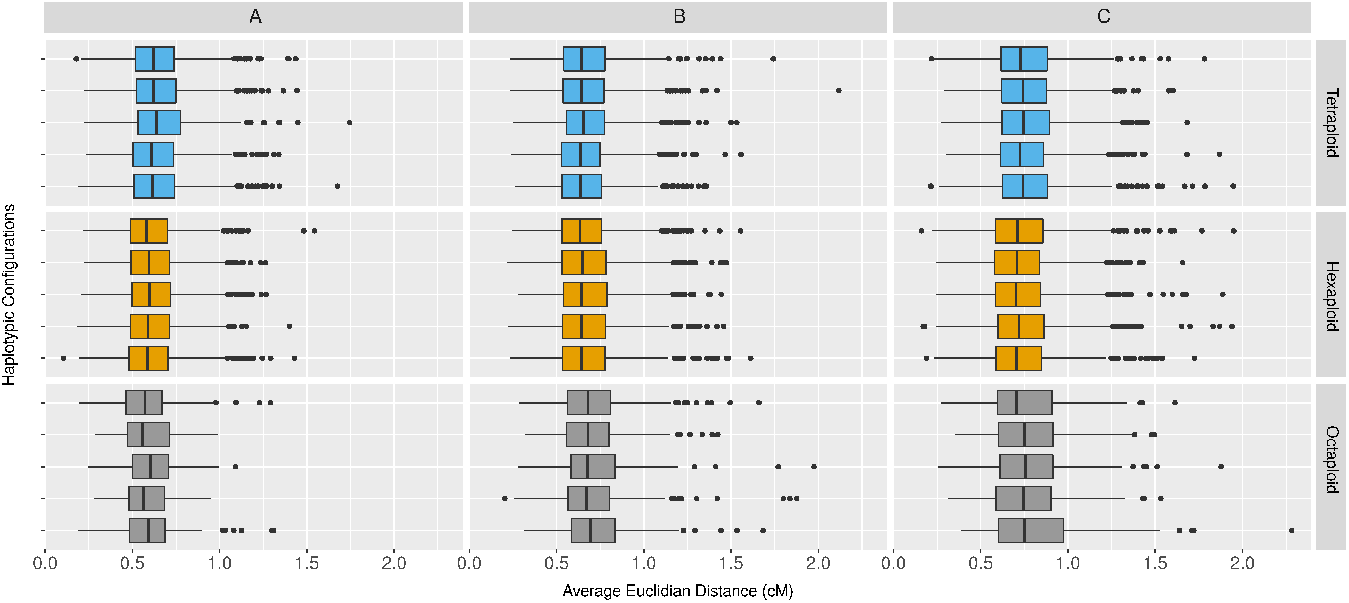
Distributions of the average Euclidean distances between the estimated and simulated distance vectors considering correctly estimated linkage phase configurations. The order of boxplots is the same as the order of haplotypes in Figure S4. Each column indicates the results for different linkage phase configuration scenarios, namely, A, B and C, and each row indicates a different haplotypic configuration within three ploidy levels.

#### Simulation 2

The proportion of correctly estimated linkage phase configurations for the dense chromosome-wise map is shown in Table 2. In general, results for tetraploid maps were superior when compared to results for hexaploid maps. It is also possible to observe a better performance for the threshold level *η* = 5 in comparison to *η* = 3. Similarly to Simulation 1, maps resulting from configurations with no preferential pairing or quadrivalent formation showed a high proportion of correctly estimated linkage phase configurations. Results ranged from 100% to 99% for tetraploid maps and from 100% to 84% for hexaploid maps. Different levels of quadrivalent formation rate had no substantial influence in estimating the correct linkage phase configurations in tetraploids. Within the preferential pairing level 0.0, the percentage of maps with correct linkage phases varied from 100% to 90%. For hexaploids, there was a decrease in this percentage as the quadrivalent formation increases from 0.0 to 0.50, with proportions varying from 100% to 70.5%. Especially for autohexaploids, there was a considerable variation between the five simulated configurations. This occurred, because the effect of the quadrivalent formation can be more pronounced depending on the level of information contained in a particular configuration. Also, the use of a more stringent two-point threshold *η* = 5, improved the performance of the phasing algorithm.

**Table 2.**
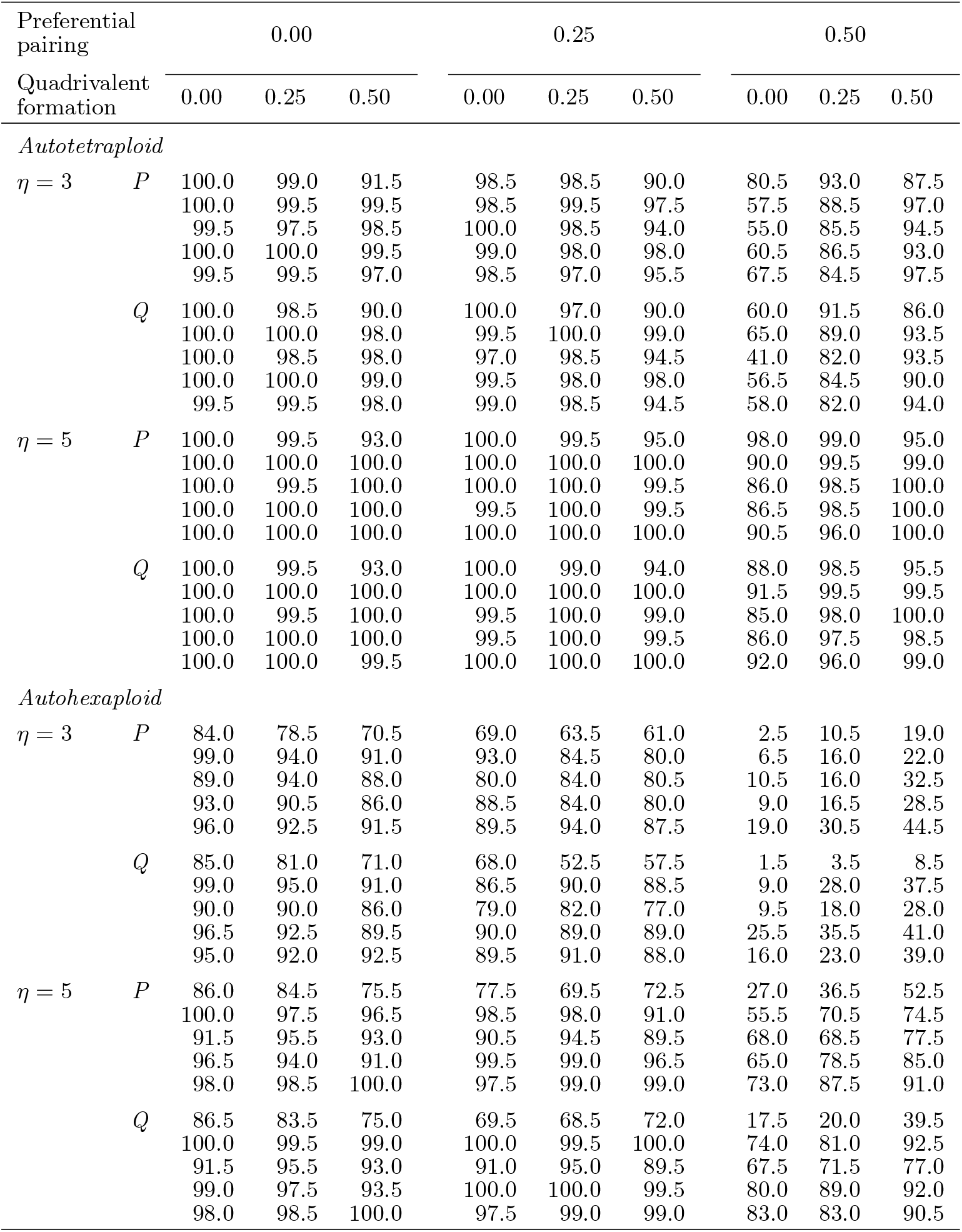
Percentage of data sets where linkage phase configuration was correctly estimated for parents *P* and *Q* in simulation 2.

Within the preferential pairing level 0.25, results showed decay of correctly estimated linkage phases, which was more pronounced for hexaploid cases with threshold level *η* = 3, reaching a minimum value of 52.5% for parent Q in configuration 1. Again, the use of a higher two-point threshold level, *η* = 5, helped to improve this number to 68.5%. For preferential pairing level 0.50, there was a clear distinction between the results in tetraploid and hexaploid cases. In the former, the effect was not as pronounced as it was in the latter, where in several cases, the proportion of correctly estimated linkage phases was close to zero. As expected, the usage of a higher threshold level of n = 5 helped to improve the number of corrected estimated linkage phase configurations. Interestingly, for both cases with preferential pairing (0.25 and 0.50), the formation of quadrivalents had an overall tendency to improve the algorithm’s performance. This improvement was expected because when a quadrivalent is formed, each chromosome involved can exchange segments with two others, providing more information regarding their phase configuration.

Given a correctly estimated linkage phase, the recombination fractions were consistently estimated for all levels of preferential pairing with no quadrivalent formation (Fig. 6). However, they were overestimated in the presence of quadrivalent formation. This effect was mainly observed at the terminal regions of the chromosome, especially in the long arm, where double reduction is more pronounced. In this case, tetraploid maps were the most affected. This is in agreement with our expectations since in autohexaploid simulations, there was always the formation of a bivalent which was not involved in the double reduction process (although the rates of double reduction were very similar in both ploidy levels, Fig. 6). In addition to the quadrivalent, the bivalent serves as an extra source of information to access the recombination events. The average Euclidean distances reflect the overestimation of recombination fractions in cases with quadrivalent formation, showing distributions with higher medians and interquartile ranges in tetraploid cases when compared to hexaploids (Supplementary Information, Figure S6). Nevertheless, all the Euclidean distances distributions were located relatively close to zero, with a maximum value of 1.41 cM, indicating that although we observed overestimated recombination fractions towards the terminal ends of the chromosome, they were equally distributed, causing no severe disturbances in the final map. Figure S7, in Supplementary Information, shows an example of the effect of increasing quadrivalent formation rate in autotetraploid and autohexaploid maps. As the markers get further away from the centromere, the recombination fractions become overestimated.

**Figure 6.**
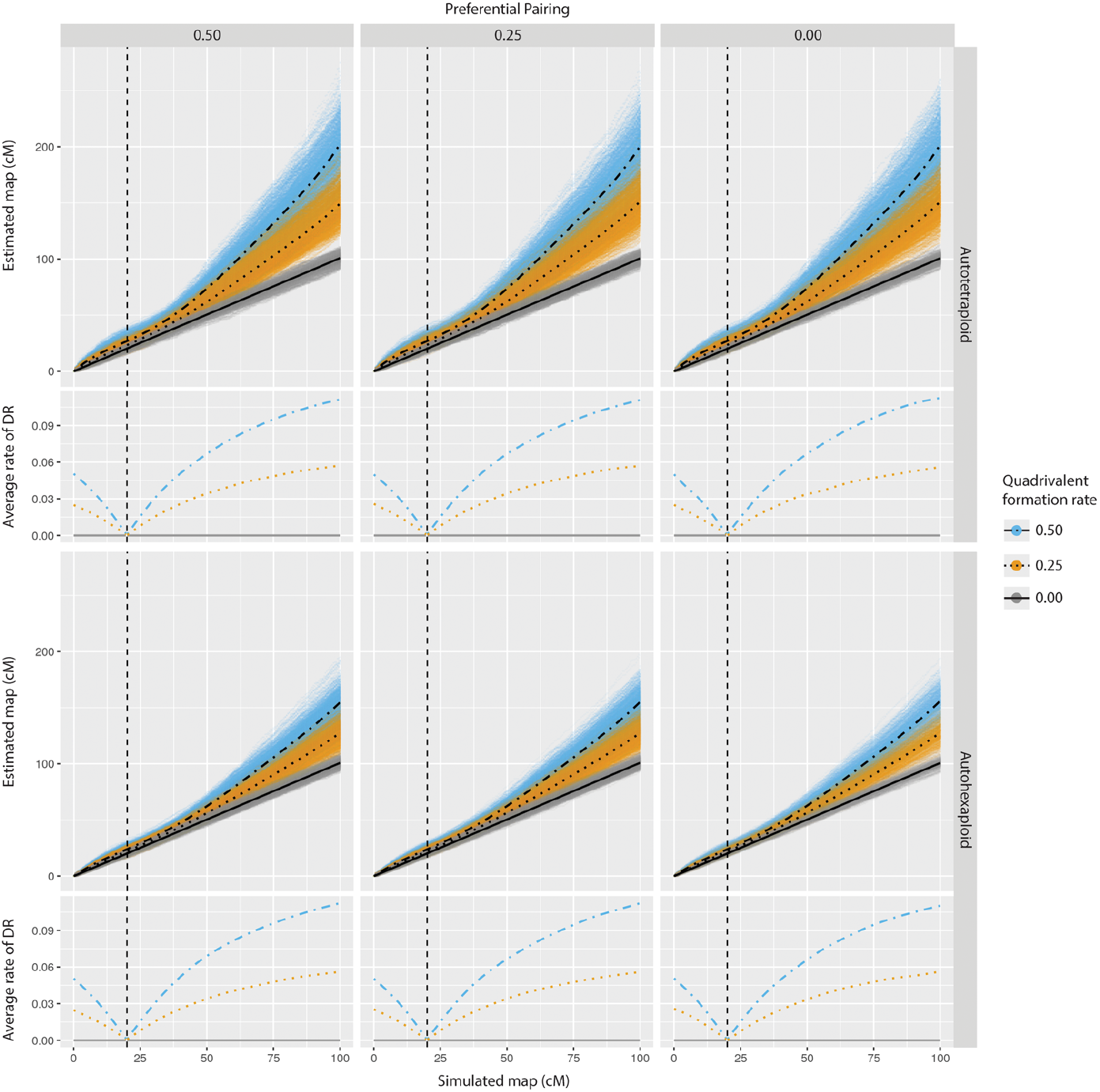
Comparison of estimated versus simulated maps given a correct estimation of linkage phases in simulation 2. Smoothed conditional means of the observed average rate of double reduction is presented along with the simulated chromosome. The centromere was positioned at 20 cM from its beginning (vertical dashed line). Upper panels show the results for tetraploid simulations while lower panels show the results for hexaploid simulations. Three levels of preferential pairing (0.00, 0.25, 0.50) and three levels of quadrivalent formation rate (0.00, 0.25, 0.50) were simulated. The lines superimposed to the scatter plots are smoothed conditional means of the distances using a generalized additive model. Both two-point thresholds were considered since they only affect the phasing procedure.

## Analysis of real tetraploid potato SNP data set

We applied our method to construct a genetic map of the B2721 population which is a cross between tetraploid potato varieties *Atlantic* x *B1829-5.* The population comprises 160 offsprings genotyped with the SolCAP Infinium 8303 potato array. The genotype calling was performed using fitTetra R package [56]. We obtained 4017 SNPs and computed all the pairwise recombination fraction between them for all possible linkage phase configurations. For each pair, we selected the configuration that yields the higher likelihood and applied the Unweighted Pair Group Method with Arithmetic Mean (UPGMA) clustering algorithm to assign markers into 12 linkage groups. Within each linkage group, we ordered the markers using the MDSMap R package [41]. We applied the unconstrained multidimensional scaling algorithm (MDS) to the pairwise marker distance obtained using Haldane’s mapping function and *LOD*^2^ weighting. We performed two rounds of marker removal by inspecting the nearest neighbor measure scatter plot and also the monotony of the resulting recombination fraction matrix. Given the marker order obtained for each group, we applied our algorithm using η =10. For each round of marker inclusion, we limited our phase search to the last 50 markers inserted at the end of the map and eliminated markers that caused map inflation greater than 10 cM. The resulting map consisted of 3374 SNPs (58% simplex and double simplex markers and 42% multiplex) distributed in 12 linkage groups with lengths varying from 181.4 cM to 390.9 cM with no visible gaps between markers (Supplementary Information, Table S8.1). These values seem inflated when compared to tetraploid potato maps available in the literature [6,21,33,43,46]. The map expansion observed here was mainly caused by two sources of error, namely, local marker misplacement and genotyping errors. Both factors cause the detection of spurious recombination events which are propagated through the HMM causing a global overestimation of recombination fractions.

Since we have the likelihood as an objective function to compare alternative orders, we reestimated the map using the genomic order obtained from the *Solanum tuberosum* genome version 4.03 [46]. When using the genomic order, the length of all linkage groups are smaller, and the likelihoods were substantially superior when compared to the *de novo* MDS-based order (Supplementary Information, Table S8.1 and Figure S8.1). Furthermore, our algorithm estimated the same linkage phase in both cases, indicating the robustness of the phasing method to local marker misplacement. To address the genotyping errors, we used the approaches presented in Eqs 8 and 9, i.e., (*i*) use the probability distribution provided by the mixture proportions of the doses from fitTetra software and (*ii*) assuming a global genotyping error; in this case we assumed 5%. We applied this prior information in both *de novo* MDS-based and the genome-based orders. The result also can be observed in Table S8.1 and Figure S8.1. Both approaches produced smaller maps when compared to their relative original maps. However, since (*i*) relies on the dosage proportions based in single SNPs, the adjustment of the map was not as flexible as observed in (*ii*), which assumes an equal global error for all markers. Thus the usage of global error, allowed genotypes to conform to a global chromosomal structure, rather than restrain the markers in certain genotypic classes proposed by the classification method. Furthermore, the usage of a global error mitigates the effect of the local markers misplacement caused by the MDS algorithm by clustering markers into linkage disequilibrium blocks. This effect can be observed in linkage groups 1, 5, 8, 9 and 12, where the difference between *de novo* MDS-based and the genome-based order when modeling a global error was less than 10 cM.

## Comparison with polymapR software

Among the available methods to construct maps in high-dose autopolyploids, namely, pergola [17], netgaws [3], and polymapR [5], only the latter is capable of inferring parental haplotypes and estimating recombination fraction in outcrossing populations. Thus, we limited our comparison to polymapR software. To assess the performance of the methods, we simulated 30 full-sib hexaploid populations with 200 each individuals in five marker density scenarios: 200, 400, 600, 800, and 1000 equally spaced markers in a 100 cM linkage group. Similarly to Simulation 2 described before, we randomly assigned allelic variants, from 0 to 3 doses, to the six homologous chromosomes. Additionally, in this study, we considered two different dosage proportions scenarios: the first one considers a higher proportion of simplex and double simplex markers, with 40% of the simulated markers being nulliplex, 40% simplex, 10% duplex and 10% triplex in both parents; the second, considers equal proportions for all doses, with 25% for all dosage types, from nulliplex to triplex (Supplementary Information, Figure S9). In total, 300 populations were simulated (5 × 2 × 30) and for each, we obtained the phased map using polymapR and our HMM-based procedure. All simulations were performed using the software PedigreeSim [57] considering no preferential pairing and no quadrivalent formation. For both methods, we recorded the percentage of correctly estimated linkage phases, the final map length, and the number of markers inserted in each map.

To employ our HMM-based method, we ordered the markers using the unconstrained MDS algorithm [41] weighted by *LOD*^2^ with no rounds of marker removal. Given the MDS order, two levels of *η* were used: 3 and 5; the same levels were used for the multilocus LOD threshold. The phase search was limited to the last 50 markers inserted at the end of the map. Also, we eliminated markers that caused map expansion higher than 5 cM. To construct maps using polymapR, we first applied the function `cluster_SN_markers` to perform a grid search from LOD Scores 1 to 20 and chose the lowest one that yields six homologous chromosomes based on simplex markers in coupling linkage phase. Subsequently, we connected the homologous based on the linkage between simplex × duplex markers using the function `bridgeHomologues`. In the next step, we assigned double simplex and duplex markers to the linkage group using the function `assign_linkage_group`, and the remaining marker types were assigned using the function `homologue_lg_assignment`. This procedure was performed for both parents assuming LOD Score thresholds of 3 and 5. All remaining pairwise recombination fractions were computed, and the MDS algorithm [41] was used to order the markers. Marker positions were estimated using the projection of the MDS result onto a single dimension principal curve. Finally, a phased map was created using the function `create_phased_maplist`. Differently from the HMM-based method, where the *η* indicates the LOD threshold from which the multipoint likelihood should be used to chose the best phase configuration, polymapR uses LOD thresholds to make decisions whether a marker should be used in a certain mapping context, such as, clustering homologous chromosomes or to assign it into assembled linkage groups. Thus, they are not directly comparable.

Table S10.1, in Supplementary Information, shows the results obtained using both methods. Overall, in the presence of a high number of single-dose markers, both approaches performed well. On average, polymapR was able to estimate 97.65% of the phases correctly and to position 89.56% of markers. Our HMM-based method correctly estimated 99.87% phases and positioned 99.96% of markers. In the cases where the dosages were uniformly assigned, our method outperformed polymapR. For 200 and 400 marker cases, polymapR was not able to estimate any correct linkage phase for both LOD thresholds and for 600 markers case, it obtained 55.85% of correct phases, on average. For 800 and 1000 markers, polymapR phased all markers correctly. Nevertheless, the number of positioned markers was quite low in those cases, varying from 24.6% to 77.4% for LOD = 3 and from 20.2% to 56.9% for LOD = 5. As pointed out by [53], when using polymapR’s method, the presence of a sufficiently large number of simplex markers uniformly distributed throughout the genome is essential to define each homologous chromosomes. On the other hand, our method showed to be capable of dealing with the absence of single-dose markers, especially under the more stringent LOD threshold 5, where, except from one parent in one simulation, it estimated all linkage phase configuration correctly, including in average 99.3% of markers in the phased map.

Table S10.3, Supplementary Information, shows the average map length and the associated standard deviation obtained in all simulations. In cases with a high number of single-dose markers, the average map length produced by polymapR ranged from 91.4 to 87.3 cM, while in the case where the dosages where uniformly assigned, map lengths ranged from 79.7 and 79.5 cM. These results confirm the underestimation tendency in the MDS algorithm when using when using *LOD*^2^ as weighting function, as observed by [41]. In our HMM-based method, considering the MDS order, the maps lengths were highly overestimated, ranging from 143.0 cM to 548.0 cM. Since we did not introduce errors in our simulation procedures, the observed map inflation is exclusively due to local marker misplacement caused by the MDS algorithm. Nonetheless, even with local marker misplacements, the linkage phase configuration was correctly estimated in the vast majority of the cases. To accommodate the local marker misplacements, we used a global error of 5% in the HMM emission function. As observed in the tetraploid potato analysis, this strategy allowed markers in the wrong order, but in the same linkage disequilibrium block, being positioned closely together through the HMM estimation process. The resulting maps lengths were close to the simulated 100 cM (Supplementary Information, Table S10.3 and Figure S10.1). Although in general, our method achieved a superior performance, it is worthwhile to mention that, polymapR’s method is substantially faster than ours, notably when our method uses high values for η, in which case the HMM computations play a significant role in the phasing procedures. Nonetheless, it was precisely the multipoint procedure that allowed our method to resolve situations where polymapR was not able to recover the simulated map.

## Availability of Data and Materials

All the methods and procedures described here are available in the R package *MAPpoly*, which is freely available from https://github.com/mmollina/mappoly. R scripts to perform the simulations and the potato map construction presented in this article can be accessed at https://github.com/mmollina/Autopolyploid_Linkage. The tetraploid potato data set is available though the Solanaceae Coordinated Agricultural Project at http://solcap.msu.edu/potato_infinium.shtml. Supplemental material available at Figshare

## Discussion

Although the concept of linkage mapping is relatively simple, the combinatorial properties and increasingly missing information that arise from the multiple sets of chromosomes make the construction of genetic maps in high-level autopolyploids challenging. In this work, we frame and solve two fundamental steps towards the construction of such maps, namely multipoint recombination fraction estimation and linkage phase estimation. We showed that, combined with standard grouping and ordering procedures [41], these maps could be reliably constructed. Our method can be applied to biallelic codominant markers and, due to the flexibility of the HMM framework upon which it was derived, it is extendable to any codominant molecular marker. The HMM used in this work takes into account the linkage phase configuration of the whole linkage group to estimate the recombination fractions between adjacent markers. An efficient two-point approach was also presented to reduce the search space of linkage phase configurations. As a result, our method provides the likelihood of the model, which can be used as an objective function to compare different map configurations, including linkage phases and marker orders. When considering experimental populations, our method is a generalization, for any even ploidy level, of well established genetic linkage mapping methods. For diploid (*m* = 2) populations derived from biparental crosses, our method is equivalent to the influential Lander and Green algorithm [25]; considering full-sib phase-unknown crosses, it is equivalent to [62]. For tetraploids (*m* = 4) the method is equivalent to [27], disregarding double reduction.

To assess the statistical power of our method, we conducted two simulation studies. In simulation 1, we demonstrated that our model was capable of correctly estimating the majority of parental linkage phase configurations and recombination fractions in a limited number of markers, even for complex linkage phase configurations and high ploidy levels. Since other methods are based on single-dose markers to assemble homology groups, to the best of our knowledge, this is the only method capable of phasing markers in high-dose autopolyploid genomes in small regions. These well-assembled regions could function as multiallelic codominant markers which propagate their information through the HMM to the rest of the chain, improving the quality of the final map. In simulation 2, we analyzed a sequence of 200 markers in combinations of different levels of preferential pairing and rates of quadrivalent formation. In this situation, quadrivalent formation rate had a marginal effect on the phasing procedure, whereas preferential pairing reduced its performance, especially for autohexaploids. The usage of a higher two-point threshold (*η*) improved the linkage phase estimation in all cases. This fact indicates that the haplotype phasing is more accurate when HMM-based likelihood is used as objective function to evaluate linkage phases. We also observed that quadrivalent formation yield overestimated recombination fractions between adjacent markers located further away from the centromere. This behavior was expected since our model disregards double reduction and, consequently, was not able to correctly estimate the number of crossing over events when this phenomenon was present. Although our model is robust enough to cope with certain levels of preferential pairing and tetravalent rate formation, it is possible to include both phenomena in specific points of its derivation. Preferential paring can be included in Eq 4 by not considering Pr(ψ¿) as uniformly distributed. Double reduction can be included in the definition of the genotypic states in the full transition space (Eq 5). These two phenomena add extra layers of complexity to the genetic mapping of polyploid organisms with high ploidy levels and should be addressed in future studies.

We also build a tetraploid map using the ideas presented in this study coupled with standard grouping and ordering procedures. We demonstrate that it is possible to use prior information on the HMM framework, including the probability distribution of the marker dosages for each SNP and a global error to avoid map inflation caused by local marker misplacement and genotyping errors. Finally, we compared our HMM-based method to the polimapR two-point based method and, as already pointed out by [53], we concluded that with a number sufficiently large of single-dose markers uniformly distributed across the homologous chromosomes, both methods performed well. However, when those markers are absent in a specific homologous or chromosome region, our method over-performed polymapR’s. Moreover, some autopolyploids with ploidy level higher than six, such as sugarcane, could benefit only from our method, since polymapR is limited only to tetraploid and hexaploid species. The difficulty in correctly estimating entangled linkage phase configurations lies in two significant aspects of the experiments studied here: (i) the outbred nature of the experimental crosses and (ii) the incomplete information of the markers based on dosage (i.e., by not being multiallelic). In experimental population derived from inbred lines, the origin of the haplotypes can be easily inferred from the genetic design. However, obtaining pure inbred lines in high-level autopolyploids has been proven to be impractical due to the high number of crosses and generations necessary to achieve homozygous genotypes and to the inbreeding depression which some species undergo [15]. In our method, the linkage phase configuration is obtained by comparing the likelihood of a set of models with different linkage phase configurations (Eq 18). The capability of estimating the correct configuration is directly related to the information contained in the marker data. Some of these limitations are overcome through the use of HMMs which take into account the information of the linkage group as a whole.

HMMs provide an excellent avenue to assemble genetic maps in complex scenarios, but they are remarkably computational demanding and, in some cases, unfeasible to use. Apart from parallel computing, which can greatly speed up the estimation process and is ubiquitous nowadays, the usage of two-point approaches is a viable option to reduce the dimension of the original problem efficiently. The dimension reduction is achieved by collapsing genotypic states in the full transition space according to the marker information. However, in several cases, the two-point based method can result in low statistical power which is related to the amount of information contained in markers in certain combinations of allelic dosage and linkage phase configurations. This lack of information is exacerbated as markers get distant from each other. Fig 7 shows nine possible configurations of pairs of markers in one autohexaploid parent. Considering one of the parents non-informative, we computed the Fisher’s information equations based on the likelihood Eq (22) [32, 35, 44]. The equations were plotted as a function of the recombination fraction. The information profiles are related to the number of different haplotypes present on the parental configuration for a given marker dosage. For instance, for two single-dose markers (Fig 7, panel I), when the alleles share the same homologous chromosome (*w_k_* = 1), it is always possible to detect if the gamete contains at least one recombinant chromosome. However, when the alleles are in different homologous chromosomes (*w_k_* = 0), the detection of recombination events is limited to meiotic configurations containing a bivalent where these chromosomes paired with each other. Intermediate situations involving multi-dose markers can be observed in the other panels in Fig 7. Additionally, the model proposed here contemplates both parents on the analyses, leading to more complicated linkage phase configurations and information equations.

**Figure 7.**
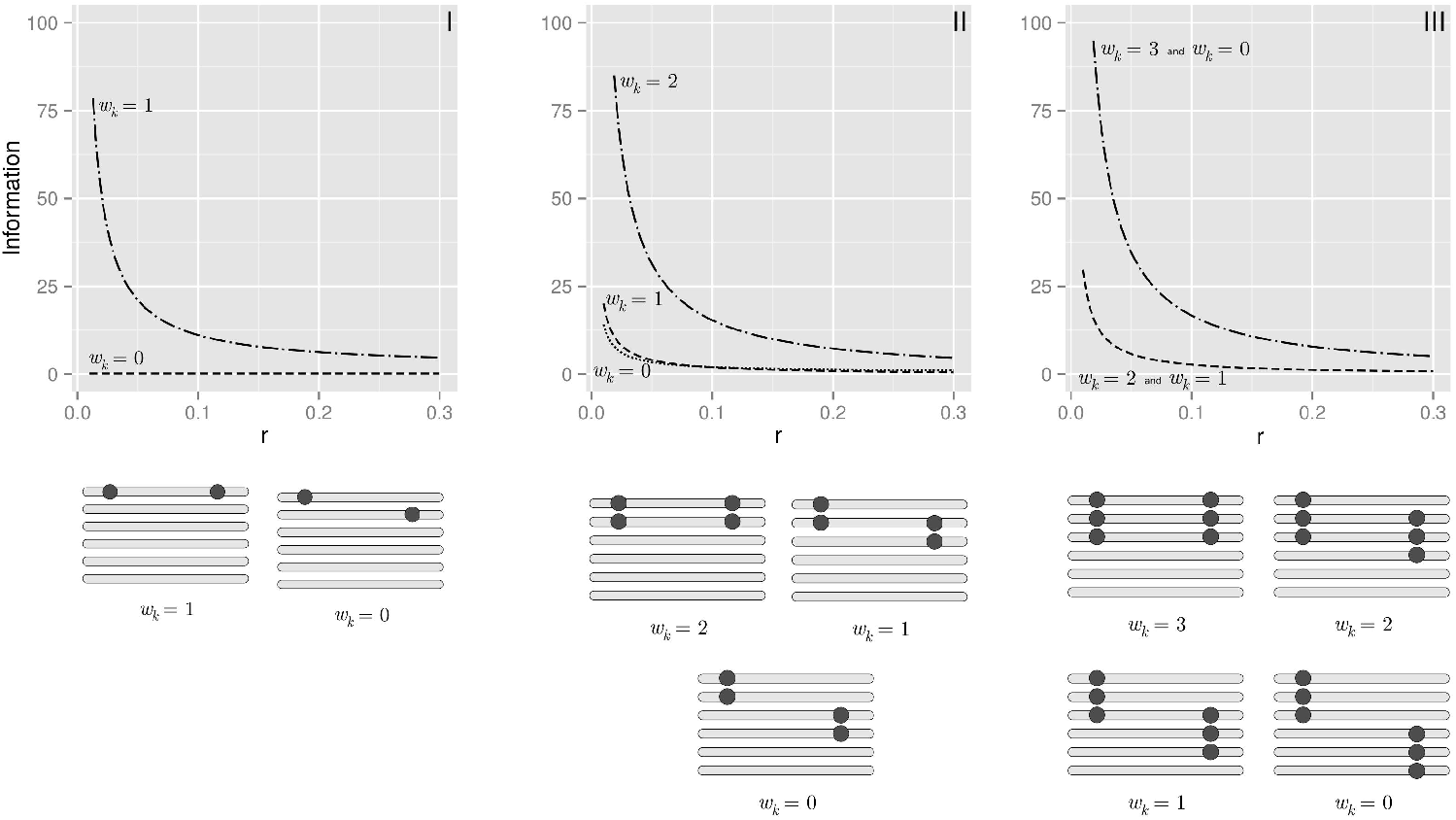
Fisher’s information for the two-point maximum likelihood estimators in different combinations of dosages and linkage phases configurations considering one informative hexaploid parent. (I) single-dose markers; alleles share 1 and 0 homologous. (II) double-dose markers; alleles share 2, 1 and 0 homologous. (III) triple dose markers; alleles share 3, 2, 1 and 0 homologous.

The lack of information for some phase configurations in two-point procedures is essentially caused by the biallelic nature of the dosage-based markers. However, in several situations, genomic and transcriptomic references are available for related diploid species and often provide the physical order of the SNPs in small regions. Our phasing procedure could be applied in these regions to obtain local haplotypes, which could function as multiallelic markers improving the information in a two-point analysis. Moreover, in a multipoint context, when using multiallelic markers, the number of visited states in the Markov model can be significantly reduced, making the HMM procedure much more efficient. Ideally, in a full-sib population, the number of different alleles should be as high as two times the ploidy level (fully informative). In this case, the Markov model would be fully observed and, the task of estimating recombination fraction reduces to count the number of recombinant events given a linkage phase configuration. Since our algorithm does not need the entire transition space to work, only a subset of states should be visited, making the calculation much faster when compared to the biallelic case.

Once the map is assembled, given the HMM framework, it is a trivial exercise to obtain the probability of a specific genotype at any map position, conditioned on the whole linkage group and to compute the probability of any unobserved genotype given the genetic map using this information. These conditional probabilities are the basis for answering a series of fundamental questions about quantitative trait loci analysis in high-level autopolyploids, such as the effect of the dosage level on the variation of quantitative traits, the interaction of the alleles within (dominance effects) and between loci (epistatic effects). Therefore, the present study provides a sound basis for unveiling the complex structure of autopolyploid genomes through genetic mapping.

## Acknowledgments

The authors wish to thank Dr. Guilherme da Silva Pereira and Dr. Zhao-Bang Zeng for their invaluable suggestions for elaboration of the manuscript. This work was supported by the Bill and Melinda Gates Foundation [OPP1052983] and is part of the Genomic Tools for Sweetpotato Improvement project (GT4SP); MM was also supported by Fundação de Amparo a Pesquisa do Estado de São Paulo (FAPESP) [2012/17009-8, 2013/12245-8]; AAFG was supported by FAPESP [08/52197-4]; AAFG has a scholarship from CNPq.

