## Supplementary material for "Linkage analysis and haplotype phasing in experimental autopolyploid populations with high ploidy level using hidden Markov models": File S1

### File S1: Algebraic simplifications for transition probabilities

Given a vector $\mathbf{p}_k$, the number of possible vectors $\mathbf{p}_{k+1}$ which have $l$ recombinant chromosomes between $k$ and $k+1$ is $\binom{\frac{m}{2}}{l}$; the number of possible $\mathbf{p}_{k+1}$ which have $\frac{m}{2}-l$ parental chromosomes is $\binom{\frac{m}{2}}{\frac{m}{2}-l}$. Thus for a given $\mathbf{p}_k$ and $l$ , the number of possibles $\mathbf{p}_{k+1}$ (i.e. all possible states in locus $k+1$ with $l$ recombinants) is

$$\binom{\frac{m}{2}}{l}\binom{\frac{m}{2}}{\frac{m}{2}-l} = \binom{\frac{m}{2}}{l}^2$$

Thus, the probability of $\mathbf{p}_k$ is

$$\Pr(\mathbf{p}_k) = \sum_{l=0}^{\frac{m}{2}} \Pr(\mathbf{p}_k,\ \mathbf{p}_{k+1})\binom{\frac{m}{2}}{l}^2 \tag{1}$$

Using (1),

$$\Pr(\mathbf{p}_{k+1}|\mathbf{p}_k) = \frac{\Pr(\mathbf{p}_k,\ \mathbf{p}_{k+1})}{\sum_{l=0}^{\frac{m}{2}} \Pr(\mathbf{p}_k,\ \mathbf{p}_{k+1})\binom{\frac{m}{2}}{l}^2} \tag{2}$$

The numerator of (2) can be written as

$$\begin{aligned}
\Pr(\mathbf{p}_k,\ \mathbf{p}_{k+1}) &= \frac{\frac{(1-r_k)^{\frac{m}{2}-l}(r_k)^l}{2^{\frac{m}{2}}} l!(\frac{m}{2}-l)!}{\frac{1}{\frac{m}{2}!}\prod_{i=2,4,\cdots,m}\binom{i}{2}} \\
&= (1-r_k)^{\frac{m}{2}-l}(r_k)^l \frac{l!(\frac{m}{2}-l)!\frac{m}{2}!}{2^{\frac{m}{2}}\prod_{i=2,4,\cdots,m}\binom{i}{2}} \qquad (3)
\end{aligned}$$

The denominator of (2) can be written as

$$
\begin{aligned}
\sum_{l=0}^{\frac{m}{2}} \Pr(\mathbf{p}_k,\ \mathbf{p}_{k+1}) \binom{\frac{m}{2}}{l}^2 &= \sum_{l=0}^{\frac{m}{2}} \binom{\frac{m}{2}}{l}^2 \frac{\frac{(1-r_k)^{\frac{m}{2}-l}(r_k)^l}{2^{\frac{m}{2}}} l!(\frac{m}{2}-l)!}{\frac{1}{\frac{m}{2}!} \prod_{i=2,4,\cdots,m} \binom{i}{2}} \\
&= \sum_{l=0}^{\frac{m}{2}} \binom{\frac{m}{2}}{l} (1-r_k)^{\frac{m}{2}-l}(r_k)^l \frac{l!(\frac{m}{2}-l)!\frac{m}{2}!}{2^{\frac{m}{2}} \prod_{i=2,4,\cdots,m} \binom{i}{2}} \frac{\frac{m}{2}!}{l!(\frac{m}{2}-l)!} \\
&= \frac{\frac{m}{2}!^2}{2^{\frac{m}{2}} \prod_{i=2,4,\cdots,m} \binom{i}{2}} \sum_{l=0}^{\frac{m}{2}} \binom{\frac{m}{2}}{l} (1-r_k)^{\frac{m}{2}-l}(r_k)^l \\
&= \frac{\frac{m}{2}!^2}{2^{\frac{m}{2}} \prod_{i=2,4,\cdots,m} \binom{i}{2}} \left\{(1-r_k)r_k\right\}^{\frac{m}{2}} \\
&= \frac{\frac{m}{2}!^2}{2^{\frac{m}{2}} \prod_{i=2,4,\cdots,m} \binom{i}{2}} \qquad (4)
\end{aligned}
$$

Dividing (3) by (4),

$$
\begin{aligned}
\Pr(\mathbf{p}_{k+1}|\mathbf{p}_k) &= (1-r_k)^{\frac{m}{2}-l}(r_k)^l \frac{l!(\frac{m}{2}-l)!}{\frac{m}{2}!} \\
&= \frac{(1-r_k)^{\frac{m}{2}-l}(r_k)^l}{\binom{\frac{m}{2}}{l}} \qquad (5)
\end{aligned}
$$
