## Supplementary material for "Linkage analysis and haplotype phasing in experimental autopolyploid populations with high ploidy level using hidden Markov models": File S2

### File S2: Algorithm for obtaining $l_P$ and $l_Q$ given two genotypic indices

This algorithm is used to find the number of recombinant bivalents between loci $k$ and $k+1$ for parents $P$ and $Q$ (i.e. $l_P$ and $l_Q$) given a ploidy level and the two genotypic indices $j$ and $j'$ which indicate the order of the genotypic states in the transition space in the full-sib population, defined in Eq 6. Consider a binary vector $\mathbf{a}_i$ of length $m$ indicating the presence and absence of alleles in the gametic genotypes $\theta^m_{P,i}$ or $\theta^m_{Q,i}$. See Table S2.1 for an example of vectors in an autohexaploid with their gametic genotypes and indices. With all possible vectors listed, the number of recombinant bivalents $l_P$ between any two gametic genotype, $\theta^m_{P,i}$ and $\theta^m_{P,i'}$ is $\frac{1}{2}\sum \mid \mathbf{a}_i - \mathbf{a}_{i'} \mid$. In real situations, for high ploidy levels the enumeration of all possible $\mathbf{a}$ vectors is a highly intensive task. Thus, Algorithm 1 returns the binary vector for any ploidy level $m$ in any $i$ position in the transition space. This procedure holds for both parents and can be applied to obtain $l_P$ and $l_Q$ given $j$ and $j'$. To obtain $l_P$, consider $i = 1 + (j-1)\ \mathbf{div}\ \binom{m}{\frac{m}{2}}$ and $i' = 1 + (j'-1)\ \mathbf{div}\ \binom{m}{\frac{m}{2}}$, where **div** denotes the integer division operator. Then, $\mathbf{a}_i$ and $\mathbf{a}_{i'}$ can be obtained using Algorithm 1 and $l_P = \frac{1}{2}\sum \mid \mathbf{a}_i - \mathbf{a}_{i'} \mid$. To obtain $l_Q$, consider $h = \binom{m}{\frac{m}{2}} + j - i\binom{m}{\frac{m}{2}}$ and $h' = \binom{m}{\frac{m}{2}} + j' - i'\binom{m}{\frac{m}{2}}$. Then, $l_Q = \frac{1}{2}\sum \mid \mathbf{a}_h - \mathbf{a}_{h'} \mid$.

Table S2.1 : **Example of binary vectors in an autohexaploid with their genotypes and indices.**

| $i$ | Gametic genotype ($\theta^m_{P,i}$) | $\mathbf{a}_i$ |
|---|---|---|
| 1 | $\{P^1_k, P^2_k, P^3_k\}$ | $\{1,1,1,0,0,0\}$ |
| 2 | $\{P^1_k, P^2_k, P^4_k\}$ | $\{1,1,0,1,0,0\}$ |
| 3 | $\{P^1_k, P^2_k, P^5_k\}$ | $\{1,1,0,0,1,0\}$ |
| 4 | $\{P^1_k, P^2_k, P^6_k\}$ | $\{1,1,0,0,0,1\}$ |
| 5 | $\{P^1_k, P^3_k, P^4_k\}$ | $\{1,0,1,1,0,0\}$ |
| 6 | $\{P^1_k, P^3_k, P^5_k\}$ | $\{1,0,1,0,1,0\}$ |
| 7 | $\{P^1_k, P^3_k, P^6_k\}$ | $\{1,0,1,0,0,1\}$ |
| 8 | $\{P^1_k, P^4_k, P^5_k\}$ | $\{1,0,0,1,1,0\}$ |
| 9 | $\{P^1_k, P^4_k, P^6_k\}$ | $\{1,0,0,1,0,1\}$ |
| 10 | $\{P^1_k, P^5_k, P^6_k\}$ | $\{1,0,0,0,1,1\}$ |
| 11 | $\{P^2_k, P^3_k, P^4_k\}$ | $\{0,1,1,1,0,0\}$ |
| 12 | $\{P^2_k, P^3_k, P^5_k\}$ | $\{0,1,1,0,1,0\}$ |
| 13 | $\{P^2_k, P^3_k, P^6_k\}$ | $\{0,1,1,0,0,1\}$ |
| 14 | $\{P^2_k, P^4_k, P^5_k\}$ | $\{0,1,0,1,1,0\}$ |
| 15 | $\{P^2_k, P^4_k, P^6_k\}$ | $\{0,1,0,1,0,1\}$ |
| 16 | $\{P^2_k, P^5_k, P^6_k\}$ | $\{0,1,0,0,1,1\}$ |
| 17 | $\{P^3_k, P^4_k, P^5_k\}$ | $\{0,0,1,1,1,0\}$ |
| 18 | $\{P^3_k, P^4_k, P^6_k\}$ | $\{0,0,1,1,0,1\}$ |
| 19 | $\{P^3_k, P^5_k, P^6_k\}$ | $\{0,0,1,0,1,1\}$ |
| 20 | $\{P^4_k, P^5_k, P^6_k\}$ | $\{0,0,0,1,1,1\}$ |

**Algorithm 1**

**function** BOLVEC($m, i$)<br>
    $s_0 \leftarrow 0$<br>
    $s_1 \leftarrow 1$<br>
    $increment \leftarrow 0$<br>
    $sentinel \leftarrow 0$<br>
    vector $a(m)$ ▷ Vector **a** of size $m$<br>
    **while** $sentinel < \frac{m}{2}$ **do**<br>
        $temp \leftarrow \binom{m-s_1}{\frac{m}{2}-s_0-1}$<br>
        **if** $i > temp + increment$ **then**<br>
            $a(s_1 - 1) \leftarrow 0$<br>
            $increment \leftarrow increment + temp$<br>
        **else**<br>
            $a(s_1 - 1) \leftarrow 1$<br>
            $s_0 + +$<br>
        **end if**<br>
        $sentinel \leftarrow sentinel + a(s_1 - 1)$<br>
        $s_1 + +$<br>
    **end while**<br>
    **return** $a$<br>
**end function**
