## Supplementary material for "Linkage analysis and haplotype phasing in experimental autopolyploid populations with high ploidy level using hidden Markov models": File S3

### File S3: Example of usage of the two-point and multipoint procedures to infer the linkage phase configuration in both parents and estimate recombination fractions in a sequence of markers in high-level autopolyploids.

In order to show the mechanics of the mapping reconstruction using the combination of two-point and multipoint strategies, we present a simple full-bib autotetraploid mapping population example. This example is easily extendable to higher ploidy levels, since it does not involve matrix forms whose high dimensions would preclude the operations.

**Parent haplotypes and dataset simulation** The mapping population comprise 30 offsprings derived from two autotetraploid parents, $P$ and $Q$. We simulated one linkage group genotyped with three biallelic markers positioned at a fixed distance of 1cM between them. The linkage phase configuration of both parents is presented in Figure S3.1.

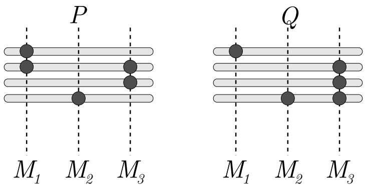


Figure S3.1 : Linkage phase configuration for three biallelic markers in two parents, $P$ and $Q$.

The observed doses in parent $P$ are $d_P^1 = 2$, $d_P^2 = 1$, $d_P^3 = 2$ and in $Q$ are $d_Q^1 = 1$, $d_Q^2 = 1$, $d_Q^3 = 3$. The simulated dataset of 30 offsprings is shown in Table S3.1. For simplicity purposes, we simulated the dosage information with no error, thus the probability distribution of the genotypes always have the value 1 associated to the simulated dosage and 0 to the remaining ones.

Table S3.1 : Simulated data set of 30 offsprings. Since the dosage informaton has no error, $O_{k,i}$ represents the dosage of the marker at position $k$ for individual $i$. We also present the vector of probabilities associated to the dosage genotypes $\boldsymbol{\pi}_{k,i}$ for marker $k$, individual $i$.

| Individual ($i$) | $O_{1,i}$ | $\boldsymbol{\pi}_{1,i}$ | $O_{2,i}$ | $\boldsymbol{\pi}_{2,i}$ | $O_{3,i}$ | $\boldsymbol{\pi}_{3,i}$ |
|---|---|---|---|---|---|---|
| 1 | 2 | $\{0,0,1,0,0\}$ | 0 | $\{1,0,0,0,0\}$ | 3 | $\{0,0,0,1,0\}$ |
| 2 | 1 | $\{0,1,0,0,0\}$ | 1 | $\{0,1,0,0,0\}$ | 2 | $\{0,0,1,0,0\}$ |
| 3 | 3 | $\{0,0,0,1,0\}$ | 0 | $\{1,0,0,0,0\}$ | 2 | $\{0,0,1,0,0\}$ |
| 4 | 1 | $\{0,1,0,0,0\}$ | 0 | $\{1,0,0,0,0\}$ | 3 | $\{0,0,0,1,0\}$ |
| 5 | 2 | $\{0,0,1,0,0\}$ | 1 | $\{0,1,0,0,0\}$ | 3 | $\{0,0,0,1,0\}$ |
| 6 | 1 | $\{0,1,0,0,0\}$ | 2 | $\{0,0,1,0,0\}$ | 3 | $\{0,0,0,1,0\}$ |
| 7 | 2 | $\{0,0,1,0,0\}$ | 1 | $\{0,1,0,0,0\}$ | 3 | $\{0,0,0,1,0\}$ |
| 8 | 2 | $\{0,0,1,0,0\}$ | 1 | $\{0,1,0,0,0\}$ | 3 | $\{0,0,0,1,0\}$ |
| 9 | 2 | $\{0,0,1,0,0\}$ | 0 | $\{1,0,0,0,0\}$ | 3 | $\{0,0,0,1,0\}$ |
| 10 | 3 | $\{0,0,0,1,0\}$ | 1 | $\{0,1,0,0,0\}$ | 2 | $\{0,0,1,0,0\}$ |
| 11 | 1 | $\{0,1,0,0,0\}$ | 1 | $\{0,1,0,0,0\}$ | 2 | $\{0,0,1,0,0\}$ |
| 12 | 2 | $\{0,0,1,0,0\}$ | 0 | $\{1,0,0,0,0\}$ | 2 | $\{0,0,1,0,0\}$ |
| 13 | 1 | $\{0,0,0,1,0\}$ | 1 | $\{0,1,0,0,0\}$ | 3 | $\{0,0,0,1,0\}$ |
| 14 | 2 | $\{0,0,1,0,0\}$ | 0 | $\{1,0,0,0,0\}$ | 2 | $\{0,0,1,0,0\}$ |
| 15 | 1 | $\{0,1,0,0,0\}$ | 2 | $\{0,0,1,0,0\}$ | 2 | $\{0,0,1,0,0\}$ |
| 16 | 1 | $\{0,1,0,0,0\}$ | 2 | $\{0,0,1,0,0\}$ | 2 | $\{0,0,1,0,0\}$ |
| 17 | 3 | $\{0,0,0,1,0\}$ | 1 | $\{0,1,0,0,0\}$ | 2 | $\{0,0,1,0,0\}$ |
| 18 | 2 | $\{0,0,1,0,0\}$ | 0 | $\{1,0,0,0,0\}$ | 3 | $\{0,0,0,1,0\}$ |
| 19 | 2 | $\{0,0,1,0,0\}$ | 1 | $\{0,1,0,0,0\}$ | 3 | $\{0,0,0,1,0\}$ |
| 20 | 2 | $\{0,0,1,0,0\}$ | 1 | $\{0,1,0,0,0\}$ | 3 | $\{0,0,0,1,0\}$ |
| 21 | 2 | $\{0,0,1,0,0\}$ | 0 | $\{1,0,0,0,0\}$ | 3 | $\{0,0,0,1,0\}$ |
| 22 | 3 | $\{0,0,0,1,0\}$ | 0 | $\{1,0,0,0,0\}$ | 2 | $\{0,0,1,0,0\}$ |
| 23 | 2 | $\{0,0,1,0,0\}$ | 0 | $\{1,0,0,0,0\}$ | 2 | $\{0,0,1,0,0\}$ |
| 24 | 2 | $\{0,0,1,0,0\}$ | 1 | $\{0,1,0,0,0\}$ | 1 | $\{0,1,0,0,0\}$ |
| 25 | 1 | $\{0,1,0,0,0\}$ | 1 | $\{0,1,0,0,0\}$ | 4 | $\{0,0,0,0,1\}$ |
| 26 | 3 | $\{0,0,0,1,0\}$ | 0 | $\{1,0,0,0,0\}$ | 2 | $\{0,0,1,0,0\}$ |
| 27 | 2 | $\{0,0,1,0,0\}$ | 1 | $\{0,1,0,0,0\}$ | 2 | $\{0,0,1,0,0\}$ |
| 28 | 2 | $\{0,0,1,0,0\}$ | 1 | $\{0,1,0,0,0\}$ | 2 | $\{0,0,1,0,0\}$ |
| 29 | 1 | $\{0,1,0,0,0\}$ | 2 | $\{0,0,1,0,0\}$ | 3 | $\{0,0,0,1,0\}$ |
| 30 | 1 | $\{0,1,0,0,0\}$ | 2 | $\{0,0,1,0,0\}$ | 2 | $\{0,0,1,0,0\}$ |

**Two-point recombination fraction** The efficient construction of the genetic map starts with the two-point analysis which allow a fast estimation of recombination fraction and linkage phase configuration between all possible pairs of markers in the dataset. The key point in this analysis is the reduction of the full transition space given a ploidy level, the dosage and linkage phase configuration of the markers in the parents. For each possible combination of dosage and linkage phase configuration, a specific reduction of dimension must be done. After that, for a given pair of markers with their specific doses, the best linkage phase configuration is assessed based on the likelihood of the evaluated models. Moreover, genetic linkage can also be tested.

Since $d_P^1 = 2$, $d_P^2 = 1$, $d_P^3 = 2$, $d_Q^1 = 1$, $d_Q^2 = 1$ and $d_Q^3 = 3$, we can write:

$$
\begin{aligned}
\phi_P^1 &= \{\{P_1^1, P_1^2\}, \{P_1^1, P_1^3\}, \{P_1^1, P_1^4\}, \{P_1^2, P_1^3\}, \{P_1^2, P_1^4\}, \{P_1^3, P_1^4\}\} \\
\phi_P^2 &= \{\{P_2^1\}, \{P_2^2\}, \{P_2^3\}, \{P_2^4\}\} \\
\phi_P^3 &= \{\{P_3^1, P_3^2\}, \{P_3^1, P_3^3\}, \{P_3^1, P_3^4\}, \{P_3^2, P_3^3\}, \{P_3^2, P_3^4\}, \{P_3^3, P_3^4\}\} \\
\phi_Q^1 &= \{\{Q_1^1\}, \{Q_1^2\}, \{Q_1^3\}, \{Q_1^4\}\} \\
\phi_Q^2 &= \{\{Q_2^1\}, \{Q_2^2\}, \{Q_2^3\}, \{Q_2^4\}\} \\
\phi_Q^3 &= \{\{Q_3^1, Q_3^2, Q_3^3\}, \{Q_3^1, Q_3^2, Q_3^4\}, \{Q_3^1, Q_3^3, Q_3^4\}, \{Q_3^2, Q_3^3, Q_4^4\}\}
\end{aligned}
$$

Table S3.2 shows all possible linkage phase configurations for markers $M_1$ and $M_2$

Table S3.2 : $\Phi_P^{1,2} = \phi_P^1 \times \phi_P^2$ and $\Phi_Q^{1,2} = \phi_Q^1 \times \phi_Q^2$ for the simulated data. In this case $d_P^1 = 2$, $d_P^2 = 1$, $d_Q^1 = 1$ and $d_Q^2 = 1$. In both cases there are two partitions. The gray cells represent the partition where $w_P^{1,2} = 1$ and the white cells represent the partition where $w_P^{1,2} = 0$

| | $\phi_P^2$ | | | |
|---|---|---|---|---|
| $\phi_P^1$ | $\{P_2^1\}$ | $\{P_2^2\}$ | $\{P_2^3\}$ | $\{P_2^4\}$ |
| $\{P_1^1, P_1^2\}$ | $\{\{P_1^1, P_1^2\}, \{P_2^1\}\}$ | $\{\{P_1^1, P_1^2\}, \{P_2^2\}\}$ | $\{\{P_1^1, P_1^2\}, \{P_2^3\}\}$ | $\{\{P_1^1, P_1^2\}, \{P_2^4\}\}$ |
| $\{P_1^1, P_1^3\}$ | $\{\{P_1^1, P_1^3\}, \{P_2^1\}\}$ | $\{\{P_1^1, P_1^3\}, \{P_2^2\}\}$ | $\{\{P_1^1, P_1^3\}, \{P_2^3\}\}$ | $\{\{P_1^1, P_1^3\}, \{P_2^4\}\}$ |
| $\{P_1^1, P_1^4\}$ | $\{\{P_1^1, P_1^4\}, \{P_2^1\}\}$ | $\{\{P_1^1, P_1^4\}, \{P_2^2\}\}$ | $\{\{P_1^1, P_1^4\}, \{P_2^3\}\}$ | $\{\{P_1^1, P_1^4\}, \{P_2^4\}\}$ |
| $\{P_1^4, P_1^3\}$ | $\{\{P_1^2, P_1^3\}, \{P_2^1\}\}$ | $\{\{P_1^2, P_1^3\}, \{P_2^2\}\}$ | $\{\{P_1^2, P_1^3\}, \{P_2^3\}\}$ | $\{\{P_1^2, P_1^3\}, \{P_2^4\}\}$ |
| $\{P_1^2, P_1^4\}$ | $\{\{P_1^2, P_1^4\}, \{P_2^1\}\}$ | $\{\{P_1^2, P_1^4\}, \{P_2^2\}\}$ | $\{\{P_1^2, P_1^4\}, \{P_2^3\}\}$ | $\{\{P_1^2, P_1^4\}, \{P_2^4\}\}$ |
| $\{P_1^3, P_1^4\}$ | $\{\{P_1^3, P_1^4\}, \{P_2^1\}\}$ | $\{\{P_1^3, P_1^4\}, \{P_2^2\}\}$ | $\{\{P_1^3, P_1^4\}, \{P_2^3\}\}$ | $\{\{P_1^3, P_1^4\}, \{P_2^4\}\}$ |

| | $\phi_Q^2$ | | | |
|---|---|---|---|---|
| $\phi_Q^1$ | $\{Q_2^1\}$ | $\{Q_2^2\}$ | $\{Q_2^3\}$ | $\{Q_2^4\}$ |
| $\{Q_1^1\}$ | $\{\{Q_1^1\}, \{Q_2^1\}\}$ | $\{\{Q_1^1\}, \{Q_2^2\}\}$ | $\{\{Q_1^1\}, \{Q_2^3\}\}$ | $\{\{Q_1^1\}, \{Q_2^4\}\}$ |
| $\{Q_1^2\}$ | $\{\{Q_1^2\}, \{Q_2^1\}\}$ | $\{\{Q_1^2\}, \{Q_2^2\}\}$ | $\{\{Q_1^2\}, \{Q_2^3\}\}$ | $\{\{Q_1^2\}, \{Q_2^4\}\}$ |
| $\{Q_1^3\}$ | $\{\{Q_1^3\}, \{Q_2^1\}\}$ | $\{\{Q_1^3\}, \{Q_2^2\}\}$ | $\{\{Q_1^3\}, \{Q_2^3\}\}$ | $\{\{Q_1^3\}, \{Q_2^4\}\}$ |
| $\{Q_1^4\}$ | $\{\{Q_1^4\}, \{Q_2^1\}\}$ | $\{\{Q_1^4\}, \{Q_2^2\}\}$ | $\{\{Q_1^4\}, \{Q_2^3\}\}$ | $\{\{Q_1^4\}, \{Q_2^4\}\}$ |

In order to show the mechanics of the dimension reduction, let us consider for $w_P^{1,2} = 0$ and $w_Q^{1,2} = 0$ the configurations $\{\varphi_P^1 = \{P_1^1, P_1^2\}, \varphi_P^2 = \{P_2^3\}\}$ and $\left\{\varphi_Q^1 = \{Q_1^1\}, \varphi_Q^2 = \{Q_2^2\}\right\}$. Table S3.8 shows the full transition space for autotetraploids. Notice that for higher ploidy levels, the dimension of the transition space can be unwieldy, but in real cases we do not need to represent it in its complete form. The reduction of dimensionality of the full transition space is based on the proper combination of the rows and columns in Table S3.8. As result, instead of a space with dimensions (36 × 36), the dimension of the new transition space is (5 × 5). As pointed out in Eq 20, each elements in the reduced transition space is composed by a combination of

$$\frac{(1 - r_k)^{m - l_P - l_Q} (r_k)^{l_P + l_Q}}{\binom{\frac{m}{2}}{l_P} \binom{\frac{m}{2}}{l_Q}}$$

with $l_P, l_Q = \{0, \cdots, m\}$. The weight of each element corresponding to the ordered pair $(l_P, l_Q)$ is given by the variable $\zeta_{D_1, D_2}(l_P, l_Q)$. Table S3.3 shows all possible $\zeta_{D_1, D_2}(l_P, l_Q)$ given $\varphi_P^1 = \{P_1^1, P_1^2\}$, $\varphi_Q^1 = \{Q_1^1\}$, $\varphi_P^2 = \{P_2^3\}$ and $\varphi_Q^2 = \{Q_2^2\}$ for all $(l_P, l_Q)$ for each dosage pair $(\mathcal{D}_1^4, \mathcal{D}_2^4)$.

| | | $(l_P, l_Q)$ | | | | | | | | |
|---|---|---|---|---|---|---|---|---|---|---|
| $\mathcal{D}_1^4$ | $\mathcal{D}_2^4$ | (0,0) | (0,1) | (0,2) | (1,0) | (1,1) | (1,2) | (2,0) | (2,1) | (2,2) |
| 0 | 0 | 0 | 0 | 0 | 1/18 | 1/3 | 1/9 | 1/36 | 1/6 | 1/18 |
| 0 | 1 | 1/36 | 1/6 | 1/18 | 1/6 | 2/3 | 1/6 | 1/18 | 1/6 | 1/36 |
| 0 | 2 | 1/18 | 1/6 | 1/36 | 1/9 | 1/3 | 1/18 | 0 | 0 | 0 |
| 0 | 3 | 0 | 0 | 0 | 0 | 0 | 0 | 0 | 0 | 0 |
| 0 | 4 | 0 | 0 | 0 | 0 | 0 | 0 | 0 | 0 | 0 |
| 1 | 0 | 1/18 | 1/3 | 1/9 | 1/3 | 5/3 | 1/2 | 1/9 | 1/2 | 5/36 |
| 1 | 1 | 2/9 | 5/6 | 7/36 | 5/6 | 10/3 | 5/6 | 7/36 | 5/6 | 2/9 |
| 1 | 2 | 5/36 | 1/2 | 1/9 | 1/2 | 5/3 | 1/3 | 1/9 | 1/3 | 1/18 |
| 1 | 3 | 0 | 0 | 0 | 0 | 0 | 0 | 0 | 0 | 0 |
| 1 | 4 | 0 | 0 | 0 | 0 | 0 | 0 | 0 | 0 | 0 |
| 2 | 0 | 5/36 | 1/2 | 1/9 | 1/2 | 5/3 | 1/3 | 1/9 | 1/3 | 1/18 |
| 2 | 1 | 2/9 | 5/6 | 7/36 | 5/6 | 10/3 | 5/6 | 7/36 | 5/6 | 2/9 |
| 2 | 2 | 1/18 | 1/3 | 1/9 | 1/3 | 5/3 | 1/2 | 1/9 | 1/2 | 5/36 |
| 2 | 3 | 0 | 0 | 0 | 0 | 0 | 0 | 0 | 0 | 0 |
| 2 | 4 | 0 | 0 | 0 | 0 | 0 | 0 | 0 | 0 | 0 |
| 3 | 0 | 1/18 | 1/6 | 1/36 | 1/9 | 1/3 | 1/18 | 0 | 0 | 0 |
| 3 | 1 | 1/36 | 1/6 | 1/18 | 1/6 | 2/3 | 1/6 | 1/18 | 1/6 | 1/36 |
| 3 | 2 | 0 | 0 | 0 | 1/18 | 1/3 | 1/9 | 1/36 | 1/6 | 1/18 |
| 3 | 3 | 0 | 0 | 0 | 0 | 0 | 0 | 0 | 0 | 0 |
| 3 | 4 | 0 | 0 | 0 | 0 | 0 | 0 | 0 | 0 | 0 |
| 4 | 0 | 0 | 0 | 0 | 0 | 0 | 0 | 0 | 0 | 0 |
| 4 | 1 | 0 | 0 | 0 | 0 | 0 | 0 | 0 | 0 | 0 |
| 4 | 2 | 0 | 0 | 0 | 0 | 0 | 0 | 0 | 0 | 0 |
| 4 | 3 | 0 | 0 | 0 | 0 | 0 | 0 | 0 | 0 | 0 |
| 4 | 4 | 0 | 0 | 0 | 0 | 0 | 0 | 0 | 0 | 0 |

Table S3.3 : $\zeta_{D_1,D_2}(l_P, l_Q)$ given $\varphi_P^1 = \{P_1^1, P_1^2\}$, $\varphi_Q^1 = \{Q_1^1\}$, $\varphi_P^2 = \{P_2^3\}$ and $\varphi_Q^2 = \{Q_2^2\}$. The results would be the same for any other combination of $\varphi$'s that result in $(w_P^{1,2} = 0, w_Q^{1,2} = 0)$.

Thus, $\mathbf{A}_{\varphi_P^1=\{P_1^1,P_1^2\},\varphi_P^2=\{P_2^3\},\varphi_Q^1=\{Q_1^1\},\varphi_Q^2=\{Q_2^2\}}(r_k)$ can be easily obtained using article's Eq. 21

$$\begin{bmatrix} \frac{r(1+r)}{36} & \frac{1+2r-2r^2}{36} & \frac{(r-2)(r-1)}{36} & 0 & 0 \\ \frac{2+4r-r^2}{36} & \frac{4-r+r^2}{18} & \frac{5-2r-r^2}{36} & 0 & 0 \\ \frac{5-2r-r^2}{36} & \frac{4-r+r^2}{18} & \frac{2+4r-r^2}{36} & 0 & 0 \\ \frac{(r-2)(r-1)}{36} & \frac{1+2r-2r^2}{36} & \frac{r(1+r)}{36} & 0 & 0 \\ 0 & 0 & 0 & 0 & 0 \end{bmatrix}$$

Using article's Eq 22, one can specify the likelihood function $L(r_1 \mid w_P^{1,2}, w_Q^{1,2})$ which will be maximized in order obtain $\hat{r}$. Since the simulated data set contains no error, it is possible to simply count the number of individuals in each one of the dosage category and represent them in a double entry table

Table S3.4 : Number of individuals in each one of the dosage-based genotypic classes for markers $M_1$ and $M_2$

| | $d^{M_2}$ | | |
|---|---|---|---|
| $d^{M_1}$ | 0 | 1 | 2 |
| 0 | 0 | 0 | 0 |
| 1 | 1 | 4 | 5 |
| 2 | 7 | 8 | 0 |
| 3 | 3 | 4 | 0 |

For this data set, the likelihood function is

$$L(r_1 \mid w_P^{1,2} = 0, w_Q^{1,2} = 0) \propto \left[\frac{2 + 4r_1 - r_1{}^2}{36}\right] \left[\frac{4 - r_1 + r_1{}^2}{18}\right]^{12} \left[\frac{5 - 2r_1 - r_1{}^2}{36}\right]^{12} \left[\frac{(r_1 - 2)(r_1 - 1)}{36}\right]^{3} \left[\frac{1 + 2r_1 - 2r_1{}^2}{36}\right]^{2} \quad (1)$$

The maximum likelihood estimator of $r_1$ can be obtained using any iterative procedure. In this example, we used the golden section search combined with parabolic interpolation described in [1] and implemented in the function `optim` in R software. Given $w_P^{1,2} = 0$ and $w_Q^{1,2} = 0$, $\hat{r} = 4.58 \times 10^{-5}$ and the associated natural logarithm of the likelihood is $-60.47$. Traditionally, genetic mapping procedures use LOD Scores (base-10 log likelihood ratio) as a means for decision making. In our method we use two LOD Scores: $(i)$ first takes into account the logarithm of the ratio between the highest likelihood among all linkage phase configurations and the likelihood of all possible linkage phase configurations; (*ii*) the second uses the ratio between the model under $H_a : r = \hat{r}$ and under the null hypothesis of no linkage $H_o : r = 0.5$, given a linkage phase configuration. The results for the pair of markers $(M_1, M_2)$ are presented in Table S3.5

| $w_P^{1,2}$ | $w_Q^{1,2}$ | $\hat{r}$ | log-likelihood | $LOD_{ph}$ | $LOD_{\hat{r}}$ |
|---|---|---|---|---|---|
| 0 | 0 | 0.00 | −60.47 | 0.00 | 2.49 |
| 0 | 1 | 0.50 | −66.20 | 2.49 | 0.00 |
| 1 | 0 | 0.50 | −66.20 | 2.49 | 0.00 |
| 1 | 1 | 0.50 | −66.20 | 2.49 | 0.00 |

Table S3.5 : Recombination fraction estimates and associated LOD Scores between markers $(M_1, M_2)$. $LOD_{ph}$ indicates the logarithm to base 10 of the ratio between the highest likelihood among all linkage phase configurations and the likelihood of all possible linkage phase configurations. $LOD_{\hat{r}}$ uses the ratio between the model under $H_a : r = \hat{r}$ and under the null hypothesis of no linkage $H_o : r = 0.5$.

The same reasoning can be applied for pairs $(M_1, M_3)$ and $(M_2, M_3)$:

| $w_P^{1,3}$ | $w_Q^{1,3}$ | $\hat{r}$ | log-likelihood | $LOD_{ph}$ | $LOD_{\hat{r}}$ |
|---|---|---|---|---|---|
| 1 | 0 | 0.00 | −58.61 | 0.00 | 2.21 |
| 0 | 1 | 0.00 | −59.06 | 0.20 | 2.18 |
| 1 | 1 | 0.50 | −63.68 | 2.21 | 0.00 |
| 0 | 0 | 0.42 | −63.77 | 2.24 | 0.13 |
| 2 | 0 | 0.50 | −64.08 | 2.38 | 0.00 |
| 2 | 1 | 0.50 | −64.08 | 2.38 | 0.00 |

| $w_P^{2,3}$ | $w_Q^{2,3}$ | $\hat{r}$ | log-likelihood | $LOD_{ph}$ | $LOD_{\hat{r}}$ |
|---|---|---|---|---|---|
| 0 | 1 | 0.00 | −60.59 | 0.00 | 0.34 |
| 1 | 0 | 0.00 | −60.68 | 0.04 | 0.30 |
| 1 | 1 | 0.50 | −61.37 | 0.34 | 0.00 |
| 0 | 0 | 0.50 | −61.37 | 0.34 | 0.00 |

Table S3.6 : Recombination fraction estimates and associated LOD Scores between markers $(M_1, M_3)$ and $(M_2, M_3)$. $LOD_{ph}$ indicates the logarithm to base 10 of the ratio between the highest likelihood among all linkage phase configurations and the likelihood of all possible linkage phase configurations. $LOD_{\hat{r}}$ uses the ratio between the model under $H_a : r = \hat{r}$ and under the null hypothesis of no linkage $H_o : r = 0.5$.

Analyzing Tables S3.5 and S3.6, it is possible to narrow down the number of linkage phase configurations that need to be evaluated using the multipoint procedure. Let us assume $LOD_{ph} > 2.00$ the threshold for the linkage phase elimination criteria. For the first two markers $M_1$ and $M_2$, $\Phi_P^{1,2}(\eta = 2)$ contains only situations where $w_P^{1,2} = 0$ and $w_Q^{1,2} = 0$ (white cells in Table S3.2). In this case, the size of $\Phi_P^{1,2}(\eta = 2)$ is $U = 12$ and the $\mathbf{H}_2^u$ matrices are

$$\mathbf{H}_2^1 = \begin{pmatrix}1 & 0\\1 & 0\\0 & 1\\0 & 0\end{pmatrix}, \quad \mathbf{H}_2^2 = \begin{pmatrix}1 & 0\\1 & 0\\0 & 0\\0 & 1\end{pmatrix}, \quad \mathbf{H}_2^3 = \begin{pmatrix}1 & 0\\0 & 1\\1 & 0\\0 & 0\end{pmatrix}, \quad \mathbf{H}_2^4 = \begin{pmatrix}1 & 0\\0 & 0\\1 & 0\\0 & 1\end{pmatrix}, \quad \mathbf{H}_2^5 = \begin{pmatrix}1 & 0\\0 & 1\\0 & 0\\1 & 0\end{pmatrix}, \quad \mathbf{H}_2^6 = \begin{pmatrix}1 & 0\\0 & 0\\0 & 1\\1 & 0\end{pmatrix}$$

$$\mathbf{H}_2^7 = \begin{pmatrix}0 & 1\\1 & 0\\1 & 0\\0 & 0\end{pmatrix}, \quad \mathbf{H}_2^8 = \begin{pmatrix}0 & 0\\1 & 0\\1 & 0\\0 & 1\end{pmatrix}, \quad \mathbf{H}_2^9 = \begin{pmatrix}0 & 1\\1 & 0\\0 & 0\\1 & 0\end{pmatrix}, \quad \mathbf{H}_2^{10} = \begin{pmatrix}0 & 0\\1 & 0\\0 & 1\\1 & 0\end{pmatrix}, \quad \mathbf{H}_2^{11} = \begin{pmatrix}0 & 0\\0 & 1\\1 & 0\\1 & 0\end{pmatrix}, \quad \mathbf{H}_2^{12} = \begin{pmatrix}0 & 0\\0 & 1\\1 & 0\\1 & 0\end{pmatrix}$$

Notice that any of the matrices can be obtained from any other just by permuting the rows. Thus, we chose the configuration related to the first matrix ($\mathbf{H}_2^1 \rightarrow \{\{P_1^1, P_1^2\}, \{P_2^3\}\}$) to proceed with the analysis. The same reasoning applies to parent $Q$. In that case, we choose the configuration $\{\{Q_1^1\}, \{Q_2^2\}\}$. Notice that, at threshold level $\eta = 2$, the two-point analysis did not provide any information between markers $M_2$ and $M_3$. In any case, it is possible to remove redundant configurations when evaluating $\mathbf{H}_3$. For markers $M_1$ and $M_3$ the procedure is similar to markers $M_1$ and $M_2$. Figure  shows possible configurations for parents $P$ and $Q$ using two-point information for pair of markers $M_1$-$M_2$ and $M_2$-$M_3$. Using informations of $M_1$ and $M_3$, it is possible to narrow down the number of configurations to be analyzed by multipoint procedures.

In linkage groups with more markers, this procedure can be applied until there is no more information to be extracted from the two-point analysis at the assumed threshold level. In this simulation, only four linkage phase configurations needed to be evaluated using multipoint procedure.

The remaining linkage phase configurations were analyzed using the full HMM procedure. The log-likelihood and LOD Scores of the models are presented in Table S3.7

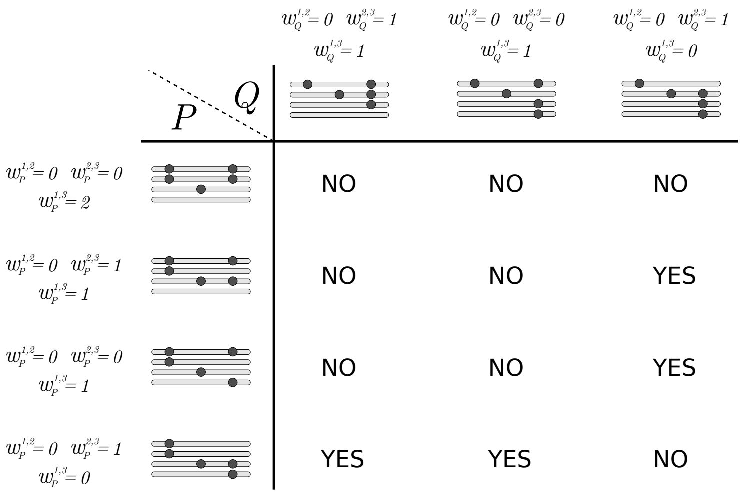


Figure S3.2 : Possible linkage phase configurations in parents $P$ and $Q$ eliminating the ones with $LOD_{ph} > 2.00$ for pairs of markers $(M_1, M_2)$ and $(M_2, M_3)$. $w_P^{k,k'}$ and $w_Q^{k,k'}$ indicate the number of homologous chromosomes that share allelic variants for loci $k$ and $k'$ in parents $P$ and $Q$, respectively. Some of the combinations can be eliminated using information from the pair $(M_1, M_3)$. For instance, with a threshold of $LOD_{ph} > 2.00$, any of the combination in the first line of the table is not possible since $w_P^{1,3} = 2$ and the associated $LOD_{ph}$ regardless the configuration in $Q$ is 2.38. Using this reasoning it is possible to eliminate several configurations (marked with a NO) based on the information from the pair $(M_1, M_3)$. Thus, there are only four configurations left to be tested using the multipoint procedure.

Table S3.7

| Linkage phase | | | $LOD$ | log-likelihood |
|---|---|---|---|---|
| $M_1$ | $M_2$ | $M_3$ | | |
| $\{P_1^1 P_1^2\}$ | $\{P_2^3\}$ | $\{P_3^1 P_3^4\}$ | 0.00 | −81.90 |
| $\{Q_1^1\}$ | $\{Q_2^2\}$ | $\{Q_3^2 Q_3^3 Q_3^4\}$ | | |
| $\{P_1^1 P_1^2\}$ | $\{P_2^3\}$ | $\{P_3^3 P_3^4\}$ | 1.75 | −85.93 |
| $\{Q_1^1\}$ | $\{Q_2^2\}$ | $\{Q_3^1 Q_3^3 Q_3^4\}$ | | |
| $\{P_1^1 P_1^2\}$ | $\{P_2^3\}$ | $\{P_3^1 P_3^3\}$ | 3.07 | −88.96 |
| $\{Q_1^1\}$ | $\{Q_2^2\}$ | $\{Q_3^2 Q_3^3 Q_3^4\}$ | | |
| $\{P_1^1 P_1^2\}$ | $\{P_2^3\}$ | $\{P_3^3 P_3^4\}$ | 3.45 | −89.86 |
| $\{Q_1^1\}$ | $\{Q_2^2\}$ | $\{Q_3^2 Q_3^3 Q_3^4\}$ | | |

Finally, the best configuration obtained using both two-point and multipoint procedure is

Figure S3.3

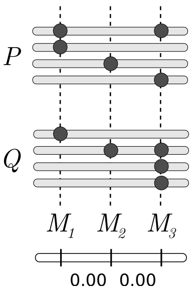


Figure S3.3

This configuration is a permutation of the homologous chromosomes of the simulation. In practical terms it consists in the same linkage phase configuration. The codes to perform the analysis step by step, as presented in this supplement, are available at `https://github.com/mmollina/Autopolyploid_Linkage/blob/master/src/SI3_example.R`

| | | | | $j'$ | | | | | | | | | | | | | | | | | | | | | | | | | | | | | | | | | | | |
|---|---|---|---|---|---|---|---|---|---|---|---|---|---|---|---|---|---|---|---|---|---|---|---|---|---|---|---|---|---|---|---|---|---|---|---|---|---|---|---|
| $j$ | $i$ | $h$ | $\mathcal{G}^4_{k,j} = \{\theta^4_{P,i}, \theta^4_{Q,h}\}$ | 1 | 2 | 3 | 4 | 5 | 6 | 7 | 8 | 9 | 10 | 11 | 12 | 13 | 14 | 15 | 16 | 17 | 18 | 19 | 20 | 21 | 22 | 23 | 24 | 25 | 26 | 27 | 28 | 29 | 30 | 31 | 32 | 33 | 34 | 35 | 36 |
| 1 | 1 | 1 | $\{(P_k^1,P_k^2),(Q_k^1,Q_k^2)\}$ | $a^2$ | $ab$ | $ab$ | $ab$ | $ab$ | $ac$ | $ab$ | $b^2$ | $b^2$ | $b^2$ | $b^2$ | $bc$ | $ab$ | $b^2$ | $b^2$ | $b^2$ | $b^2$ | $bc$ | $ab$ | $b^2$ | $b^2$ | $b^2$ | $b^2$ | $bc$ | $ab$ | $b^2$ | $b^2$ | $b^2$ | $b^2$ | $bc$ | $ac$ | $bc$ | $bc$ | $bc$ | $bc$ | $c^2$ |
| 2 | 1 | 2 | $\{(P_k^1,P_k^2),(Q_k^1,Q_k^3)\}$ | $ab$ | $a^2$ | $ab$ | $ab$ | $ac$ | $ab$ | $b^2$ | $ab$ | $b^2$ | $b^2$ | $bc$ | $b^2$ | $b^2$ | $ab$ | $b^2$ | $b^2$ | $bc$ | $b^2$ | $b^2$ | $ab$ | $b^2$ | $b^2$ | $bc$ | $b^2$ | $b^2$ | $ab$ | $b^2$ | $b^2$ | $bc$ | $b^2$ | $bc$ | $ac$ | $bc$ | $bc$ | $c^2$ | $bc$ |
| 3 | 1 | 3 | $\{(P_k^1,P_k^2),(Q_k^1,Q_k^4)\}$ | $ab$ | $ab$ | $a^2$ | $ac$ | $ab$ | $ab$ | $b^2$ | $b^2$ | $ab$ | $bc$ | $b^2$ | $b^2$ | $b^2$ | $b^2$ | $ab$ | $bc$ | $b^2$ | $b^2$ | $b^2$ | $b^2$ | $ab$ | $bc$ | $b^2$ | $b^2$ | $b^2$ | $b^2$ | $ab$ | $bc$ | $b^2$ | $b^2$ | $bc$ | $bc$ | $ac$ | $c^2$ | $bc$ | $bc$ |
| 4 | 1 | 4 | $\{(P_k^1,P_k^2),(Q_k^2,Q_k^3)\}$ | $ab$ | $ab$ | $ac$ | $a^2$ | $ab$ | $ab$ | $b^2$ | $b^2$ | $bc$ | $ab$ | $b^2$ | $b^2$ | $b^2$ | $b^2$ | $bc$ | $ab$ | $b^2$ | $b^2$ | $b^2$ | $b^2$ | $bc$ | $ab$ | $b^2$ | $b^2$ | $b^2$ | $b^2$ | $bc$ | $ab$ | $b^2$ | $b^2$ | $bc$ | $bc$ | $c^2$ | $ac$ | $bc$ | $bc$ |
| 5 | 1 | 5 | $\{(P_k^1,P_k^2),(Q_k^2,Q_k^4)\}$ | $ab$ | $ac$ | $ab$ | $ab$ | $a^2$ | $ab$ | $b^2$ | $bc$ | $b^2$ | $b^2$ | $ab$ | $b^2$ | $b^2$ | $bc$ | $b^2$ | $b^2$ | $ab$ | $b^2$ | $b^2$ | $bc$ | $b^2$ | $b^2$ | $ab$ | $b^2$ | $b^2$ | $bc$ | $b^2$ | $b^2$ | $ab$ | $b^2$ | $bc$ | $c^2$ | $bc$ | $bc$ | $ac$ | $bc$ |
| 6 | 1 | 6 | $\{(P_k^1,P_k^2),(Q_k^3,Q_k^4)\}$ | $ac$ | $ab$ | $ab$ | $ab$ | $ab$ | $a^2$ | $bc$ | $b^2$ | $b^2$ | $b^2$ | $b^2$ | $ab$ | $bc$ | $b^2$ | $b^2$ | $b^2$ | $b^2$ | $ab$ | $bc$ | $b^2$ | $b^2$ | $b^2$ | $b^2$ | $ab$ | $bc$ | $b^2$ | $b^2$ | $b^2$ | $b^2$ | $ab$ | $c^2$ | $bc$ | $bc$ | $bc$ | $bc$ | $ac$ |
| 7 | 2 | 1 | $\{(P_k^1,P_k^3),(Q_k^1,Q_k^2)\}$ | $ab$ | $b^2$ | $b^2$ | $b^2$ | $b^2$ | $bc$ | $a^2$ | $ab$ | $ab$ | $ab$ | $ab$ | $ac$ | $ab$ | $b^2$ | $b^2$ | $b^2$ | $b^2$ | $bc$ | $ab$ | $b^2$ | $b^2$ | $b^2$ | $b^2$ | $bc$ | $ac$ | $bc$ | $bc$ | $bc$ | $bc$ | $c^2$ | $ab$ | $b^2$ | $b^2$ | $b^2$ | $b^2$ | $bc$ |
| 8 | 2 | 2 | $\{(P_k^1,P_k^3),(Q_k^1,Q_k^3)\}$ | $b^2$ | $ab$ | $b^2$ | $b^2$ | $bc$ | $b^2$ | $ab$ | $a^2$ | $ab$ | $ab$ | $ac$ | $ab$ | $b^2$ | $ab$ | $b^2$ | $b^2$ | $bc$ | $b^2$ | $b^2$ | $ab$ | $b^2$ | $b^2$ | $bc$ | $b^2$ | $bc$ | $ac$ | $bc$ | $bc$ | $c^2$ | $bc$ | $b^2$ | $ab$ | $b^2$ | $b^2$ | $bc$ | $b^2$ |
| 9 | 2 | 3 | $\{(P_k^1,P_k^3),(Q_k^1,Q_k^4)\}$ | $b^2$ | $b^2$ | $ab$ | $bc$ | $b^2$ | $b^2$ | $ab$ | $ab$ | $a^2$ | $ac$ | $ab$ | $ab$ | $b^2$ | $b^2$ | $ab$ | $bc$ | $b^2$ | $b^2$ | $b^2$ | $b^2$ | $ab$ | $bc$ | $b^2$ | $b^2$ | $bc$ | $bc$ | $ac$ | $c^2$ | $bc$ | $bc$ | $b^2$ | $b^2$ | $ab$ | $bc$ | $b^2$ | $b^2$ |
| 10 | 2 | 4 | $\{(P_k^1,P_k^3),(Q_k^2,Q_k^3)\}$ | $b^2$ | $b^2$ | $bc$ | $ab$ | $b^2$ | $b^2$ | $ab$ | $ab$ | $ac$ | $a^2$ | $ab$ | $ab$ | $b^2$ | $b^2$ | $bc$ | $ab$ | $b^2$ | $b^2$ | $b^2$ | $b^2$ | $bc$ | $ab$ | $b^2$ | $b^2$ | $bc$ | $bc$ | $c^2$ | $ac$ | $bc$ | $bc$ | $b^2$ | $b^2$ | $bc$ | $ab$ | $b^2$ | $b^2$ |
| 11 | 2 | 5 | $\{(P_k^1,P_k^3),(Q_k^2,Q_k^4)\}$ | $b^2$ | $bc$ | $b^2$ | $b^2$ | $ab$ | $b^2$ | $ab$ | $ac$ | $ab$ | $ab$ | $a^2$ | $ab$ | $b^2$ | $bc$ | $b^2$ | $b^2$ | $ab$ | $b^2$ | $b^2$ | $bc$ | $b^2$ | $b^2$ | $ab$ | $b^2$ | $bc$ | $c^2$ | $bc$ | $bc$ | $ac$ | $bc$ | $b^2$ | $bc$ | $b^2$ | $b^2$ | $ab$ | $b^2$ |
| 12 | 2 | 6 | $\{(P_k^1,P_k^3),(Q_k^3,Q_k^4)\}$ | $bc$ | $b^2$ | $b^2$ | $b^2$ | $b^2$ | $ab$ | $ac$ | $ab$ | $ab$ | $ab$ | $ab$ | $a^2$ | $bc$ | $b^2$ | $b^2$ | $b^2$ | $b^2$ | $ab$ | $bc$ | $b^2$ | $b^2$ | $b^2$ | $b^2$ | $ab$ | $c^2$ | $bc$ | $bc$ | $bc$ | $bc$ | $ac$ | $bc$ | $b^2$ | $b^2$ | $b^2$ | $b^2$ | $ab$ |
| 13 | 3 | 1 | $\{(P_k^1,P_k^4),(Q_k^1,Q_k^2)\}$ | $ab$ | $b^2$ | $b^2$ | $b^2$ | $b^2$ | $bc$ | $ab$ | $b^2$ | $b^2$ | $b^2$ | $b^2$ | $bc$ | $a^2$ | $ab$ | $ab$ | $ab$ | $ab$ | $ac$ | $ac$ | $bc$ | $bc$ | $bc$ | $bc$ | $c^2$ | $ab$ | $b^2$ | $b^2$ | $b^2$ | $b^2$ | $bc$ | $ab$ | $b^2$ | $b^2$ | $b^2$ | $b^2$ | $bc$ |
| 14 | 3 | 2 | $\{(P_k^1,P_k^4),(Q_k^1,Q_k^3)\}$ | $b^2$ | $ab$ | $b^2$ | $b^2$ | $bc$ | $b^2$ | $b^2$ | $ab$ | $b^2$ | $b^2$ | $bc$ | $b^2$ | $ab$ | $a^2$ | $ab$ | $ab$ | $ac$ | $ab$ | $bc$ | $ac$ | $bc$ | $bc$ | $c^2$ | $bc$ | $b^2$ | $ab$ | $b^2$ | $b^2$ | $bc$ | $b^2$ | $b^2$ | $ab$ | $b^2$ | $b^2$ | $bc$ | $b^2$ |
| 15 | 3 | 3 | $\{(P_k^1,P_k^4),(Q_k^1,Q_k^4)\}$ | $b^2$ | $b^2$ | $ab$ | $bc$ | $b^2$ | $b^2$ | $b^2$ | $b^2$ | $ab$ | $bc$ | $b^2$ | $b^2$ | $ab$ | $ab$ | $a^2$ | $ac$ | $ab$ | $ab$ | $bc$ | $bc$ | $ac$ | $c^2$ | $bc$ | $bc$ | $b^2$ | $b^2$ | $ab$ | $bc$ | $b^2$ | $b^2$ | $b^2$ | $b^2$ | $ab$ | $bc$ | $b^2$ | $b^2$ |
| 16 | 3 | 4 | $\{(P_k^1,P_k^4),(Q_k^2,Q_k^3)\}$ | $b^2$ | $b^2$ | $bc$ | $ab$ | $b^2$ | $b^2$ | $b^2$ | $b^2$ | $bc$ | $ab$ | $b^2$ | $b^2$ | $ab$ | $ab$ | $ac$ | $a^2$ | $ab$ | $ab$ | $bc$ | $bc$ | $c^2$ | $ac$ | $bc$ | $bc$ | $b^2$ | $b^2$ | $bc$ | $ab$ | $b^2$ | $b^2$ | $b^2$ | $b^2$ | $bc$ | $ab$ | $b^2$ | $b^2$ |
| 17 | 3 | 5 | $\{(P_k^1,P_k^4),(Q_k^2,Q_k^4)\}$ | $b^2$ | $bc$ | $b^2$ | $b^2$ | $ab$ | $b^2$ | $b^2$ | $bc$ | $b^2$ | $b^2$ | $ab$ | $b^2$ | $ab$ | $ac$ | $ab$ | $ab$ | $a^2$ | $ab$ | $bc$ | $c^2$ | $bc$ | $bc$ | $ac$ | $bc$ | $b^2$ | $bc$ | $b^2$ | $b^2$ | $ab$ | $b^2$ | $b^2$ | $bc$ | $b^2$ | $b^2$ | $ab$ | $b^2$ |
| 18 | 3 | 6 | $\{(P_k^1,P_k^4),(Q_k^3,Q_k^4)\}$ | $bc$ | $b^2$ | $b^2$ | $b^2$ | $b^2$ | $ab$ | $bc$ | $b^2$ | $b^2$ | $b^2$ | $b^2$ | $ab$ | $ac$ | $ab$ | $ab$ | $ab$ | $ab$ | $a^2$ | $c^2$ | $bc$ | $bc$ | $bc$ | $bc$ | $ac$ | $bc$ | $b^2$ | $b^2$ | $b^2$ | $b^2$ | $ab$ | $bc$ | $b^2$ | $b^2$ | $b^2$ | $b^2$ | $ab$ |
| 19 | 4 | 1 | $\{(P_k^2,P_k^3),(Q_k^1,Q_k^2)\}$ | $ab$ | $b^2$ | $b^2$ | $b^2$ | $b^2$ | $bc$ | $ab$ | $b^2$ | $b^2$ | $b^2$ | $b^2$ | $bc$ | $ac$ | $bc$ | $bc$ | $bc$ | $bc$ | $c^2$ | $a^2$ | $ab$ | $ab$ | $ab$ | $ab$ | $ac$ | $ab$ | $b^2$ | $b^2$ | $b^2$ | $b^2$ | $bc$ | $ab$ | $b^2$ | $b^2$ | $b^2$ | $b^2$ | $bc$ |
| 20 | 4 | 2 | $\{(P_k^2,P_k^3),(Q_k^1,Q_k^3)\}$ | $b^2$ | $ab$ | $b^2$ | $b^2$ | $bc$ | $b^2$ | $b^2$ | $ab$ | $b^2$ | $b^2$ | $bc$ | $b^2$ | $bc$ | $ac$ | $bc$ | $bc$ | $c^2$ | $bc$ | $ab$ | $a^2$ | $ab$ | $ab$ | $ac$ | $ab$ | $b^2$ | $ab$ | $b^2$ | $b^2$ | $bc$ | $b^2$ | $b^2$ | $ab$ | $b^2$ | $b^2$ | $bc$ | $b^2$ |
| 21 | 4 | 3 | $\{(P_k^2,P_k^3),(Q_k^1,Q_k^4)\}$ | $b^2$ | $b^2$ | $ab$ | $bc$ | $b^2$ | $b^2$ | $b^2$ | $b^2$ | $ab$ | $bc$ | $b^2$ | $b^2$ | $bc$ | $bc$ | $ac$ | $c^2$ | $bc$ | $bc$ | $ab$ | $ab$ | $a^2$ | $ac$ | $ab$ | $ab$ | $b^2$ | $b^2$ | $ab$ | $bc$ | $b^2$ | $b^2$ | $b^2$ | $b^2$ | $ab$ | $bc$ | $b^2$ | $b^2$ |
| 22 | 4 | 4 | $\{(P_k^2,P_k^3),(Q_k^2,Q_k^3)\}$ | $b^2$ | $b^2$ | $bc$ | $ab$ | $b^2$ | $b^2$ | $b^2$ | $b^2$ | $bc$ | $ab$ | $b^2$ | $b^2$ | $bc$ | $bc$ | $c^2$ | $ac$ | $bc$ | $bc$ | $ab$ | $ab$ | $ac$ | $a^2$ | $ab$ | $ab$ | $b^2$ | $b^2$ | $bc$ | $ab$ | $b^2$ | $b^2$ | $b^2$ | $b^2$ | $bc$ | $ab$ | $b^2$ | $b^2$ |
| 23 | 4 | 5 | $\{(P_k^2,P_k^3),(Q_k^2,Q_k^4)\}$ | $b^2$ | $bc$ | $b^2$ | $b^2$ | $ab$ | $b^2$ | $b^2$ | $bc$ | $b^2$ | $b^2$ | $ab$ | $b^2$ | $bc$ | $c^2$ | $bc$ | $bc$ | $ac$ | $bc$ | $ab$ | $ac$ | $ab$ | $ab$ | $a^2$ | $ab$ | $b^2$ | $bc$ | $b^2$ | $b^2$ | $ab$ | $b^2$ | $b^2$ | $bc$ | $b^2$ | $b^2$ | $ab$ | $b^2$ |
| 24 | 4 | 6 | $\{(P_k^2,P_k^3),(Q_k^3,Q_k^4)\}$ | $bc$ | $b^2$ | $b^2$ | $b^2$ | $b^2$ | $ab$ | $bc$ | $b^2$ | $b^2$ | $b^2$ | $b^2$ | $ab$ | $c^2$ | $bc$ | $bc$ | $bc$ | $bc$ | $ac$ | $ac$ | $ab$ | $ab$ | $ab$ | $ab$ | $a^2$ | $bc$ | $b^2$ | $b^2$ | $b^2$ | $b^2$ | $ab$ | $bc$ | $b^2$ | $b^2$ | $b^2$ | $b^2$ | $ab$ |
| 25 | 5 | 1 | $\{(P_k^2,P_k^4),(Q_k^1,Q_k^2)\}$ | $ab$ | $b^2$ | $b^2$ | $b^2$ | $b^2$ | $bc$ | $ac$ | $bc$ | $bc$ | $bc$ | $bc$ | $c^2$ | $ab$ | $b^2$ | $b^2$ | $b^2$ | $b^2$ | $bc$ | $ab$ | $b^2$ | $b^2$ | $b^2$ | $b^2$ | $bc$ | $a^2$ | $ab$ | $ab$ | $ab$ | $ab$ | $ac$ | $ab$ | $b^2$ | $b^2$ | $b^2$ | $b^2$ | $bc$ |
| 26 | 5 | 2 | $\{(P_k^2,P_k^4),(Q_k^1,Q_k^3)\}$ | $b^2$ | $ab$ | $b^2$ | $b^2$ | $bc$ | $b^2$ | $bc$ | $ac$ | $bc$ | $bc$ | $c^2$ | $bc$ | $b^2$ | $ab$ | $b^2$ | $b^2$ | $bc$ | $b^2$ | $b^2$ | $ab$ | $b^2$ | $b^2$ | $bc$ | $b^2$ | $ab$ | $a^2$ | $ab$ | $ab$ | $ac$ | $ab$ | $b^2$ | $ab$ | $b^2$ | $b^2$ | $bc$ | $b^2$ |
| 27 | 5 | 3 | $\{(P_k^2,P_k^4),(Q_k^1,Q_k^4)\}$ | $b^2$ | $b^2$ | $ab$ | $bc$ | $b^2$ | $b^2$ | $bc$ | $bc$ | $ac$ | $c^2$ | $bc$ | $bc$ | $b^2$ | $b^2$ | $ab$ | $bc$ | $b^2$ | $b^2$ | $b^2$ | $b^2$ | $ab$ | $bc$ | $b^2$ | $b^2$ | $ab$ | $ab$ | $a^2$ | $ac$ | $ab$ | $ab$ | $b^2$ | $b^2$ | $ab$ | $bc$ | $b^2$ | $b^2$ |
| 28 | 5 | 4 | $\{(P_k^2,P_k^4),(Q_k^2,Q_k^3)\}$ | $b^2$ | $b^2$ | $bc$ | $ab$ | $b^2$ | $b^2$ | $bc$ | $bc$ | $c^2$ | $ac$ | $bc$ | $bc$ | $b^2$ | $b^2$ | $bc$ | $ab$ | $b^2$ | $b^2$ | $b^2$ | $b^2$ | $bc$ | $ab$ | $b^2$ | $b^2$ | $ab$ | $ab$ | $ac$ | $a^2$ | $ab$ | $ab$ | $b^2$ | $b^2$ | $bc$ | $ab$ | $b^2$ | $b^2$ |
| 29 | 5 | 5 | $\{(P_k^2,P_k^4),(Q_k^2,Q_k^4)\}$ | $b^2$ | $bc$ | $b^2$ | $b^2$ | $ab$ | $b^2$ | $bc$ | $c^2$ | $bc$ | $bc$ | $ac$ | $bc$ | $b^2$ | $bc$ | $b^2$ | $b^2$ | $ab$ | $b^2$ | $b^2$ | $bc$ | $b^2$ | $b^2$ | $ab$ | $b^2$ | $ab$ | $ac$ | $ab$ | $ab$ | $a^2$ | $ab$ | $b^2$ | $bc$ | $b^2$ | $b^2$ | $ab$ | $b^2$ |
| 30 | 5 | 6 | $\{(P_k^2,P_k^4),(Q_k^3,Q_k^4)\}$ | $bc$ | $b^2$ | $b^2$ | $b^2$ | $b^2$ | $ab$ | $c^2$ | $bc$ | $bc$ | $bc$ | $bc$ | $ac$ | $bc$ | $b^2$ | $b^2$ | $b^2$ | $b^2$ | $ab$ | $bc$ | $b^2$ | $b^2$ | $b^2$ | $b^2$ | $ab$ | $ac$ | $ab$ | $ab$ | $ab$ | $ab$ | $a^2$ | $bc$ | $b^2$ | $b^2$ | $b^2$ | $b^2$ | $ab$ |
| 31 | 6 | 1 | $\{(P_k^3,P_k^4),(Q_k^1,Q_k^2)\}$ | $ac$ | $bc$ | $bc$ | $bc$ | $bc$ | $c^2$ | $ab$ | $b^2$ | $b^2$ | $b^2$ | $b^2$ | $bc$ | $ab$ | $b^2$ | $b^2$ | $b^2$ | $b^2$ | $bc$ | $ab$ | $b^2$ | $b^2$ | $b^2$ | $b^2$ | $bc$ | $ab$ | $b^2$ | $b^2$ | $b^2$ | $b^2$ | $bc$ | $a^2$ | $ab$ | $ab$ | $ab$ | $ab$ | $ac$ |
| 32 | 6 | 2 | $\{(P_k^3,P_k^4),(Q_k^1,Q_k^3)\}$ | $bc$ | $ac$ | $bc$ | $bc$ | $c^2$ | $bc$ | $b^2$ | $ab$ | $b^2$ | $b^2$ | $bc$ | $b^2$ | $b^2$ | $ab$ | $b^2$ | $b^2$ | $bc$ | $b^2$ | $b^2$ | $ab$ | $b^2$ | $b^2$ | $bc$ | $b^2$ | $b^2$ | $ab$ | $b^2$ | $b^2$ | $bc$ | $b^2$ | $ab$ | $a^2$ | $ab$ | $ab$ | $ac$ | $ab$ |
| 33 | 6 | 3 | $\{(P_k^3,P_k^4),(Q_k^1,Q_k^4)\}$ | $bc$ | $bc$ | $ac$ | $c^2$ | $bc$ | $bc$ | $b^2$ | $b^2$ | $ab$ | $bc$ | $b^2$ | $b^2$ | $b^2$ | $b^2$ | $ab$ | $bc$ | $b^2$ | $b^2$ | $b^2$ | $b^2$ | $ab$ | $bc$ | $b^2$ | $b^2$ | $b^2$ | $b^2$ | $ab$ | $bc$ | $b^2$ | $b^2$ | $ab$ | $ab$ | $a^2$ | $ac$ | $ab$ | $ab$ |
| 34 | 6 | 4 | $\{(P_k^3,P_k^4),(Q_k^2,Q_k^3)\}$ | $bc$ | $bc$ | $c^2$ | $ac$ | $bc$ | $bc$ | $b^2$ | $b^2$ | $bc$ | $ab$ | $b^2$ | $b^2$ | $b^2$ | $b^2$ | $bc$ | $ab$ | $b^2$ | $b^2$ | $b^2$ | $b^2$ | $bc$ | $ab$ | $b^2$ | $b^2$ | $b^2$ | $b^2$ | $bc$ | $ab$ | $b^2$ | $b^2$ | $ab$ | $ab$ | $ac$ | $a^2$ | $ab$ | $ab$ |
| 35 | 6 | 5 | $\{(P_k^3,P_k^4),(Q_k^2,Q_k^4)\}$ | $bc$ | $c^2$ | $bc$ | $bc$ | $ac$ | $bc$ | $b^2$ | $bc$ | $b^2$ | $b^2$ | $ab$ | $b^2$ | $b^2$ | $bc$ | $b^2$ | $b^2$ | $ab$ | $b^2$ | $b^2$ | $bc$ | $b^2$ | $b^2$ | $ab$ | $b^2$ | $b^2$ | $bc$ | $b^2$ | $b^2$ | $ab$ | $b^2$ | $ab$ | $ac$ | $ab$ | $ab$ | $a^2$ | $ab$ |
| 36 | 6 | 6 | $\{(P_k^3,P_k^4),(Q_k^3,Q_k^4)\}$ | $c^2$ | $bc$ | $bc$ | $bc$ | $bc$ | $ac$ | $bc$ | $b^2$ | $b^2$ | $b^2$ | $b^2$ | $ab$ | $bc$ | $b^2$ | $b^2$ | $b^2$ | $b^2$ | $ab$ | $bc$ | $b^2$ | $b^2$ | $b^2$ | $b^2$ | $ab$ | $bc$ | $b^2$ | $b^2$ | $b^2$ | $b^2$ | $ab$ | $ac$ | $ab$ | $ab$ | $ab$ | $ab$ | $a^2$ |

where $a = (1-r)^2$, $b = \frac{(1-r)r}{2}$ and $c = r^2$

Table S3.8 : Full transition space in autotetraploids. The table contain all possible transitions $\Pr(\mathcal{G}^4_{k',j'}|\mathcal{G}^4_{k,j})$ between the 36 possible genotypic states for markers $k$ and $k'$. For simplicity, apart from $j'$, the name of the columns $i'$, $k'$ and $\mathcal{G}^4_{k',j'} = \{\theta^4_{P,i'}, \theta^4_{Q,h'}\}$ were omitted but they are analogous and in the same order as the name of the rows.
