## Supplementary material for "Linkage analysis and haplotype phasing in experimental autopolyploid populations with high ploidy level using hidden Markov models": Figure S4

**Figure S4: Haplotypes for simulation study 1 - Simulated haplotypes with 10 markers and three ploidy levels, namely autotetraploid ($m = 4$), autohexaploid ($m = 6$) and autooctaploid ($m = 8$).**

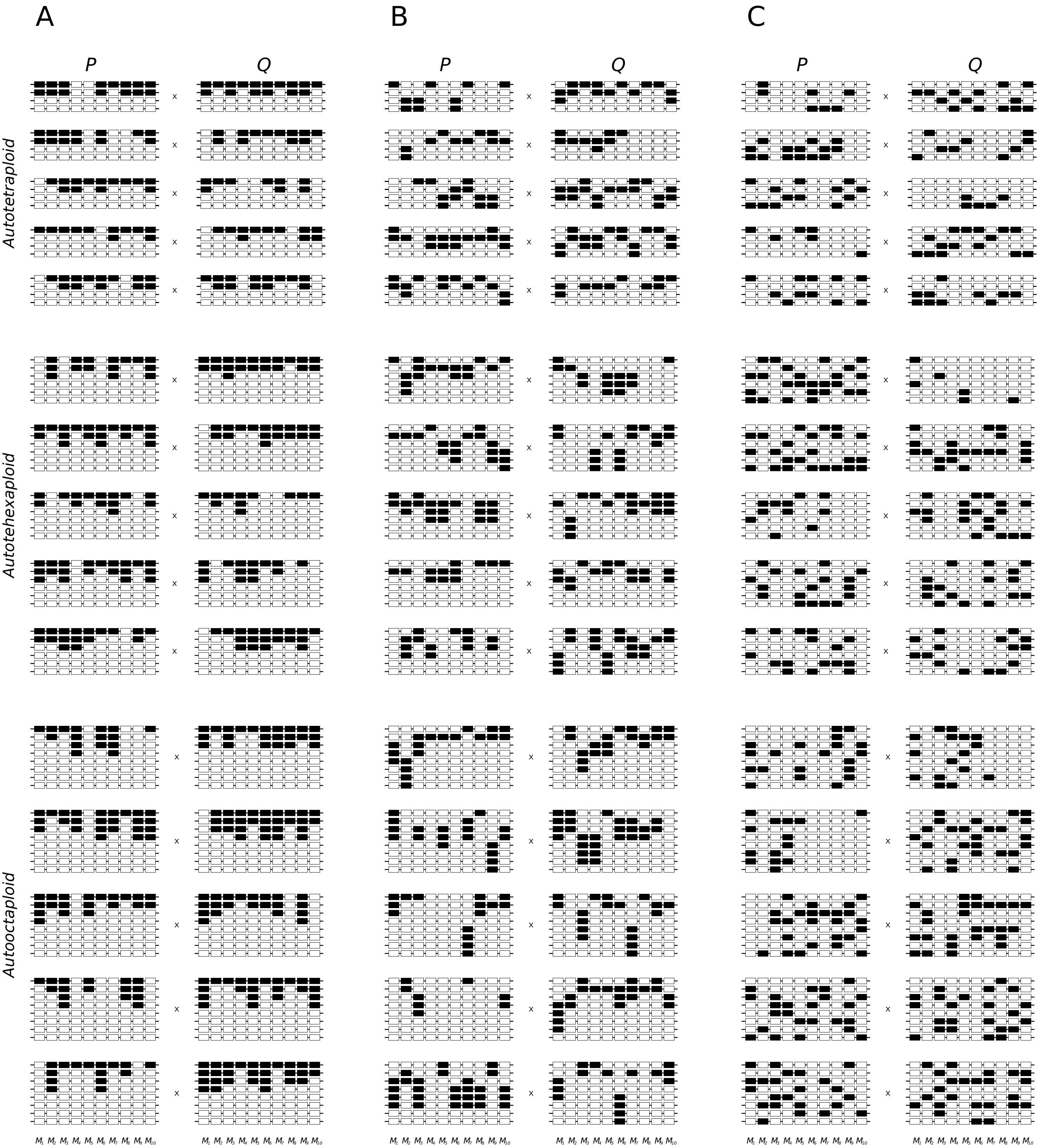


Figure S4 : Simulated haplotypes with 10 markers and three ploidy levels, namely autotetraploid ($m = 4$), autohexaploid ($m = 6$) and autooctaploid ($m = 8$). Black and white rectangles indicate two allelic variants in each marker. In all scenarios, the marker doses varied from zero to $\frac{m}{2}$. Although the method can cope with dosages greater than $\frac{m}{2}$, they are equivalent to lower doses in different linkage phases, therefore they were not considered. Each horizontal line indicates homologous chromosomes which are grouped in homology groups. Three linkage phase scenarios were simulated: In scenario A, allelic variants represented by black rectangle were assigned to the first homologous chromosome in the homology group. The remaining variants of the same type where assigned to the subsequent homologous chromosomes. This yields patterns where allelic variants of the same type were mostly concentrated in the same homologous chromosomes. In B, one type of allelic variant was randomly assigned to one of the first $\frac{m}{2}$ homologous chromosome and the remaining allelic variants of the same type were assigned to the subsequent homologous chromosomes; in C, the allelic variants were randomly assigned to the $m$ homologous chromosomes. Five different parental haplotypes were considered for each linkage phase scenario. In total, 45 parental linkage phase configurations were considered ($3 \times 3 \times 5 = 45$).
