## Supplementary material for "Linkage analysis and haplotype phasing in experimental autopolyploid populations with high ploidy level using hidden Markov models": Figure S5

**Figure S5: Haplotypes for simulation study 2 - Simulated haplotypes with 200 markers and two ploidy levels, namely autotetraploid ($m = 4$) and autohexaploid ($m = 6$).**

### *Autotetraploid*

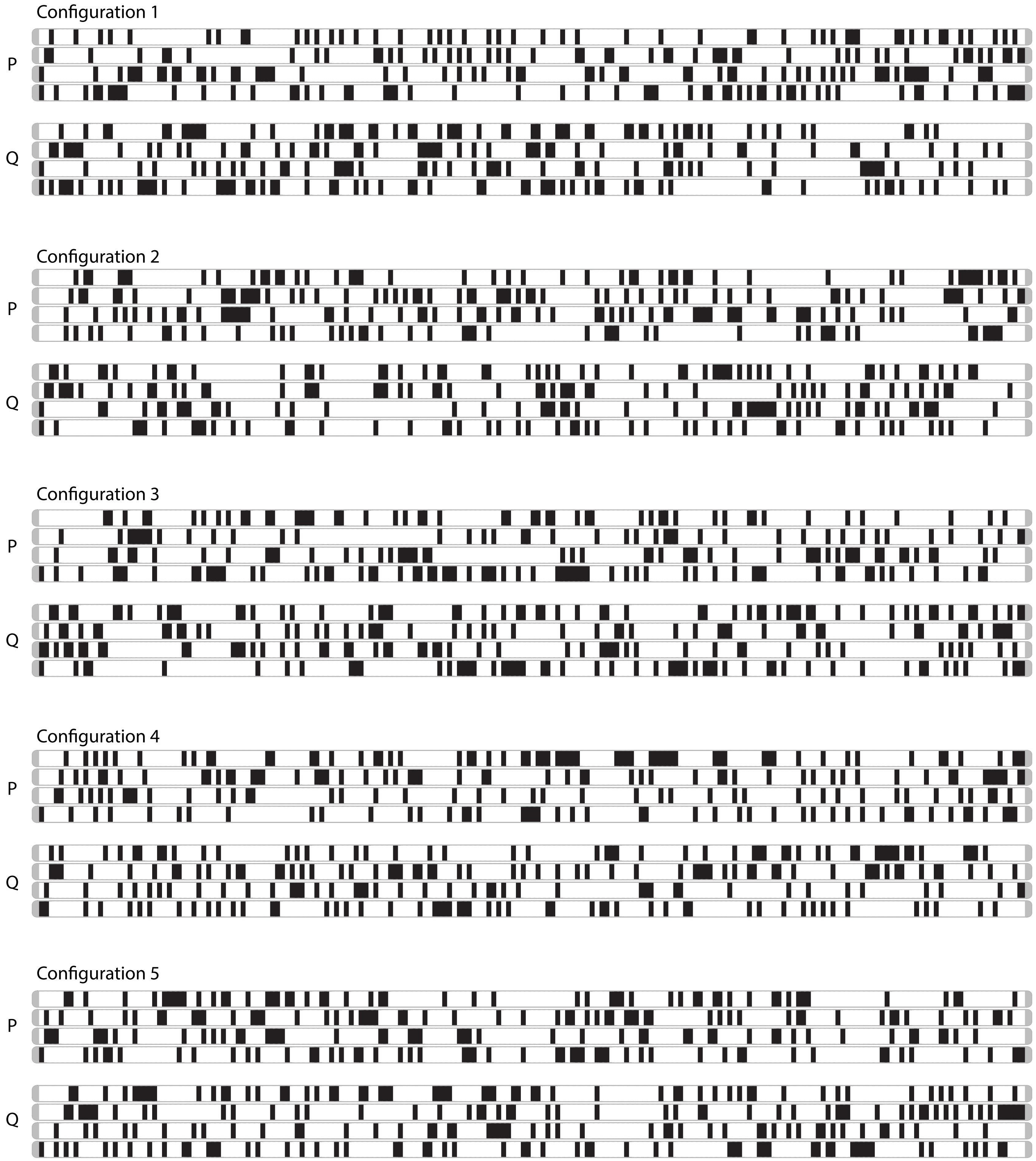


Figure S5 : Simulated haplotypes with 200 markers and two ploidy levels, namely autotetraploid ($m = 4$) and autohexaploid ($m = 6$). Black and white rectangles indicate two allelic variants in each marker. In all configurations, marker doses varied from zero to $\frac{m}{2}$. Each horizontal line indicates homologous chromosomes which are grouped in homology groups. We used scenario C, from simulation study 1, as the template for this second simulation study, i.e., the allelic variants were randomly assigned to the $m$ homologous chromosomes. Five different parental haplotypes were considered for each ploidy level.

### *Autohexaploid*

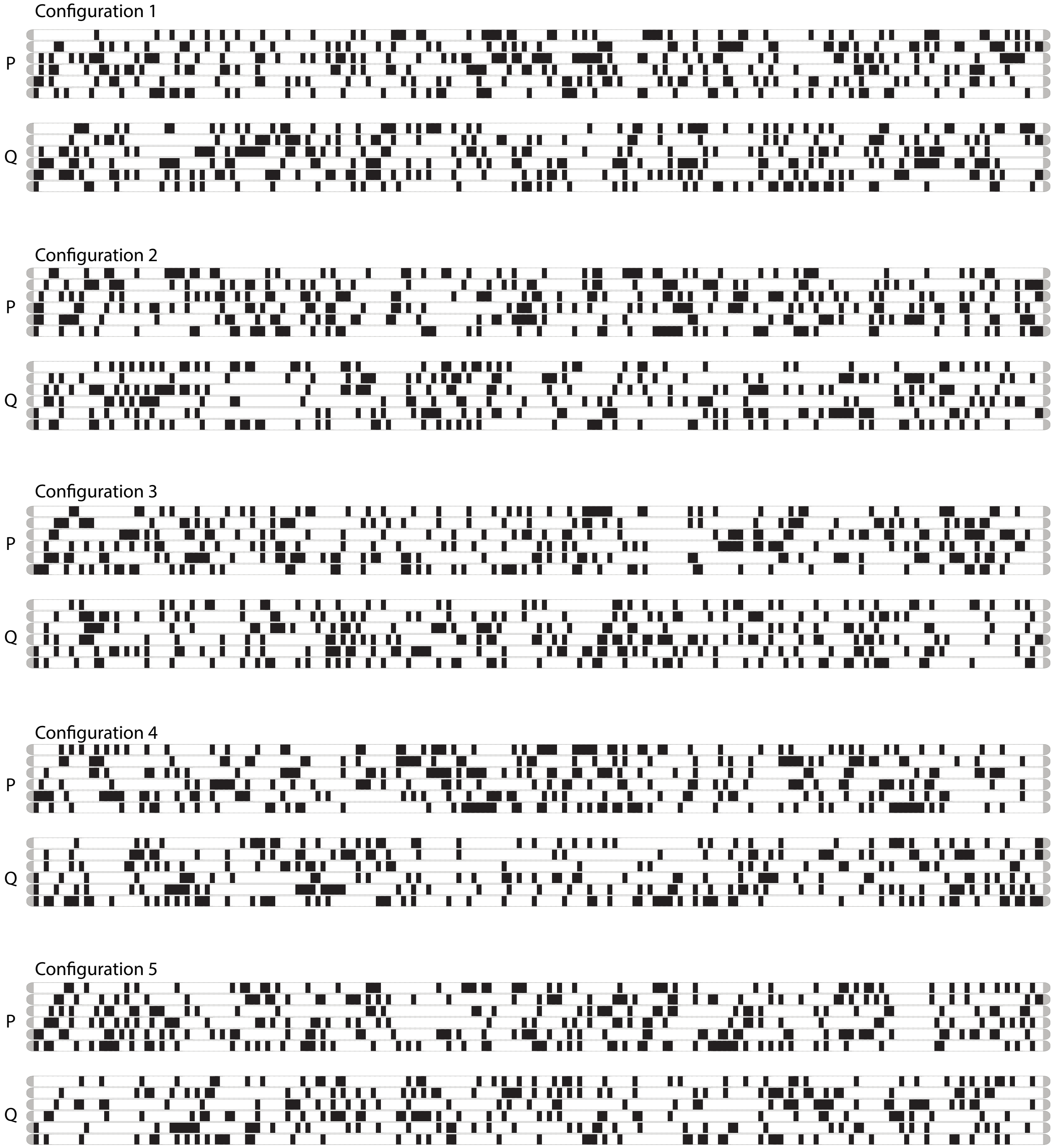


S5 Figure (cont.) : Simulated haplotypes with 200 markers and two ploidy levels, namely autotetraploid ($m = 4$) and autohexaploid ($m = 6$). Black and white rectangles indicate two allelic variants in each marker. In all configurations, marker doses varied from zero to $\frac{m}{2}$. Each horizontal line indicates homologous chromosomes which are grouped in homology groups. We used scenario C, from simulation study 1, as the template for this second simulation study, i.e., the allelic variants were randomly assigned to the $m$ homologous chromosomes. Five different parental haplotypes were considered for each ploidy level.
