## Supplementary material for "Linkage analysis and haplotype phasing in experimental autopolyploid populations with high ploidy level using hidden Markov models": Figure S6

### Figure S6: Boxplots of the average Euclidean distances between the estimated and simulated distance vectors for simulation study 2

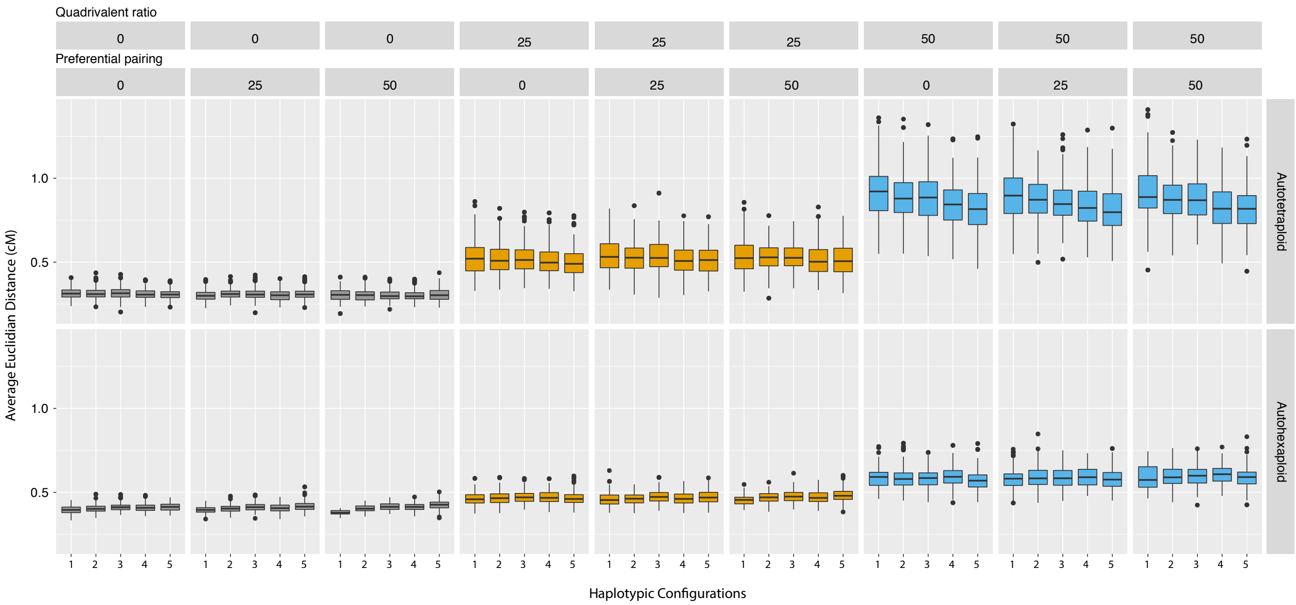


Figure S6 : Boxplots of the average Euclidean distances between the estimated and simulated distance vectors for simulation study 2. Only recombination fraction verctors from correctly estimated linkage phase configurations are considered. Each column contains a set of 5 haplotypic configurations and indicates a combination of quadrivalent formation rate and preferential pairing for autotetraploid configurations (top row) and for autohexaploid configurations (bottom row)
