## Supplementary material for "Linkage analysis and haplotype phasing in experimental autopolyploid populations with high ploidy level using hidden Markov models": Figure S7

### Figure S7: Examples autotetraploid and autohexapoloid maps estimated from datasets with three quadrivalent formation rates: $0.00$, $0.25$ and $0.50$

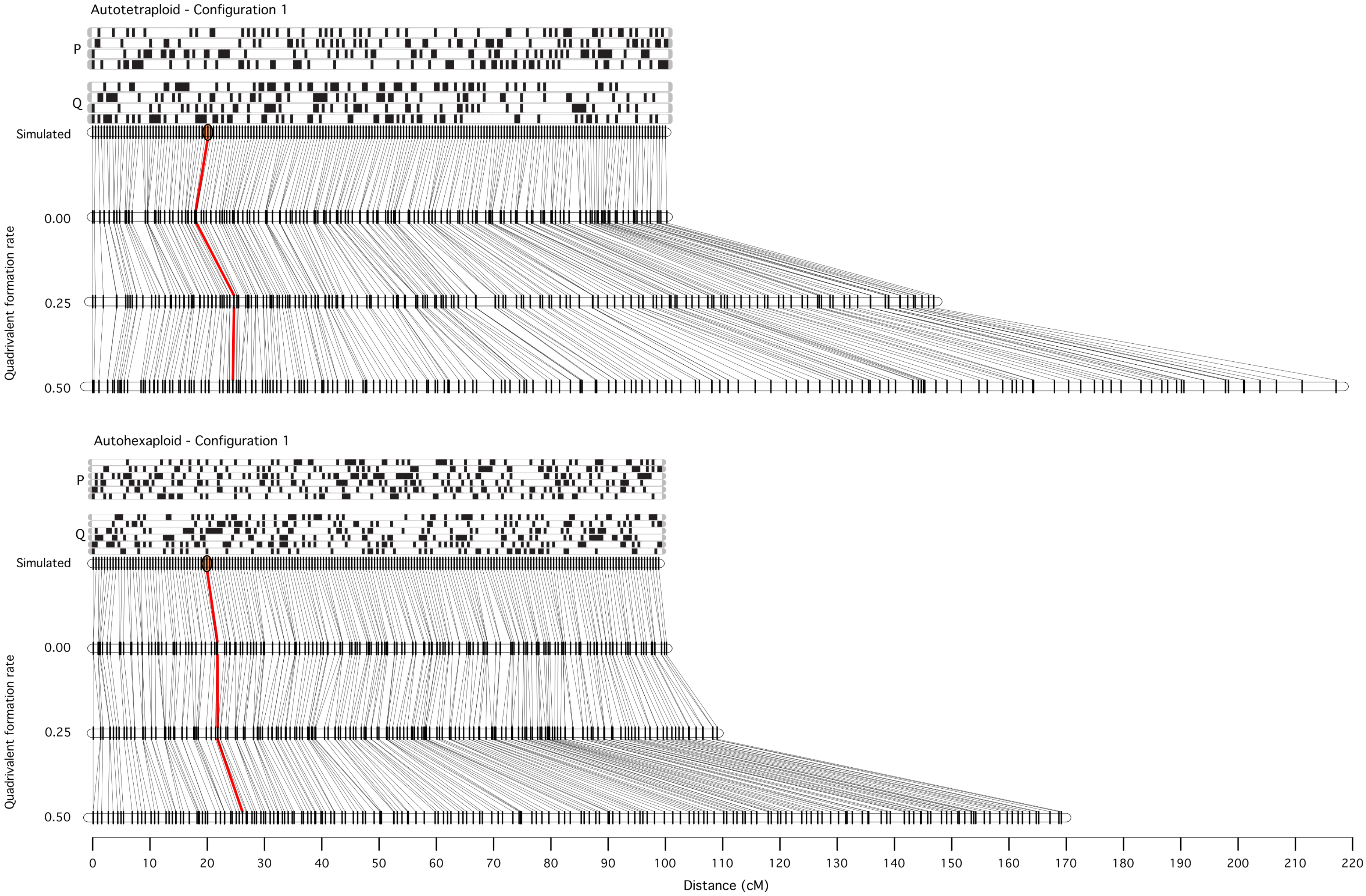


Figure S7 : Two examples of the effect of increasing quadrivalent formation rate in genetic maps selected from simulation 2. Black and white rectangles indicate two allelic variants in each marker. Each horizontal line indicates homologous chromosomes which are grouped in homology groups in parents $P$ and $Q$. Autotetraploid and autohexaploid examples are presented. Both examples show the haplotypic configuration used to simulate 200 equally spaced markers with a final length of 100 cM (configuration 1, simulation 2). The centromere was positioned at 20.0 cM from the beginning of the chromosome (orange dot). Datasets were simulated with three levels of quadrivalent formation rate (0.00, 0.25 and 0.50) and random chromosome segregation (non preferential pairing) using the software PedigreeSim [1]. Maps were estimated for each dataset and are presented here. The red line connects the marker simulated at the centromere with the position of the same marker at the estimated maps.
