## Supplementary material for "Linkage analysis and haplotype phasing in experimental autopolyploid populations with high ploidy level using hidden Markov models": File S8

### File S8: Summary of results from B2721 population map construction

Table S8.1 : Summary of the B2721 population genetic map, comprising 12 linkage groups

| Chromosome number | map length (cM) | | LOD[1] | map length (cM) | | LOD[1] | map length (cM) | | LOD[1] | Number of markers |
|---|---|---|---|---|---|---|---|---|---|---|
| | *de novo* | genomic | | *de novo* + fitTetra proportions | genomic + fitTetra proportions | | *de novo* + global error | genomic + global error | | |
| 1 | 332.3 | 262.4 | 44.3 | 293.1 | 228.2 | 46.1 | 159.3 | 149.8 | 26.5 | 369 |
| 2 | 247.5 | 176.4 | 42.5 | 227.1 | 154.2 | 45.2 | 122.8 | 103.3 | 32.5 | 229 |
| 3 | 249.5 | 173.4 | 53.8 | 224.2 | 148.0 | 57.7 | 117.6 | 101.6 | 39.5 | 277 |
| 4 | 279.8 | 186.1 | 70.3 | 247.9 | 152.3 | 76.4 | 122.9 | 95.1 | 53.0 | 376 |
| 5 | 181.9 | 185.8 | 6.6 | 148.4 | 147.0 | 7.8 | 90.7 | 91.6 | 4.8 | 247 |
| 6 | 257.8 | 168.9 | 64.2 | 237.6 | 145.5 | 67.3 | 124.5 | 93.7 | 44.7 | 375 |
| 7 | 238.0 | 162.4 | 57.8 | 203.2 | 123.6 | 63.4 | 110.1 | 91.9 | 41.2 | 367 |
| 8 | 161.1 | 124.5 | 29.9 | 140.2 | 103.6 | 32.2 | 87.0 | 80.1 | 20.0 | 242 |
| 9 | 210.0 | 159.3 | 40.0 | 174.2 | 123.7 | 43.0 | 102.4 | 94.0 | 27.8 | 264 |
| 10 | 205.0 | 178.0 | 10.9 | 184.8 | 157.5 | 12.6 | 125.0 | 112.4 | 11.7 | 204 |
| 11 | 190.7 | 158.4 | 24.3 | 162.2 | 127.7 | 26.5 | 108.8 | 97.6 | 18.2 | 237 |
| 12 | 185.2 | 157.4 | 16.5 | 151.9 | 119.1 | 20.0 | 90.3 | 88.4 | 11.7 | 187 |

1: LOD Scrore was computed using the base-10 logarithm of ratio between the multilocus likelihood of the genomic ordered map and the *de novo* ordered map, (i.e., the likelihood of the genomic ordered map was higher in all cases).

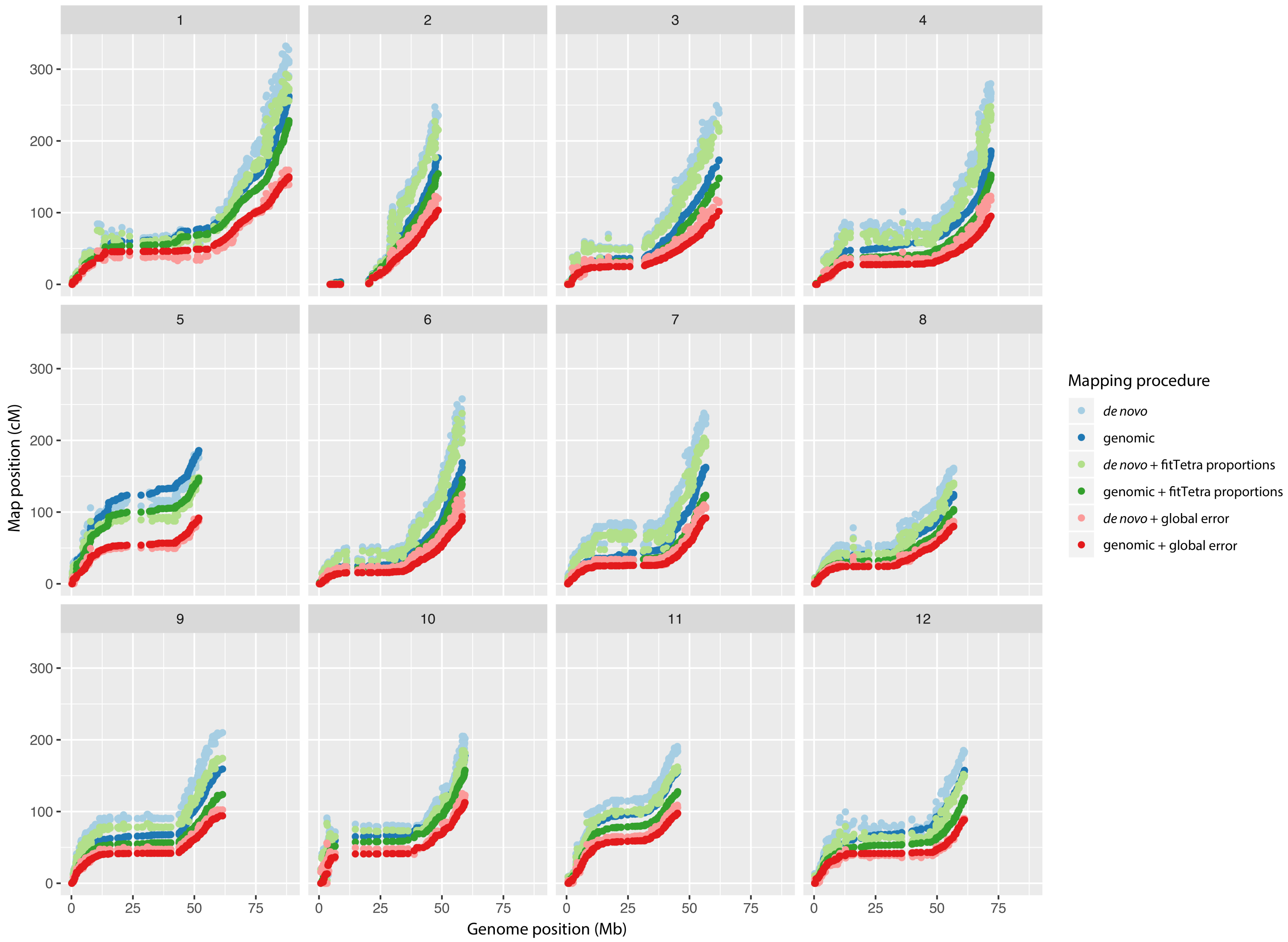


Figure S8.1 : Scatter plot of map distances versus genomic positions for 12 linkage groups in B2721 population. We used two different SNP orders: : a *de novo* order provided by MDS algorithm [1] and the order obtained from the Solanum tuberosum genome version 4.03 [2]. For each order, we applied three recombination fraction multilocus estimation procedures: $(i)$ using marker dosage, $(ii)$ using the proportions provided by the R package fitTetra [3] in the HMM model and $(iii)$ using a global error in the HMM model.
