## Supplementary material for "Linkage analysis and haplotype phasing in experimental autopolyploid populations with high ploidy level using hidden Markov models": Figure S9

### Figure S9: Simulated haplotypes for comparison between polymapR and HMM-based method

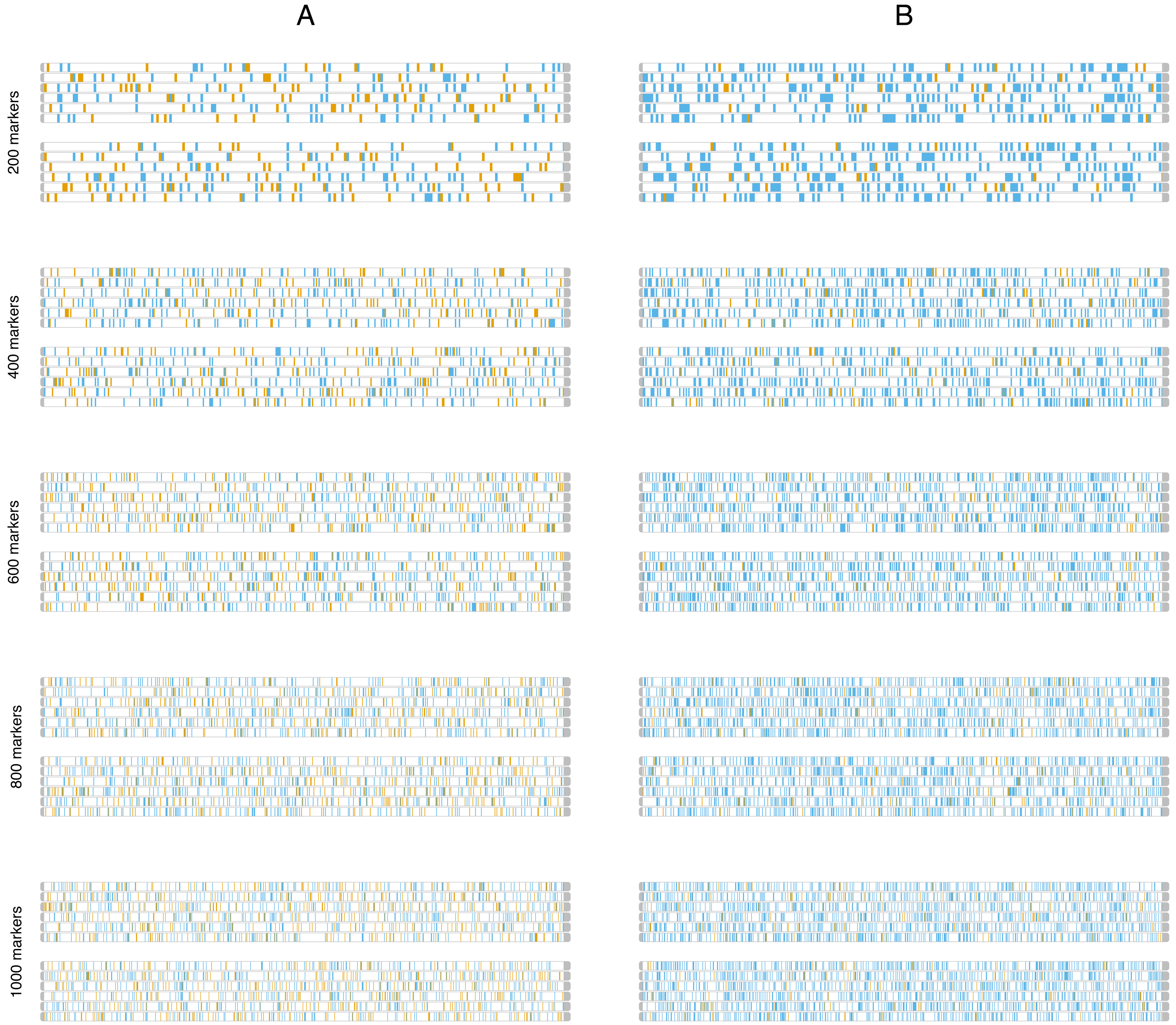


Figure S9 : Simulated haplotypes with 200, 400, 600, 800 and 1000 markers in autohexaploid ($m = 6$) parents. Orange rectangles indicate alternative allelic variants in simplex and double simplex markers; blue rectangles indicate alternative allelic variants in the remaining dosage configurations. (A) proportions of markers with 0 and 1 dose: 40%; markers with 2 and 3 doses: 10%; (B) proportions for all dosage types from 0 to 3: 25%. Proportions simulated for both parents. Each horizontal line indicates homologous chromosomes which are grouped in homology groups. Similarly to Simulation 2, the allelic variants were randomly assigned to the $m$ homologous chromosomes.
