## Supplementary material for "Linkage analysis and haplotype phasing in experimental autopolyploid populations with high ploidy level using hidden Markov models": File S10

### File S10: Results of comparison between polymapR and HMM-based method

Table S10.1 : Percentage of corrected estimated linkage phase configurations in parents P and Q across 30 simulations. Numbers in parentheses indicate the percentage of markers included in the resulting map.

| Dosage proportions | Number of markers | | LOD = 3[i] | | | | LOD = 5[i] | | | |
|---|---|---|---|---|---|---|---|---|---|---|
| | | | polymapR | | HMM-based | | polymapR | | HMM-based | |
| 40%, 40%, 10%, 10%[ii] | 200 | P | 100.0 | (90.5) | 100.0 | (100.0) | 96.7 | (81.7) | 100.0 | (99.4) |
| | | Q | 100.0 | | 96.7 | | 96.7 | | 100.0 | |
| | 400 | P | 100.0 | (93.9) | 100.0 | (99.8) | 96.7 | (81.5) | 100.0 | (99.6) |
| | | Q | 100.0 | | 100.0 | | 96.7 | | 100.0 | |
| | 600 | P | 96.7 | (95.5) | 100.0 | (100.0) | 93.3 | (86.5) | 100.0 | (100.0) |
| | | Q | 96.7 | | 96.7 | | 93.3 | | 100.0 | |
| | 800 | P | 100.0 | (95.2) | 100.0 | (100.0) | 96.7 | (86.2) | 100.0 | (100.0) |
| | | Q | 100.0 | | 100.0 | | 96.7 | | 100.0 | |
| | 1000 | P | 100.0 | (96.1) | 96.7 | (100.0) | 96.7 | (88.5) | 100.0 | (99.9) |
| | | Q | 100.0 | | 96.7 | | 96.7 | | 100.0 | |
| 25%, 25%, 25%, 25%[ii] | 200 | P | 0.0 | -[iii] | 96.7 | (99.3) | 0.0 | -[iii] | 96.7 | (97.9) |
| | | Q | 0.0 | | 100.0 | | 0.0 | | 100.0 | |
| | 400 | P | 0.0 | (24.6) | 93.3 | (99.5) | 0.0 | (20.2) | 100.0 | (98.9) |
| | | Q | 0.0 | | 90.0 | | 0.0 | | 100.0 | |
| | 600 | P | 16.7 | (59.6) | 83.3 | (99.9) | 33.3 | (46.3) | 100.0 | (99.8) |
| | | Q | 76.7 | | 86.7 | | 96.7 | | 100.0 | |
| | 800 | P | 100.0 | (69.1) | 93.3 | (99.9) | 100.0 | (51.3) | 100.0 | (99.8) |
| | | Q | 100.0 | | 100.0 | | 100.0 | | 100.0 | |
| | 1000 | P | 100.0 | (77.4) | 86.7 | (100.0) | 100.0 | (56.9) | 100.0 | (99.9) |
| | | Q | 100.0 | | 80.0 | | 100.0 | | 100.0 | |

*i*: For the HMM-based method, the LOD threshold indicates the value from which the multipoint likelihood was used to chose the best phase configuration, polymapR uses LOD thresholds to make decisions whether a marker should be used in a certain mapping context, such as, clustering homologous chromosomes or to assign it into assembled linkage groups. Thus, they are not directly comparable.
*ii*: (40%, 40%, 10%, 10%) represents higher proportion of simplex and double simplex markers, with 40% of the simulated markers being nulliplex, 40% simplex, 10% duplex and 10% triplex in both parents; (25%, 25%, 25%, 25%) represents equal proportions for all doses, with 25% for all dosage types, from nulliplex to triplex (see Figure S9)
*iii*: For all simulations, the number of single dose markers was not sufficient to identify all six homologous, thus the map was not constructed.

Table S10.2 : Mean and standard deviation (in parentheses) of the map lengths orders across the correctly phased simulations.

| Dosage proportion | Number of markers | LOD = 3 | | | | LOD = 5 | | | |
|---|---|---|---|---|---|---|---|---|---|
| | | polymapR[i] | HMM-based[ii] | | | polymapR[i] | HMM-based[ii] | | |
| | | MDS + PC | simulated | MDS | MDS + global error | MDS | simulated | MDS + PC | MDS + global error |
| 40%, 40%, 10%, 10%[iii] | 200 | 88.3(4.8) | 97.4(3.1) | 143.9(6.7) | 94.7(3.7) | 88.6(4.5) | 97.3(3.1) | 143(7.1) | 94.5(3.7) |
| | 400 | 88.7(3.6) | 98.6(2.6) | 212.7(10.2) | 97.2(3.0) | 88.6(3.6) | 98.6(2.6) | 212.1(10.3) | 97.3(3.0) |
| | 600 | 87.5(2.8) | 99.1(2.7) | 277.3(11.2) | 100(3.0) | 87.5(3.0) | 99.1(2.7) | 277.5(11.2) | 100.2(3.0) |
| | 800 | 87.2(2.9) | 99.5(2.5) | 350.3(17.3) | 100.6(3.0) | 87.3(3.0) | 99.4(2.4) | 350.3(17.3) | 100.6(2.9) |
| | 1000 | 86.6(3.2) | 99.3(2.9) | 411.1(19) | 102.8(3.3) | 91.4(17.6) | 99.6(3.3) | 410.9(18.6) | 103(3.4) |
| 25%, 25%, 25%, 25%[iii] | 200 | -[iv] | 98.7(3.1) | 173.1(9.0) | 96.5(3.9) | -[iv] | 98.7(3.1) | 170.7(8.8) | 96.3(4.0) |
| | 400 | -[iv] | 99.1(2.6) | 256(13.9) | 97.8(2.9) | -[iv] | 99.2(2.5) | 254.9(14.3) | 97.9(2.7) |
| | 600 | 77.7(2.7) | 99.6(2.7) | 333.7(13.6) | 97.8(2.9) | 78.4(2.8) | 99.4(2.7) | 334.8(15.3) | 99.5(3.1) |
| | 800 | 78.7(2.5) | 99.8(2.4) | 419.9(19.6) | 101.6(3.1) | 78.5(2.4) | 99.5(2.4) | 420.2(19.5) | 101.6(3.0) |
| | 1000 | 78.7(3.7) | 99.7(3.3) | 508.1(22.4) | 103.6(3.4) | 78.8(3.5) | 99.6(3.2) | 506.4(20.6) | 103.5(3.4) |

*i*: For polymapR, the marker positions were estimated using the projection of the MDS result onto a single dimension principal curve (PC).
*ii*: For HMM-based method, we present the map length given the simulated and the MDS order. For the MDS order, we present the results of the map re-estimation using a 5% global error.
*iii*: (40%, 40%, 10%, 10%) represents higher proportion of simplex and double simplex markers, with 40% of the simulated markers being nulliplex, 40% simplex, 10% duplex and 10% triplex in both parents; (25%, 25%, 25%, 25%) represents equal proportions for all doses, with 25% for all dosage types, from nulliplex to triplex (see Figure S9)
*iv*: None of the maps were correctly phased.

Table S10.3 : Average of the elapsed time (in muinutes) across 30 simulations.

| Dosage proportions | Number of markers | LOD = 3 | | LOD = 5 | |
|---|---|---|---|---|---|
| | | polymapR | HMM-based | polymapR | HMM-based |
| 40%, 40%, 10%, 10% | 200 | 0.7 | 6.2 | 0.6 | 10.2 |
| | 400 | 2.4 | 15.1 | 2.3 | 17.7 |
| | 600 | 4.8 | 28.6 | 4.6 | 33.0 |
| | 800 | 8.7 | 47.5 | 8.1 | 53.2 |
| | 1000 | 12.7 | 72.7 | 11.9 | 78.4 |
| 25%, 25%, 25%, 25% | 200 | - | 8.2 | - | 16.1 |
| | 400 | 4.4 | 21.7 | 2.5 | 26.5 |
| | 600 | 12.9 | 34.3 | 9.2 | 38.3 |
| | 800 | 24.9 | 58.2 | 17.7 | 62.4 |
| | 1000 | 39.4 | 86.8 | 29.0 | 91.2 |

Each map was constructed in a single Intel(R) Xeon(R) CPU E5-2670 0 @ 2.60GHz, 128GB RAM.

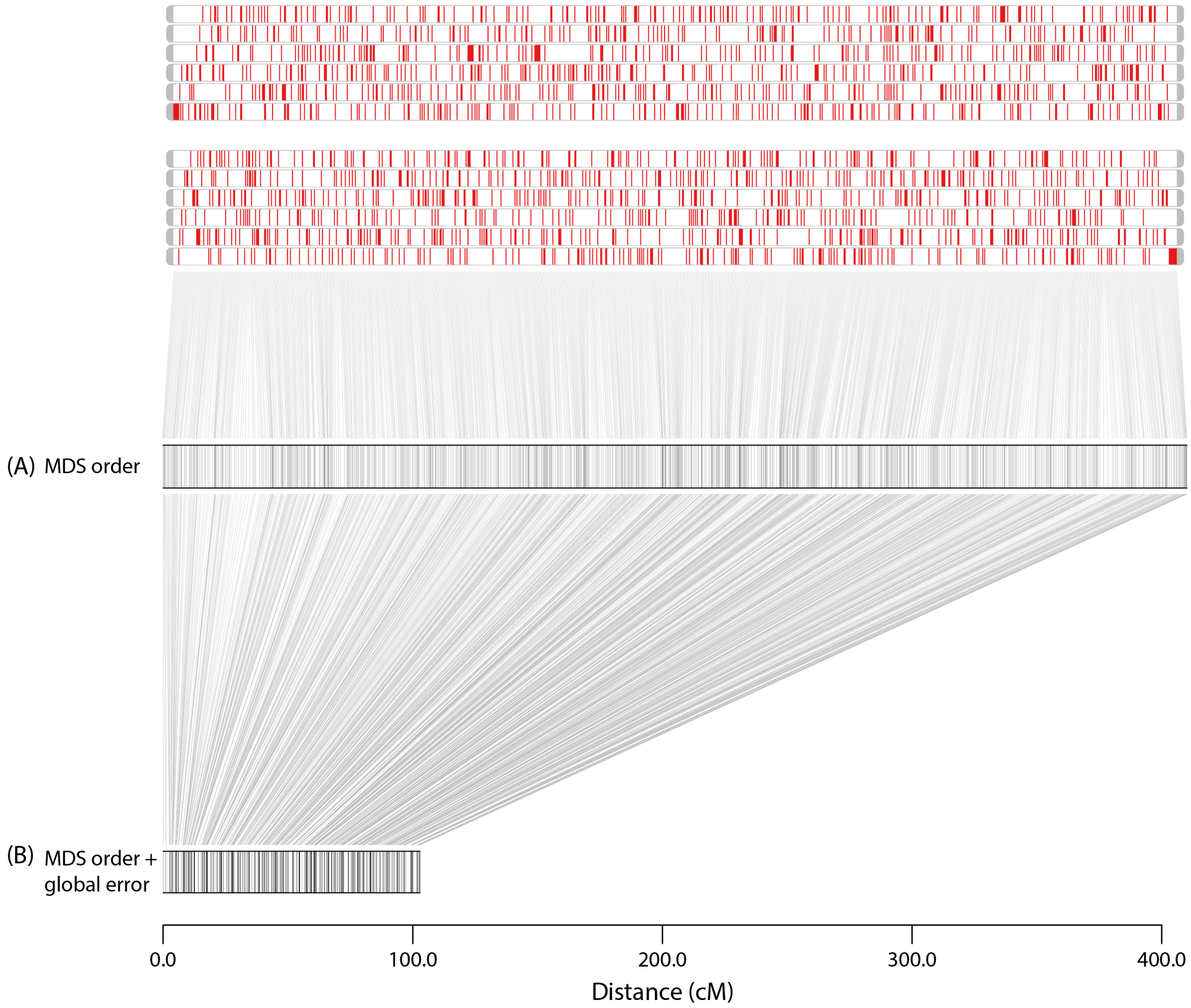


Figure S10.1 : Example of MDS ordered map estimated using marker dosages with no error modeling (A) and modeling a global error of 5% in the HMM emission function (B).
